## Supplementary Information for "Tigerfish designs oligonucleotide-based *in situ* hybridization probes targeting intervals of highly repetitive DNA at the scale of genomes"

### Supplementary Fig. 1

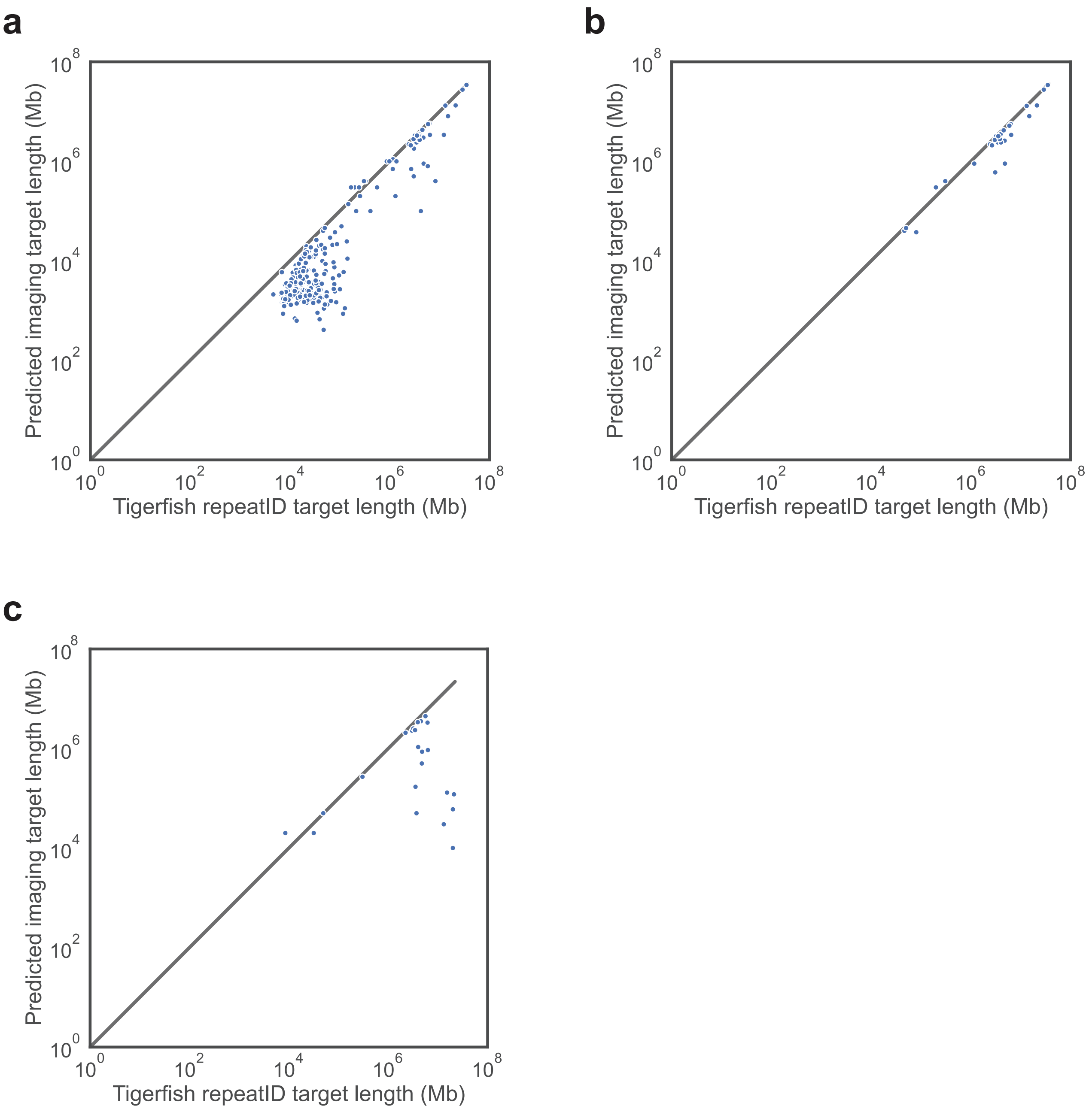


**Supplementary Fig. 1 | Effective lengths of Tigerfish target intervals. a**, Scatter plot depicting the effective lengths of intervals identified and processed successfully for probe design using permissive parameters (Y-axis) and the length of these intervals when first identified at the “Repeat discovery” step. **b**, Scatter plot depicting the effective lengths of intervals identified and processed successfully for probe design using conservative parameters (Y-axis) and the length of these intervals when first identified at the “Repeat discovery” step. **c**, Scatter plot depicting the effective lengths of intervals inputted for probe design (Y-axis) and the length of these intervals when first identified at the “Repeat discovery” step for the 24-target panel used for in situ validation experiments.

#### Supplementary Fig. 2

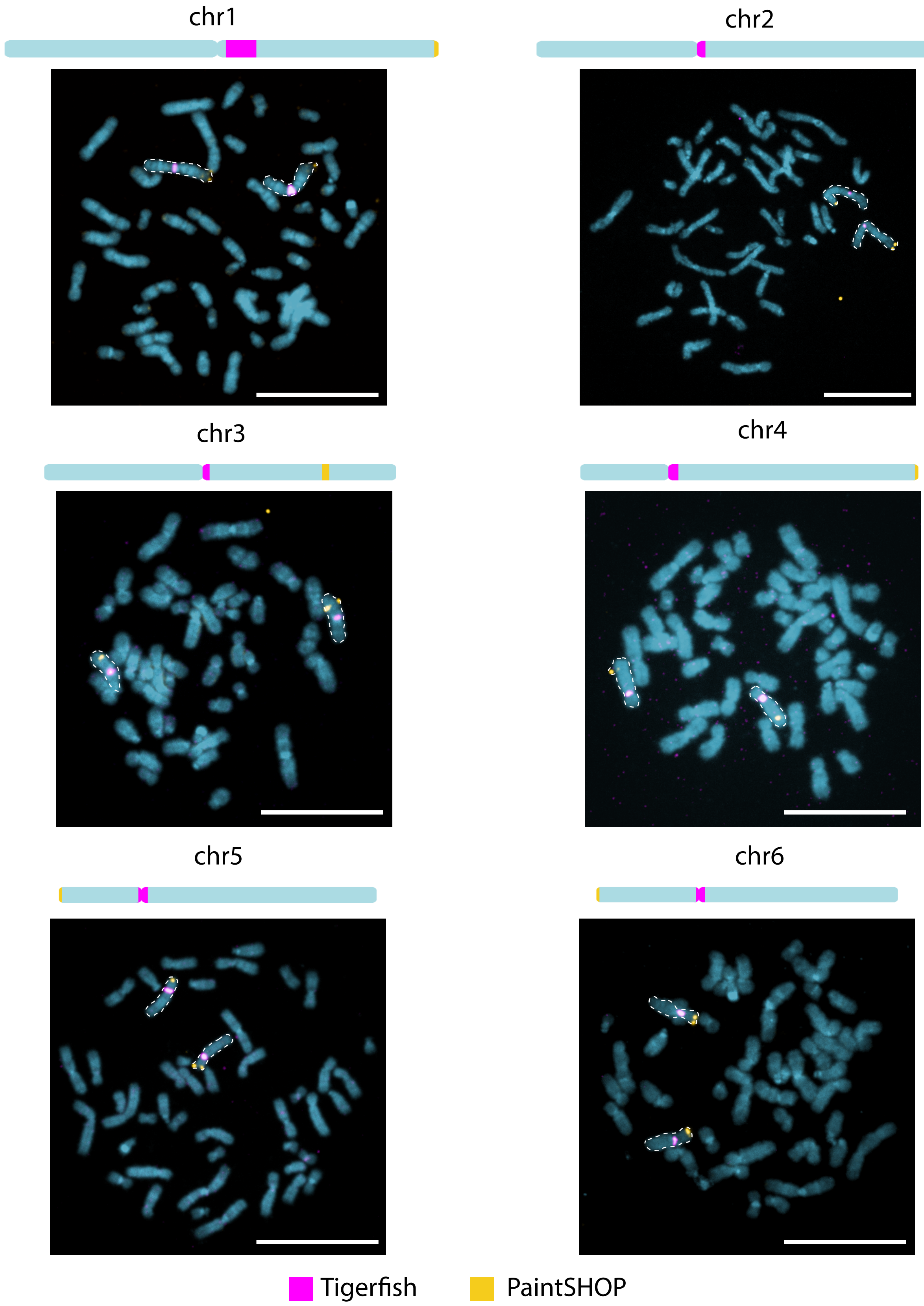


**Supplementary Fig. 2 | Full-field metaphase spreads for chromosomes 1–6.** Full-field images of the metaphase spreads from which the crops depicted in Figure 4b originated showing the staining pattern of the indicated Tigerfish (magenta) and PaintSHOP (yellow) probe sets. Images are maximum intensity projections in Z. Scale bars, 20 μm.

#### Supplementary Fig. 3

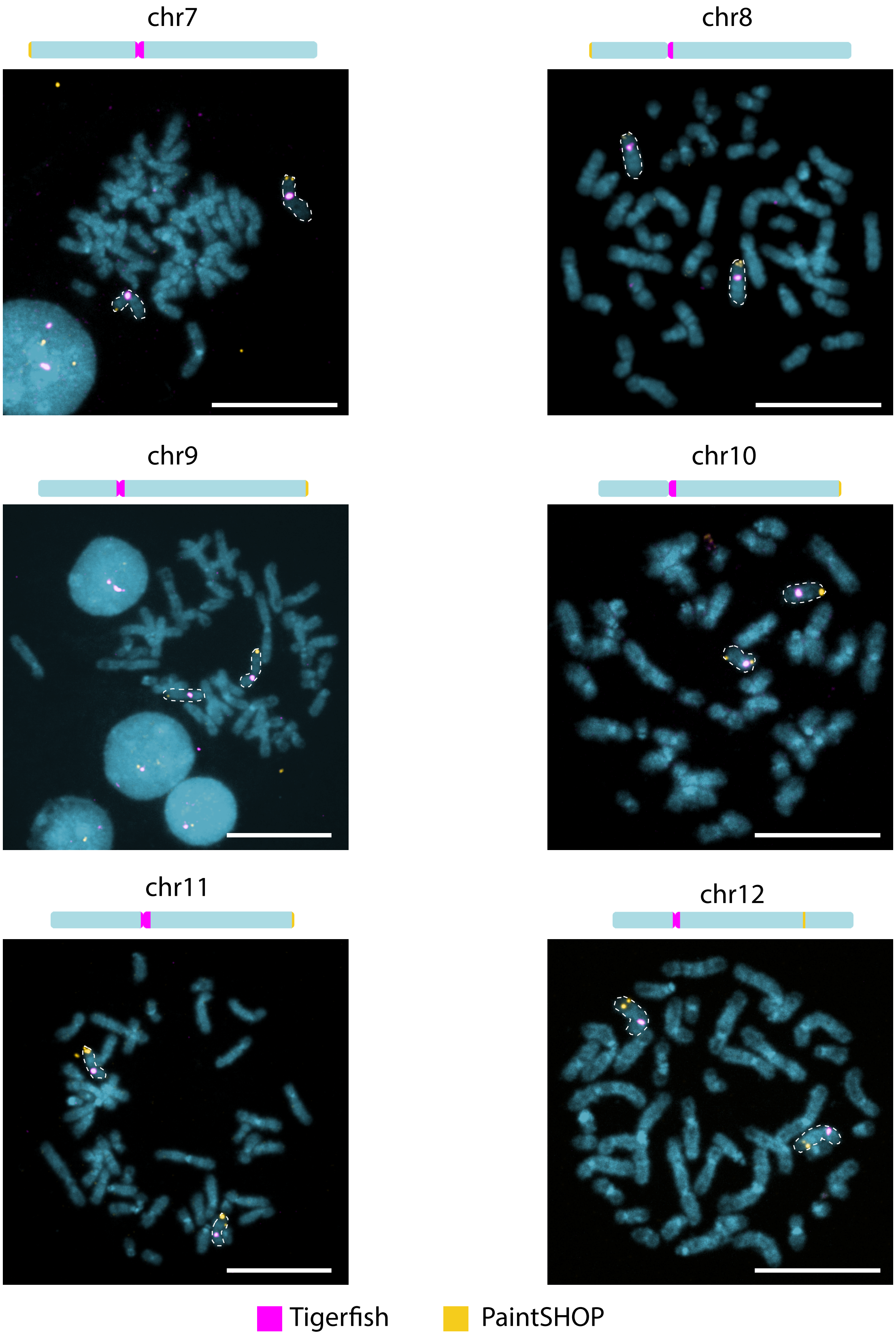


**Supplementary Fig. 3 | Full-field metaphase spreads for chromosomes 7–12.** Full-field images of the metaphase spreads from which the crops depicted in Figure 4b originated showing the staining pattern of the indicated Tigerfish (magenta) and PaintSHOP (yellow) probe sets. Images are maximum intensity projections in Z. Scale bars, 20 μm.

#### Supplementary Fig. 4

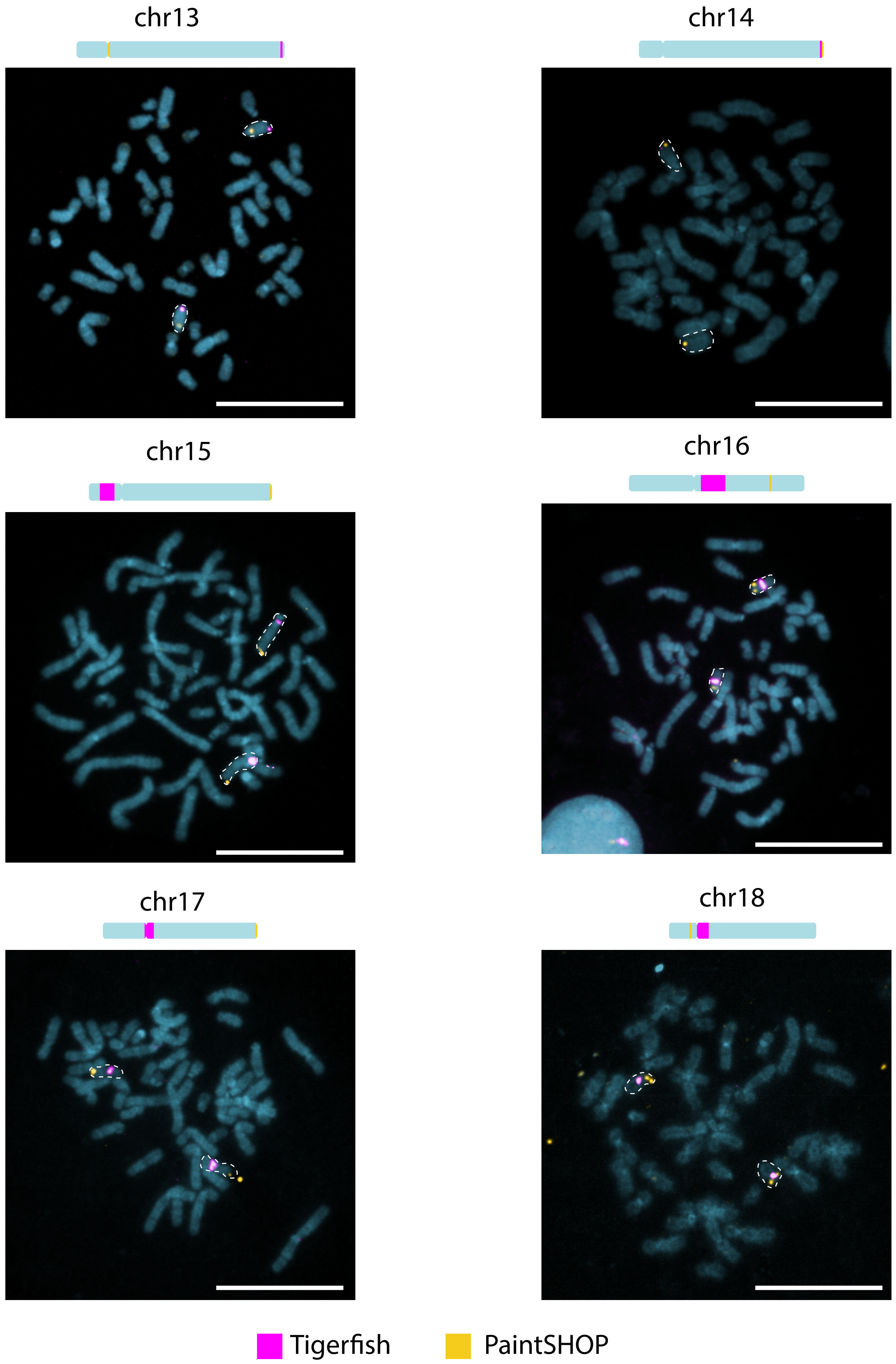


**Supplementary Fig. 4 | Full-field metaphase spreads for chromosomes 13–18.** Full-field images of the metaphase spreads from which the crops depicted in Figure 4b originated showing the staining pattern of the indicated Tigerfish (magenta) and PaintSHOP (yellow) probe sets. Images are maximum intensity projections in Z. Scale bars, 20 μm.

#### Supplementary Fig. 5

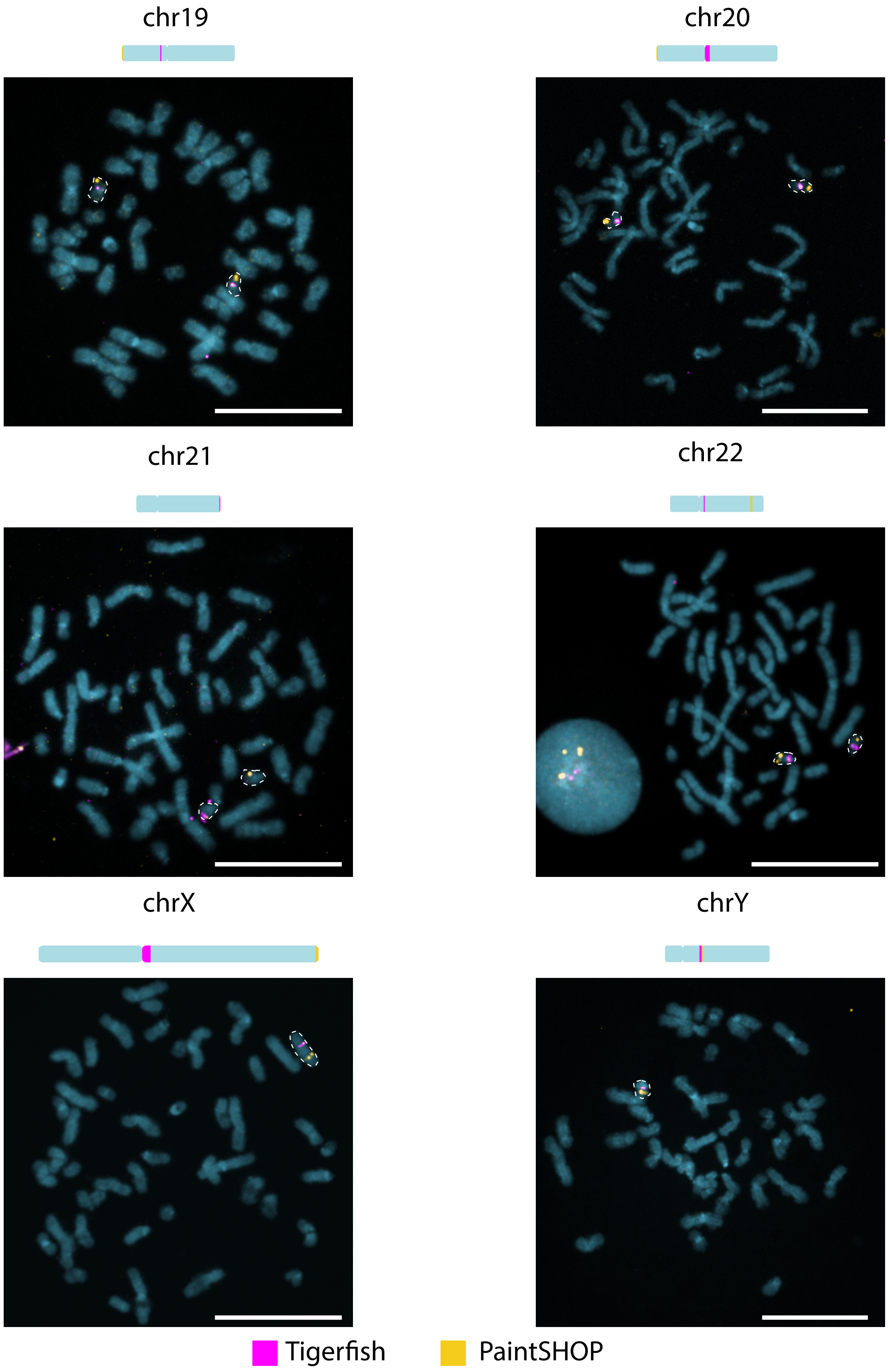


**Supplementary Fig. 5 | Full-field metaphase spreads for chromosomes 19–Y.** Full-field images of the metaphase spreads from which the crops depicted in Figure 4b originated showing the staining pattern of the indicated Tigerfish (magenta) and PaintSHOP (yellow) probe sets. Images are maximum intensity projections in Z. Scale bars, 20 μm.

#### Supplementary Fig. 6

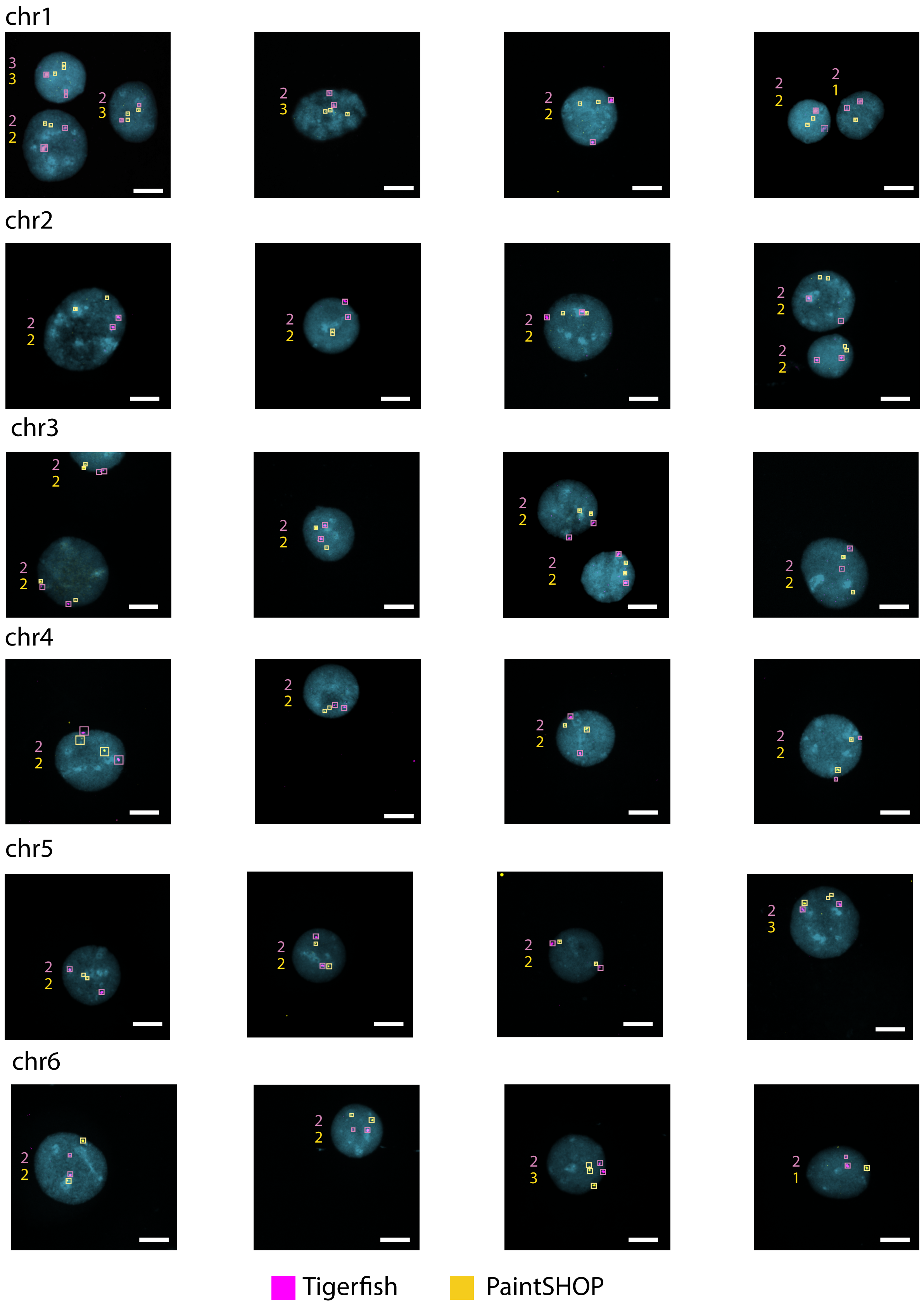


**Supplementary Fig. 6 | Representative enumeration images for chr1–6.** Four representative images of interphase nuclei and corresponding puncta counts for the specified Tigerfish (magenta) and PaintSHOP (yellow) probe sets. Images are maximum intensity projections in Z. Scale bars, 10 μm.

### Supplementary Fig. 7

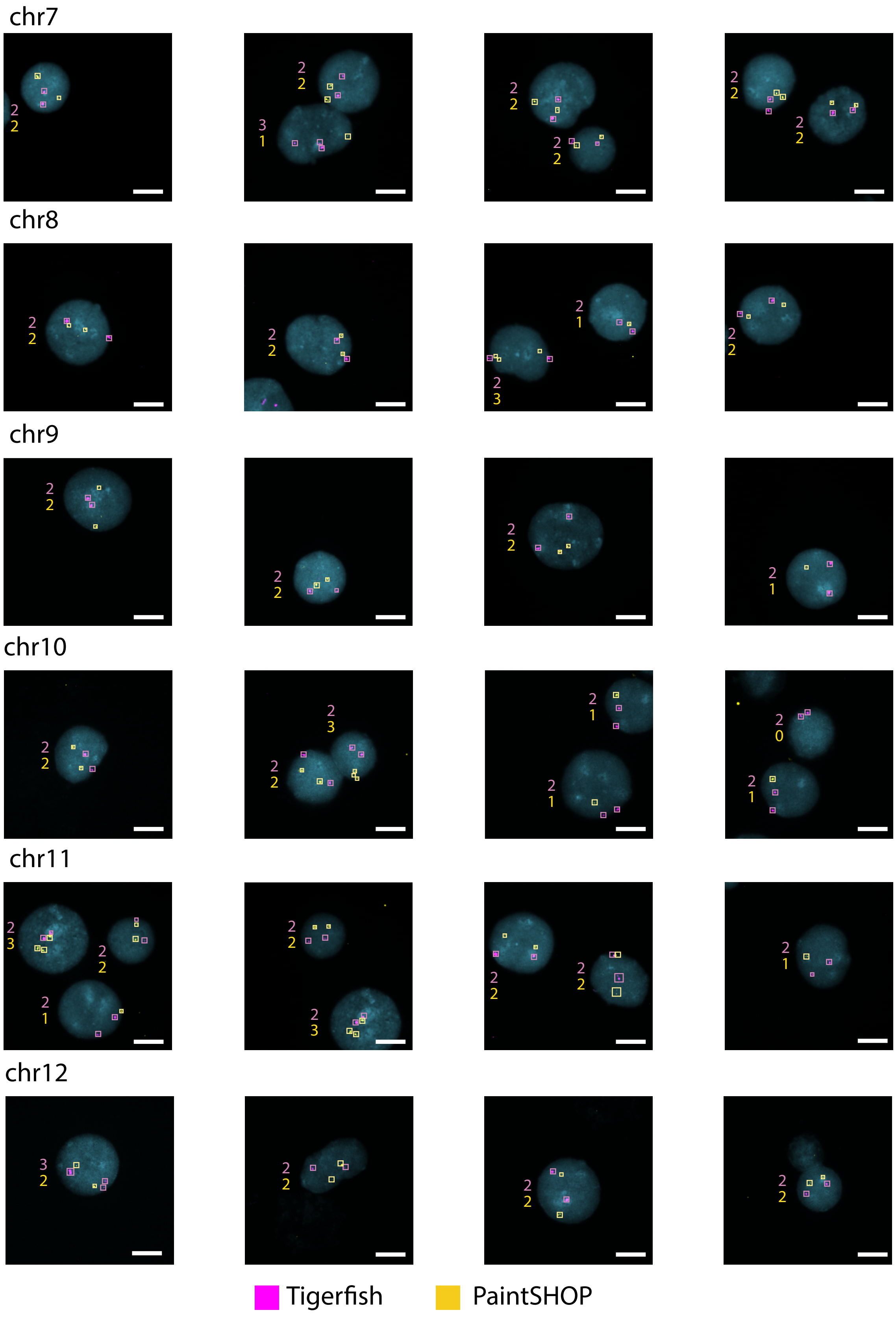


**Supplementary Fig. 8 | Representative enumeration images for chr7–12.** Four representative images of interphase nuclei and corresponding puncta counts for the specified Tigerfish (magenta) and PaintSHOP (yellow) probe sets. Images are maximum intensity projections in Z. Scale bars, 10 μm.

### Supplementary Fig. 8

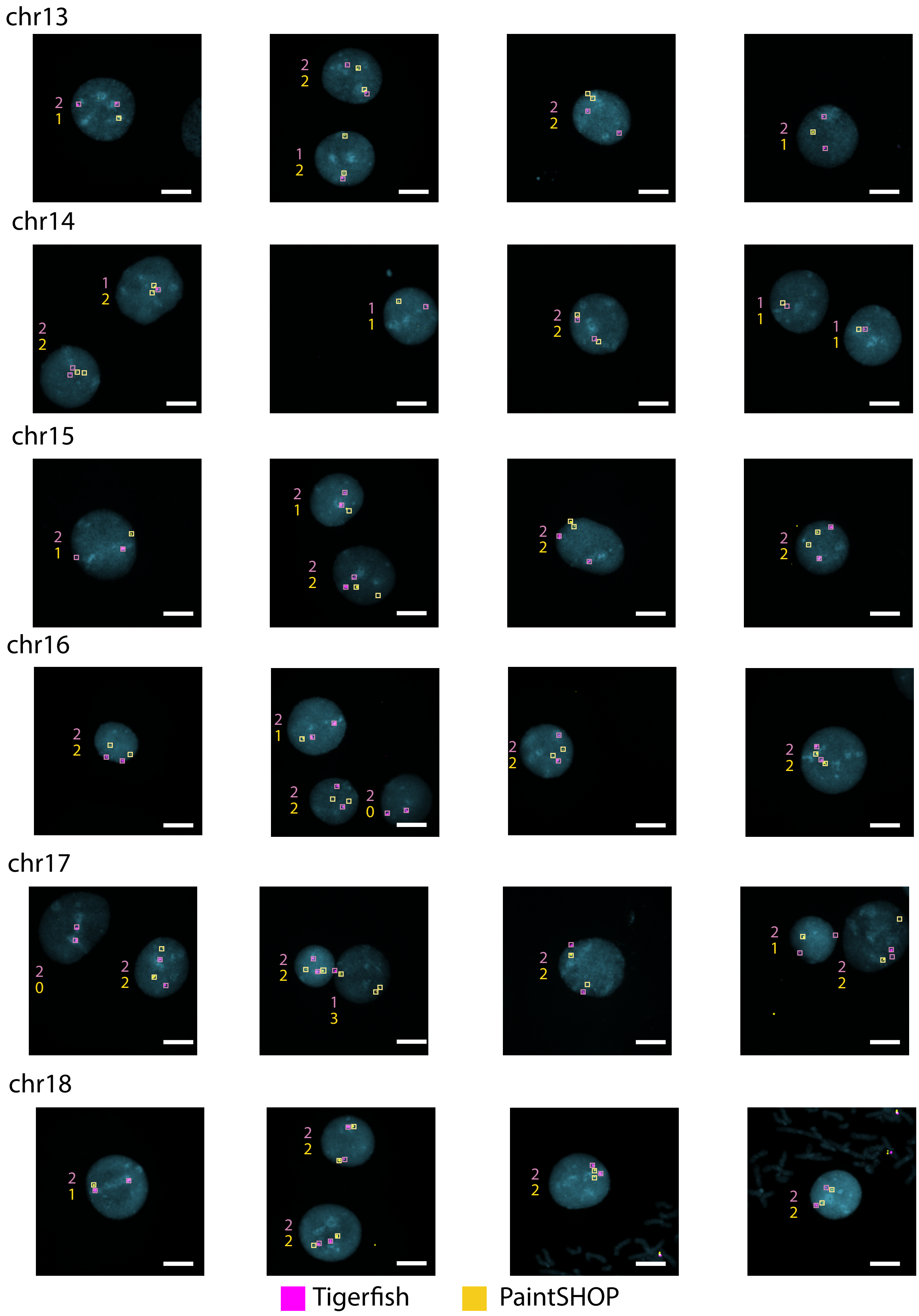


**Supplementary Fig. 8 | Representative enumeration images for chr13–18.** Four representative images of interphase nuclei and corresponding puncta counts for the specified Tigerfish (magenta) and PaintSHOP (yellow) probe sets. Images are maximum intensity projections in Z. Scale bars, 10 μm.

### Supplementary Fig. 9

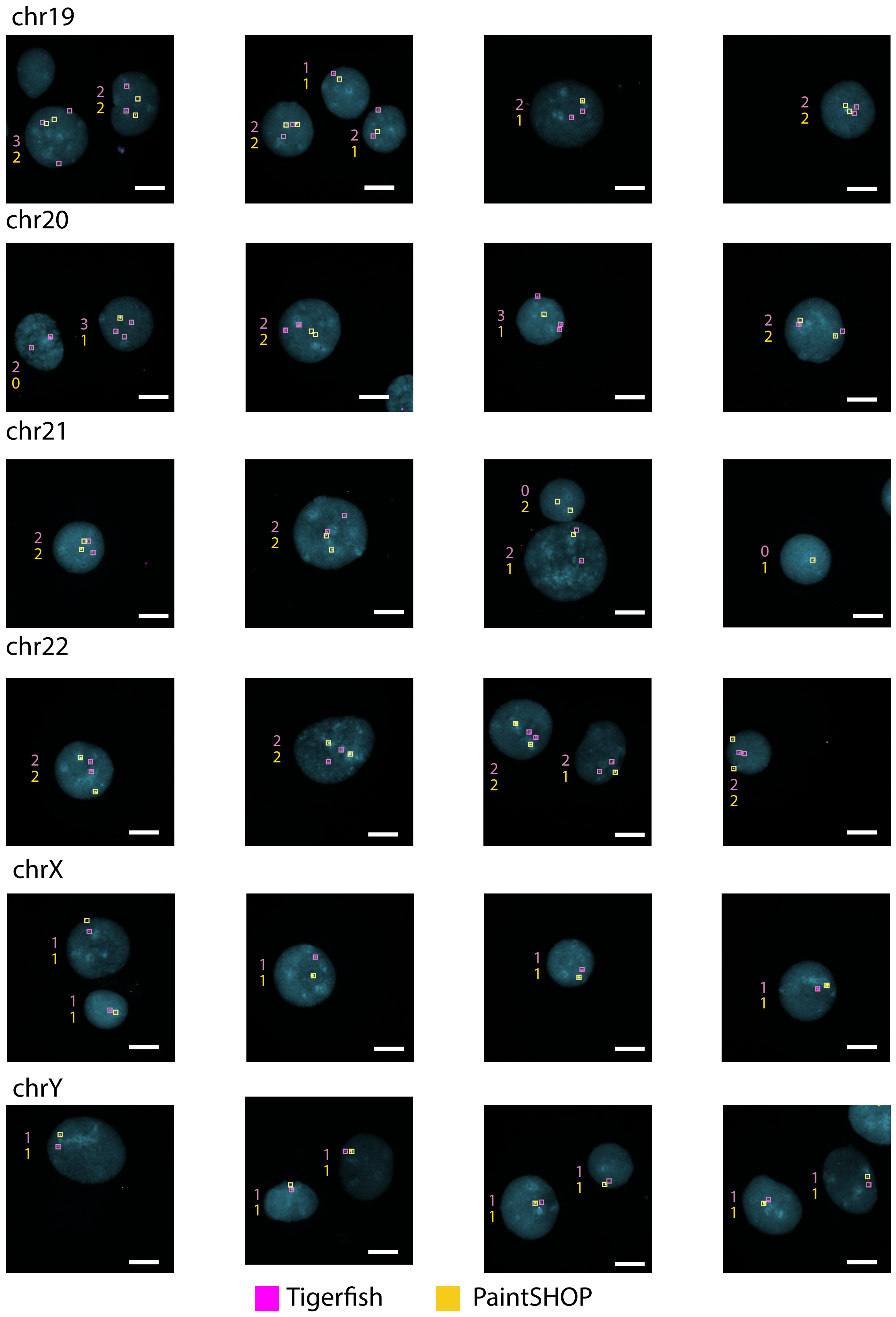


**Supplementary Fig. 9 | Representative enumeration images for chr19–Y.** Four representative images of interphase nuclei and corresponding puncta counts for the specified Tigerfish (magenta) and PaintSHOP (yellow) probe sets. Images are maximum intensity projections in Z. Scale bars, 10 μm.

#### Supplementary Fig. 10

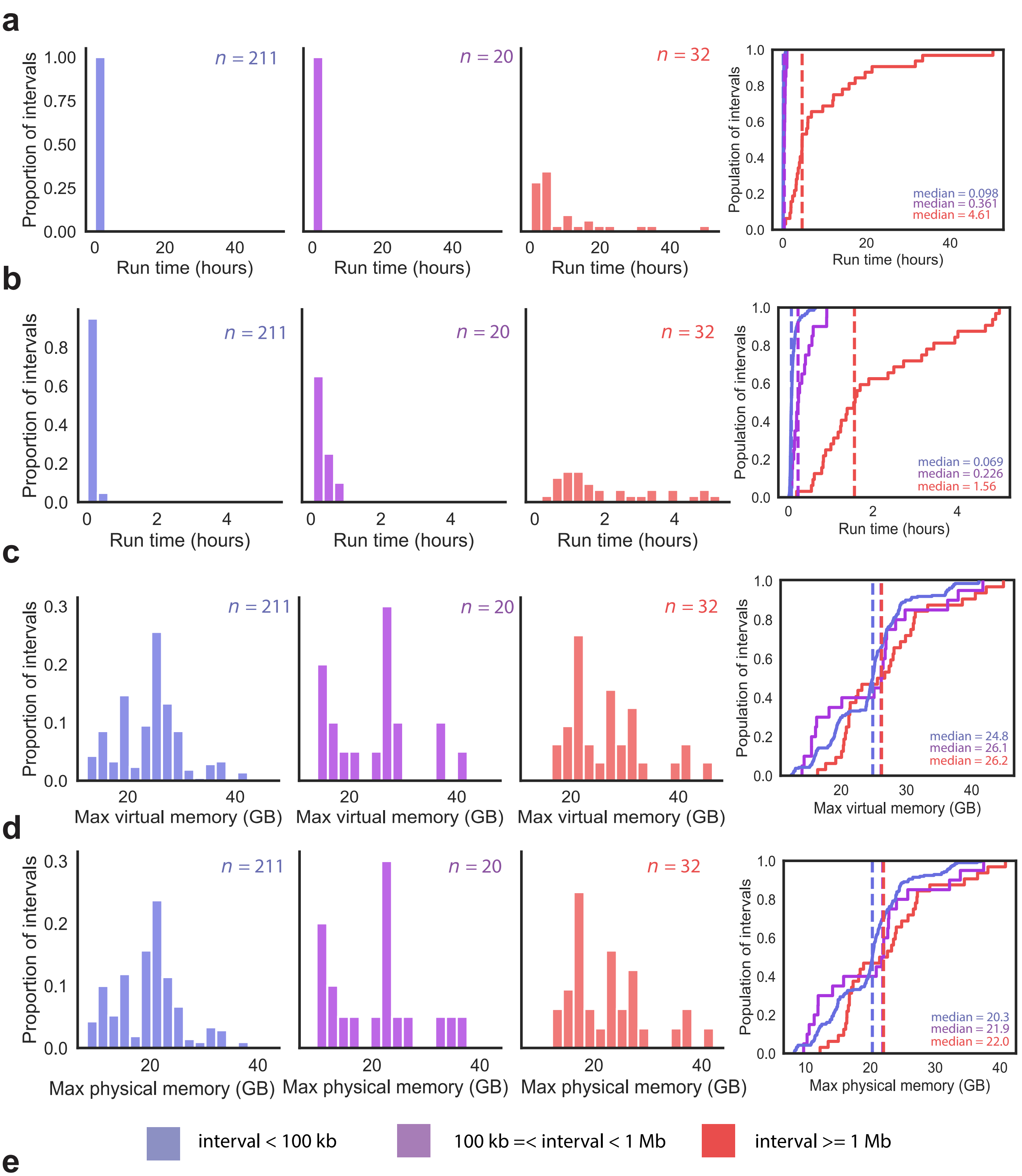


**e**

| Interval length | Min runtime | Max runtime | Min CPU time | Max CPU time | Min virtual memory | Max virtual memory | Min physical memory | Max physical memory |
|---|---|---|---|---|---|---|---|---|
| i < 100 kb | 0.03 | 0.73 | 0.01 | 0.64 | 12.35 | 41.14 | 8.25 | 37.06 |
| 1 Mb > i >= 100 kb | 0.10 | 0.99 | 0.06 | 0.91 | 14.00 | 41.71 | 9.70 | 37.47 |
| i >= 1 Mb | 0.27 | 50.18 | 0.19 | 4.99 | 16.39 | 44.87 | 12.24 | 40.84 |

**Supplementary Fig. 10 | Tigerfish computational requirements by repeat region length. a**, Histograms (left) and empirical cumulative distributions (right) of the wall-clock runtime recorded for running the 263 conservative and permissive intervals stratified by the target interval length. **b**, Histograms (left) and empirical cumulative distributions (right) of the CPU runtime recorded for running the 263 conservative and permissive intervals stratified by the target interval length. **c**, Histograms (left) and empirical cumulative distributions (right) of the maximum recorded virtual memory allocation for running the 263 conservative and permissive intervals. **d**, Histograms (left) and empirical cumulative distributions (right) of the maximum recorded physical memory allocation for running the 263 conservative and permissive intervals. Vertical dashed lines in the cumulative distribution plots correspond to the median values. **e**, Summary statistics for the values presented in panels a–d.

#### Supplementary Note
*Tigerfish inputs and outputs*

| Input(s) | Snakemake step | Output(s) | Option | Usage |
|---|---|---|---|---|
| FASTA | **generate_jf_count*** Generates a Jellyfish index file to count *k*-mers genome wide. | counts.jf | mer_val<br>fasta_file | *Required*. *k*-mer size<br>*Required*. Genomic reference FASTA |
| FASTA | **generate_bt2_indices*** Generates genome wide Bowtie2 (Bt2) index to align probes. | Bt2 Indices | fasta_file | *Required*. Genomic reference FASTA |
| counts.jf<br>FASTA | **generate_jf_idx*** Generates *k*-mer count index files for each scaffold. | counts.txt<br>index.txt<br>scaffold FASTA | fasta_file<br>mer_val<br>sample | *Required*. Genomic reference FASTA<br>*Required*. *k*-mer size<br>*Required*. Scaffold name(s) |
| BED | **split_bed**** Takes BED file and splits coordinates into distinct files. | BED<br>BED<br>... | bed_file<br>sample | *Optional*. BED file with repeat target coordinates.<br>*Required*. Scaffold name(s) |
| counts.txt<br>index.txt | **repeat_ID**** Identifies repeat regions over queried chromosomes. | BED | sample<br>file start<br>window<br>threshold<br>composition<br>mer_val | *Required*. Scaffold name(s)<br>*Required*. Base position where repeat search begins.<br>*Required*. Size of *k*-mer search window (W).<br>*Required*. Min *k*-mer count value (T).<br>*Required*. Proportion of elevated *k*-mers within W (C).<br>*Required*. *k*-mer size |
| BED (repeat_ID)<br>or<br>BED (split_bed) | **design_probes**** Designs probes from identified repeat coords or user provided BED file. | probes.tsv<br>region FASTA | fasta_file<br>sample<br>min_len<br>max_len<br>min_temp<br>max_temp | *Required*. Genomic reference FASTA<br>*Required*. Scaffold name(s)<br>*Required*. Min size of candidate probe (bp)<br>*Required*. Max size of candidate probe (bp)<br>*Required*. Min melting temp of probe (C)<br>*Required*. Max melting temp of probe (C) |
| probes.tsv<br>counts.txt<br>region FASTA | **kmer_filter** Computes each probe's on-target and off-target *k*-mer counts, then sorts probes by on-target count. | k_mer_sort.tsv | c1_val<br>c2_val<br>mer_val | *Required*. Constant to rank probes by copy_num.<br>*Required*. Constant to rank probes by enrich_score<br>*Required*. *k*-mer size |
| k_mer_sort.tsv | **probe_mer_filter** Filters probes based on probe *k*-mer similarity and target binding. | k_mer_filter.tsv | enrich_score<br>copy_num<br>mer_cutoff<br>mer_val | *Required*. Min proportion a probe's *k*-mers must bind within a repeat region target.<br>*Required*. Total sum of any probe's *k*-mers within a target repeat region.<br>*Required.* Any probes within a target repeat exceeding this proportion of shared *k*-mers will be filtered<br>*Required*. *k*-mer size |
| chrom.sizes | **generate_genome_bins:** Takes reference genome and creates bins. | alignment_bin.BED<br>threshold_bin.BED | genome_windows<br>chrom_sizes_file<br>thresh_window | *Required*. Size of genome bins for alignment.<br>*Required*. chrom.sizes file of queried genome.<br>*Required*. Size of threshold bins to flag to determine imaging target coordinates. |

** Denotes optional step if proper file paths/options are specified in config.yml*
*** Denotes step specific to either probe_design or repeat_ID run modes*

| Input(s) | Snakemake step | Output(s) | Option | Usage |
|---|---|---|---|---|
| k_mer_filter.tsv | **make_chrom_dir**<br>Seperates probe files by repeat region. | repeat.txt | | |
| repeat.txt<br>Bt2 indices<br>alignment_bin.BED | **alignment_filter**<br>Filters candidate probes using Bt2 and NUPACK to predict in silico probe binding. | filtered_probes.tsv | target_sum<br>bt2_alignments<br>seed_length<br>model_temp<br>max_pdups<br><br>min_on_target<br>max_probe_return<br>off_bin_thresh<br><br>align_thresh<br><br>ref_flag | *Required*. Total on-target sum of all probes desired.<br>*Required*. Bt2will return alignments up to this val.<br>*Required*. Controls Bt2 seed length.<br>*Required*. Temperature of NUPACK predict model.<br>*Required*. Probes in final dataset with predicted binding greater than this value will be filtered.<br>*Required*. Min on target score for any given probe.<br>*Required*. Max probes to be returned/repeat.<br>*Required*. Off-target threshold over any non-target genomic bin.<br>Required. Min binding sites required to flag a binned region as significant toward probe binding.<br>Required. Provides intermediate output files. |
| filtered_probes.tsv | **merge_alignment_filter**<br>Aggregates all candidate probes by scaffold. | chrom_probes.tsv | | |
| chrom_probes.tsv | **split_rm_alignments**<br>Creates a new directory containing all candidate probe files by repeat. | scaffold/region.tsv | | |
| scaffold/region.tsv | **align_probes**<br>Takes candidate probes corresponding to compute target specificity. SAM derived sequences are used to compute *in silico* binding. | region_align.txt | bt2_alignments<br>seed_length<br>model_temp<br><br>mer_val | *Required*. Bt2 will return alignments up to this val.<br>*Required*. Controls Bt2 seed length.<br>*Required*. Temperature of NUPACK predict model. binding greater than this value will be filtered.<br>*Required.* Any probes within a target repeat exceeding this proportion of shared *k*-mers will be filtered |
| region_align.txt | **derived_beds**<br>Creates BED file from SAM derived sequence alignment for each candidate probe. | derived_align.BED | | |
| region_align.txt | **get_region_bed**<br>Creates BED file for the target repeat region. | repeat.BED | | |
| derived_align.BED<br>repeat.BED | **bedtools_intersect**<br>Performs two bedtools intersects:<br>1. Derived alignments against threshold genome bins.<br>2. Repeat region against threshold genome bins. | derived_BEDtools.txt<br>repeat_BEDtools.txt | | |

** Denotes optional step if proper file paths/options are specified in config.yml*
*** Denotes step specific to either probe_design or repeat_ID run modes*

| Input | Snakemake step | Output | Option | Usage |
| --- | --- | --- | --- | --- |
| derived_BEDtools.txt<br>repeat_BEDtools.txt<br>region_align.txt<br>chrom.sizes | **get_alignments**<br>Computes genome wide binding summaries for all probes within a target repeat region. | binding_map.png<br>thresh_summary.txt<br>binding_quant.txt | align_thresh | *Required*. Binding sites required to flag a bin as significant toward probe signal. |
| filtered_probes.tsv<br>binding_map.png<br>thresh_summary.txt<br>binding_quant.txt | **map_region_coords**<br>Adds imaging target coordinates to the candidate probes file. Creates probe binding summary. | final_probes.tsv | | |
| final_probes.tsv<br>probe_summary.txt | **merge_mapping**<br>Aggregates all repeat regions into a single file by scaffold. | probes_merged.tsv | | |
| probes_merged.tsv | **summary**<br>Summarizes probe binding and count by repeat region target. | probe_summary.txt | | |

** Denotes optional step if proper file paths/options are specified in config.yml*
*** Denotes step specific to either probe_design or repeat_ID run modes*

| Input | Snakemake step | Output | Option | Usage |
| --- | --- | --- | --- | --- |
| filtered_probes.tsv | **gather_repeat_regions**<br>Takes filtered candidate probes and splits them by scaffold. Input file should have one repeat per scaffold. | split_probes.txt | sample | *Required*. Scaffold name(s) |
| split_probes.txt | **align_cand_probes**<br>Takes candidate probes corresponding to compute target specificity. SAM derived sequences are used to compute *in silico* binding. | region_align.txt | bt2_alignments<br>seed_length<br>model_temp<br><br>mer_val | *Required*. Bt2 will return alignments up to this val.<br>*Required*. Controls Bt2 seed length.<br>*Required*. Temperature of NUPACK predict model. binding greater than this value will be filtered.<br>*Required.* Any probes within a target repeat exceeding this proportion of shared *k*-mers will be filtered |
| region_align.txt | **derived_cand_beds**<br>Creates BED file from SAM derived sequence alignment for each candidate probe. | derived_align.BED | | |
| region_align.txt | **get_cand_region_bed**<br>Creates BED file for the target repeat region. | repeat.BED | | |
| derived_align.BED<br>repeat.BED | **bedtools_cand_intersect**<br>Performs two bedtools intersects:<br>1. Derived alignments against threshold genome bins.<br>2. Repeat region against threshold genome bins. | derived_BEDtools.txt<br>repeat_BEDtools.txt | | |
| derived_BEDtools.txt<br>repeat_BEDtools.txt<br>region_align.txt<br>chrom.sizes | **get_cand_alignments**<br>Computes genome wide binding summaries for all probes within a target repeat region. | binding_map.png<br>thresh_summary.txt<br>binding_quant.txt | align_thresh | *Required*. Binding sites required to flag a bin as significant toward probe signal. |
| chrom.sizes<br>repeat.BED | **generate_cand_chromomap***<br>Creates a karyoplot using chromoMap of the repeat target. | chromomap.HTML | | |
| filtered_probes.tsv<br>binding_map.png<br>thresh_summary.txt<br>binding_quant.txt | **map_cand_region_coords**<br>Adds imaging target coordinates to the candidate probes file. Creates probe binding summary. | final_probes.tsv | | |
| final_probes.tsv<br>probe_summary.txt | **merge_cand_mapping**<br>Aggregates all repeat regions into a single file by scaffold. | probes_merged.tsv | | |
| probes_merged.tsv | **summary**<br>Summarizes probe binding and count by repeat region target. | probe_summary.txt | | |

** Denotes optional step if proper file paths/options are specified in config.yml*
*** Denotes step specific to either probe_design or repeat_ID run modes*
