## Supplementary Software for "Tigerfish designs oligonucleotide-based *in situ* hybridization probes targeting intervals of highly repetitive DNA at the scale of genomes": make.bat

```
@ECHO OFF

pushd %~dp0

REM Command file for Sphinx documentation

if "%SPHINXBUILD%" == "" (
	set SPHINXBUILD=sphinx-build
)
set SOURCEDIR=source
set BUILDDIR=build

%SPHINXBUILD% >NUL 2>NUL
if errorlevel 9009 (
	echo.
	echo.The 'sphinx-build' command was not found. Make sure you have Sphinx
	echo.installed, then set the SPHINXBUILD environment variable to point
	echo.to the full path of the 'sphinx-build' executable. Alternatively you
	echo.may add the Sphinx directory to PATH.
	echo.
	echo.If you don't have Sphinx installed, grab it from
	echo.https://www.sphinx-doc.org/
	exit /b 1
)

if "%1" == "" goto help

%SPHINXBUILD% -M %1 %SOURCEDIR% %BUILDDIR% %SPHINXOPTS% %O%
goto end

:help
%SPHINXBUILD% -M help %SOURCEDIR% %BUILDDIR% %SPHINXOPTS% %O%

:end
popd
```
