## Supplementary Software for "Tigerfish designs oligonucleotide-based *in situ* hybridization probes targeting intervals of highly repetitive DNA at the scale of genomes": conf.py

```
# Configuration file for the Sphinx documentation builder.
#
# This file only contains a selection of the most common options. For a full
# list see the documentation:
# https://www.sphinx-doc.org/en/master/usage/configuration.html

# -- Path setup --------------------------------------------------------------

# If extensions (or modules to document with autodoc) are in another directory,
# add these directories to sys.path here. If the directory is relative to the
# documentation root, use os.path.abspath to make it absolute, like shown here.
#
import os
import sys
import subprocess
from configparser import RawConfigParser
import sphinx_rtd_theme

sys.path.insert(0, os.path.abspath(os.path.join("..", "..")))

#sys.path.insert(0, os.path.abspath('..'))
#sys.path.append(os.path.dirname(__file__))

#try:
#    import numpydoc
#except ModuleNotFoundError:
#    subprocess.run(["pip", "install", "numpydoc"], check=True)


# -- Project information -----------------------------------------------------

project = 'Tigerfish'
copyright = '2022, Robin Aguilar'
author = 'Robin Aguilar'

# The full version, including alpha/beta/rc tags
release = '2022'


# -- General configuration ---------------------------------------------------

#def get_version():
#    """Return package version from setup.cfg."""
#    config = RawConfigParser()
#    config.read(os.path.join('.', 'setup.cfg'))
#    return config.get('metadata', 'version')

# Add any Sphinx extension module names here, as strings. They can be
# extensions coming with Sphinx (named 'sphinx.ext.*') or your custom
# ones.
extensions = [
    'sphinx.ext.duration',
    'sphinx.ext.doctest',
    'sphinx.ext.autodoc',
    'sphinx_tabs.tabs',
    'sphinx-prompt',
    'sphinx_search.extension',
    'sphinx.ext.autosummary',
    'sphinx.ext.intersphinx',
    'sphinxemoji.sphinxemoji'
]


nbsphinx_execute = "never"

#autosummary_generate = True
#autodoc_default_options = {
#    "members": True,
#    "inherited-members": True,
#    "member-order": "bysource",
#}

#numpydoc_show_class_members = True
#numpydoc_show_inherited_class_members = True
#numpydoc_attributes_as_param_list = True
#intersphinx_mapping = {
#    "pandas": ("https://pandas.pydata.org/docs", None),
#    "sklearn": ("https://scikit-learn.org/stable", None),
#    "numpy": ("https://numpy.org/doc/stable", None),
#    "matplotlib": ("https://matplotlib.org", None),
#}

# Add any paths that contain templates here, relative to this directory.
templates_path = ['_templates']


# List of patterns, relative to source directory, that match files and
# directories to ignore when looking for source files.
# This pattern also affects html_static_path and html_extra_path.
exclude_patterns = ["build", "Thumbs.db", ".DS_Store", "**.ipynb_checkpoints"]


# -- Options for HTML output -------------------------------------------------

# The theme to use for HTML and HTML Help pages.  See the documentation for
# a list of builtin themes.
#
html_theme = 'sphinx_rtd_theme'

html_static_path = ["_static"]

master_doc = "index"
source_suffix = [".rst", ".md"]

html_theme_options = {
    "style_nav_header_background": "#343131",
    "logo_only": True,
}

html_css_files = [
    "custom.css",
]

# Add any paths that contain custom static files (such as style sheets) here,
# relative to this directory. They are copied after the builtin static files,
# so a file named "default.css" will overwrite the builtin "default.css".
html_logo = "_static/tigerfish.png"
```
