## Supplementary Software for "Tigerfish designs oligonucleotide-based *in situ* hybridization probes targeting intervals of highly repetitive DNA at the scale of genomes": config.yml

```
#path to genome fasta
fasta_file: "data/example.fa"

#path to file containing primary chromosome sizes
chrom_sizes_file: "data/test_chrom.sizes"

#if coordinates are provided for probe design, file goes here
bed_file: "data/dxz4_synthetic.bed"

#option for probe design that directs pipeline implementation
defined_coords: "FALSE"
repeat_discovery: "FALSE"
probe_cand_binding: "TRUE"

bowtie2_indices_given: "TRUE"
jf_hash_given: "FALSE"
jf_count_given: "FALSE"
chrom_idx_given: "FALSE"
chrom_fasta_given: "FALSE"

assembly: "chm13"
bowtie2_dir: "data/bt2/"
jf_hash_dir: ""
jf_count_dir: ""
chrom_idx_dir: ""
chrom_fasta_dir: ""
probe_cand_file: "data/chr4_X_probe_test.tsv"

#all chromosomes present in bed file or required for probe discovery are listed here
samples:
    - "chr4"
    - "chrX"

#parameters for repeat_ID step
window: 4000

threshold: 5

composition: 0.25

file_start: 0

#parameters for probe_design step
min_length: 36

max_length: 41

min_temp: 42

max_temp: 47

#parameters for kmer_filter script
mer_val: 18

c1_val: 1

c2_val: 5


#parameters used for probe_mer_filter script
enrich_score: 0.50

copy_num: 10


#parameters used in alignment_filter script
genome_windows: 5000000

thresh_window: 100000

binding_prop: 0.70

target_sum: 5000

off_bin_thresh: 100 

mer_cutoff: 0.95

bt2_alignments: 500000

max_pdups_binding: 0.90

seed_length: 15

model_temp: 69.5

min_on_target: 25

max_probe_return: 40

align_thresh: 10

ref_flag: 0
```
