## Supplementary Software for "Tigerfish designs oligonucleotide-based *in situ* hybridization probes targeting intervals of highly repetitive DNA at the scale of genomes": run_pipeline.sh

```
# configure file paths
CONFIG_FILE='config.yml'
SNAKE_FILE='../../workflow/main_pipeline/Snakefile'
CONDA_ENVS='../../shared_conda_envs'

# activate conda environment
source activate snakemake_env                

# run the pipeline
snakemake --configfile config.yml --snakefile $SNAKE_FILE \
    --use-conda --conda-frontend mamba --conda-prefix $CONDA_ENVS --cores \
    --restart-times 3

# export PDF and svg visualizations of the DAG structure of pipeline steps
echo -e "Exporting pipeline DAG to svg and pdf..."
snakemake --configfile config.yml --snakefile $SNAKE_FILE --dag > dag.dot
dot -Tpdf dag.dot > pipeline_output/pipeline.pdf
dot -Tsvg dag.dot > pipeline_output/pipeline.svg
rm dag.dot

# success
echo -e "\nDONE!\n"
```
