## Supplementary Software for "Tigerfish designs oligonucleotide-based *in situ* hybridization probes targeting intervals of highly repetitive DNA at the scale of genomes": README.md

<div align="center">
    <a href="#readme"><img src="../docs/source/imgs/tigerfish_logo.png" width="200"></a>
</div>

## Main Internal Workflow

Here, a genomic fasta which flanks the D4Z4 and DXZ4 repeats is provided as seperate scaffolds (chr4 and chrX) to serve as a toy genome to test Tigerfish functionality. Here, there are three test examples as described:

1. **probe_design_test**: Here, probes are designed against the DXZ4 repeat coordinates using a reference BED file and genome FASTA reference. 

2. **repeat_discovery_test**: Here, probes are designed against both the D4Z4 and DXZ4 repeats. 

3. **probe_candidate_binding_mode**: Once candidate probes satisfy alignment, one could take the output of the alignment_filter step and pursue specific downstream analysis to generate in silico binding predictions about single or pooled probes against a repeat region target. 

4. **probe_design_chm13**: Here, probes are designed against a full HSAT repeat on chr9 in the CHM13 genome. 

All necessary files to run test cases 1 and 2 are found within each contained directory. To test Tigerfish, users just need to run the shell scripts in each respective directory. 

For test case 3 which requires the CHM13 genome FASTA reference (v2.0), tutorials are provided on the Tigerfish ReadtheDocs to download necessary resources to deploy this example. 
