## Supplementary Software for "Tigerfish designs oligonucleotide-based *in situ* hybridization probes targeting intervals of highly repetitive DNA at the scale of genomes": run_check_repeatID.sh

```
python ../../workflow/scripts/check_repeatID.py -ft pipeline_output/03_core_output/chm13_final_probes.txt -xt pipeline_output/03_core_output/chm13_final_probes.txt -fe expected_output/03_core_output/chm13_final_probes.txt -xe expected_output/03_core_output/chm13_final_probes.txt
```
