## Supplementary Software for "Tigerfish designs oligonucleotide-based *in situ* hybridization probes targeting intervals of highly repetitive DNA at the scale of genomes": README.md

<p align="center">
  <img width="300" height="300" src="https://user-images.githubusercontent.com/46250421/132409143-8072bc08-c21f-407d-bc58-9019701a0c7f.png">
</p>

# TigerFISH Pipeline

## Overview

Tigerfish is a technology and software tool that enables users to design oligonucleotide FISH experiments specific for satellite DNA and repetitive DNA families at the scale of genomes. 

Tigergish serves as a pipeline that allows for repetitive probe discovery and design. Users may provide a particular region of interest for oligo probe design, or provide a genomics scaffold for repeat discovery and probe design. Analysis are conducted on candidate probes to determine on and off target probe specificity in silico and generate karyoplots using chromoMap.

An interactive web application for probe design (FISHtank), will include diverse model organism genomes containing information about relevant repetitive DNA probes for FISH experiments is in the works using Tigerfish. FISHtank will be integrated into the UCSC Genome Browser for probe design in newly assembled genomes.

Our documents are publicly found and hosted on Read the Docs.

## System Requirements

Tigerfish is a computational pipeline composed of a collection of Python scripts embedded in an automated Snakemake workflow and is designed to be executed in a POSIX-based command line environment. No direct knowledge of programming is required to run Tigerfish, and this bioinformatic workflow can be deployed on any modern Windows, Macintosh, or Linux system. 

Tigerfish is written in Python 3.7.8 with dependencies that include Biopython 1.77, Bowtie 2.3.5.1, NUPACK 4.0, BEDtools 2.29.2, Numpy 1.18.5, Pandas 1.0.5, pip 20.1.1, pybedtools 0.8.1, sam2pairwise 1.0.0, samtools 1.9, scikit-learn 0.23.1, scipy 1.5.0, zip 3.0, matplotlib 3.3.4, seaborn 0.11.1, pytest 6.2, and Jellyfish 2.2.10.  All Tigerfish probe collections were generated using a pipeline implemented with Snakemake 7.19. Tigerfish ships with all necessary software dependencies and their versions through the conda environment files that are required to run the software. The [tigerfish.yml](https://github.com/beliveau-lab/TigerFISH/tree/master/shared_conda_envs) environment may be found here, the [snakemake_env.yml](https://github.com/beliveau-lab/TigerFISH/blob/master/snakemake_env.yml) environment may be found here, and the [chromomap_env.yml](https://github.com/beliveau-lab/TigerFISH/tree/master/shared_conda_envs) may be found here. 

All data generated in the Tigerfish manuscript was generated on the Genome Sciences SunGrid cluster a CentOS 7.9 cluster node. Specifically, the core node used to generate the Tigerfish oligo probe sets were run on a node with 4x 24-core Intel Xeon 6252 CPUs (2.1GHz), 1.5TB memory, 4x nVidia Tesla M10 GPGPUs.

## Installation

1. Install [conda](https://docs.conda.io/en/latest/miniconda.html) as needed for your system.

2. Proceed with installing Mamba to assist with snakemake installation, as recommended in the [Snakemake installation tutorial](https://snakemake.readthedocs.io/en/stable/getting_started/installation.html#installation-via-conda-mamba).

```
$ conda install -n base -c conda-forge mamba
```

3. Clone this repo, and create and activate the provide environment for implementing Snakemake for Tigerfish:

```
$ git clone https://github.com/beliveau-lab/TigerFISH.git \
    && cd TigerFISH/ \
    && mamba env create -f snakemake_env.yml \
    && conda activate snakemake_env
```
To clone this directory onto a desktop computer, the time to solve this takes approximately 38 seconds.
Typical install time on a desktop computer including environments solving takes approximately 72 secs. 

## Demos and Tutorials

Our comprehensive [Read the Docs page](https://beliveau-lab-tigerfish.readthedocs-hosted.com/en/latest/) contains step-by-step instructions for deploying the test demons on three small datasets that summarize each of the core run modes in Tigerfish (Repeat Discovery Mode, Probe Design Mode, and Probe Analysis Mode) and seperately, a larger real dataset to design oligo probes against the chr9 HSAT repeat in the CHM13 genome. Approximate expected run times and their correspomding expected output directories for each of these demos to complete are described as follows:

- [Repeat Discovery Mode](https://github.com/beliveau-lab/TigerFISH/tree/master/example_run/repeat_discovery_test/expected_output): 110 seconds
- [Probe Design Mode](https://github.com/beliveau-lab/TigerFISH/tree/master/example_run/probe_design_test/expected_output): 83 seconds
- [Probe Analysis Mode](https://github.com/beliveau-lab/TigerFISH/tree/master/example_run/probe_candidate_binding_test/expected_output): 116 seconds
- [CHM13 Probe Design Mode, chr9 HSAT](https://github.com/beliveau-lab/TigerFISH/tree/master/example_run/probe_design_chm13/expected_output): 4 hours (due to generating large CHM13 genome reference files and Bowtie2 indices)

## Documentation

Please read our comprehensive [Read the Docs page and tutorials to learn more about Tigerfish](https://beliveau-lab-tigerfish.readthedocs-hosted.com/en/latest/). Within the Read the Docs page, this includes in-depth installation guides, expected inputs and outputs for expected pipeline performance, a summary of parameters used to generated the datasets featured in the Tigerfish manuscript, definitions of all Tigerfish parameters, reproduction instructions for tutorials and all featured datasets, and a thorough description of the functionality of all scripts called in Snakemake to deploy Tigerfish. 

## Questions

If you have questions or issues, please open an issue on GitHub

## Citation

For usage of the pipeline, please cite according to citation.bib

## License

We provide this open source software without any warranty under the MIT license.
