## Supplementary Software for "Tigerfish designs oligonucleotide-based *in situ* hybridization probes targeting intervals of highly repetitive DNA at the scale of genomes": setup.cfg

```
[metadata]
name = Tigerfish
author = Robin Aguilar
author_email =
description = Fast and efficient repeat specific oligo design workflow for repetitive DNA
long_description = file: README.md
long_description_content_type = text/markdown
url = https://github.com/beliveau-lab/TigerFISH
license = MIT
classifiers =
    Programming Language :: Python :: 3

[options]
packages = find:
python_requires = >=3.6
install_requires =
    numpy>=1.18.1
    pandas>=1.0.3
    biopython>=1.77

[options.extras_require]
docs =
    numpydoc>=1.0.0
    sphinx-argparse>=0.2.5
    sphinx-rtd-theme>=0.5.0
    nbsphinx>=0.7.1
    ipykernel>=5.3.0
    recommonmark>=0.5.0
```
