## Supplementary Software for "Tigerfish designs oligonucleotide-based *in situ* hybridization probes targeting intervals of highly repetitive DNA at the scale of genomes": README.md

# Shared Conda Environments

This snakemake pipeline consists of steps that use [conda](https://docs.conda.io/en/latest/index.html) environments (provided in [workflow/envs/](../workflow/main/envs/)). 

When snakemake is run using the `--use-conda` flag, these environments will be built by default in a `.snakemake/` directory located in the current working directory. As a result, if the pipeline is run in several different working directories (i.e. for several genomes), the default behavior would create identical, redundant environments for each run. 

To prevent this, the `--conda-prefix` argument is passed, pointing to this directory, where each conda environment will only be built once and used for all runs.

To see how to properly invoke snakemake with this conda configuration, see the included [run_pipeline.sh](../example_run/main/main_pipeline/run_pipeline.sh) shell script. To read more about executing snakemake with conda, see the [snakemake documentation](https://snakemake.readthedocs.io/en/v5.20.1/executing/cli.html#CONDA)
