## Supplementary Software for "Tigerfish designs oligonucleotide-based *in situ* hybridization probes targeting intervals of highly repetitive DNA at the scale of genomes": tigerfish.yml

```
name: tigerfish
channels:
  - beliveau-lab
  - bioconda
  - conda-forge
  - defaults
dependencies:
  - bedtools=2.29.2
  - biopython=1.77
  - bowtie2=2.3.5.1
  - jellyfish=2.2.10
  - numpy=1.18.5
  - pandas=1.0.5
  - pip=20.1.1
  - pybedtools=0.8.1
  - sam2pairwise=1.0.0
  - samtools=1.9
  - scikit-learn=0.23.1
  - scipy=1.5.0
  - zip=3.0
  - nupack=4
  - python=3.7.8
  - matplotlib=3.3.4
  - seaborn=0.11.1
  - pytest=6.2
```
