## Supplementary Software for "Tigerfish designs oligonucleotide-based *in situ* hybridization probes targeting intervals of highly repetitive DNA at the scale of genomes": snakemake_env.yml

```
name: snakemake_env
channels:
  - bioconda
  - conda-forge
  - defaults
  - beliveau-lab
dependencies:
  - biopython=1.77
  - gffread=0.12.1
  - pandas=1.1.0
  - pip=20.1.1
  - python=3.8.3
  - snakemake=7.19
  - nupack=4
```
