## Supplementary Software for "Tigerfish designs oligonucleotide-based *in situ* hybridization probes targeting intervals of highly repetitive DNA at the scale of genomes": run_pipeline.sh

```
#!/bin/bash
#$ -S "/bin/bash"
#$ -cwd
#$ -o "job.stdout"
#$ -e "job.stderr"
#$ -l m_mem_free=32.0G
#$ -R y
#$ -l h_rt=0:500:0:0

# configure file paths
CONFIG_FILE=''
SNAKE_FILE=''
CONDA_ENVS=''
WORK_DIR=''

# set working directory
cd $WORK_DIR

# configure cluster jobs
NUM_JOBS=40                 # upper limit on concurrent jobs to submit
LATENCY_WAIT=180            # time in seconds to wait for files missing due to latency
RESTART_ATTEMPTS=1          # number of times to restart any failed job

# helper function
function run_pipeline() {

        # run pipeline
        echo -e "\n~~~~~~~~~~~~~~~~~~~~\n\nTigerfish Pipeline\n\n~~~~~~~~~~~~~~~~~~~~\n"
        echo "WORK_DIR: $WORK_DIR"

        # activate conda env
        source activate snakemake_env

        # run pipeline
        snakemake --snakefile  $SNAKE_FILE --configfile $CONFIG_FILE \
          --conda-prefix $CONDA_ENVS --use-conda --conda-frontend mamba \
          --jobs $NUM_JOBS --latency-wait $LATENCY_WAIT --restart-times $RESTART_ATTEMPTS \
          --cluster "qsub -cwd -l mfree={params.mfree} -l h_rt={params.h_rt} -l centos=7 -R y -e ./pipeline_output/00_logs/stderr.log -o ./pipeline_output/00_logs/stdout.log"

        snakemake --report pipeline_output/report.html
}

# run pipeline
run_pipeline


# if snakemake failed to start, unlock directory and re-try
if [[ $? -ne 0 ]]; then
        snakemake --snakefile $SNAKE_FILE --configfile $CONFIG_FILE --cores all --unlock
        run_pipeline
fi
```
