## Supplementary Software for "Tigerfish designs oligonucleotide-based *in situ* hybridization probes targeting intervals of highly repetitive DNA at the scale of genomes": Snakefile

```
#import libraries

import os
from os import path
import glob

#specify config.yml file
configfile: "config.yml"

#define config parameter names as variables
SAMPLES = config["samples"]
ASSEMBLY = config["assembly"]
BOWTIE2_DIR = config["bowtie2_dir"]
JF_HASH = config['jf_hash_dir']
JF_COUNT = config['jf_count_dir']
CHROM_IDX = config['chrom_idx_dir']
CHROM_FASTA = config['chrom_fasta_dir']

#final output files after pipeline has completed execution
rule all:
    input:
        "pipeline_output/finished/DONE.txt"

#should user specify TRUE or FALSE for both run settings, pipeline exits() and produces message to re-define deployment mode
if (config['defined_coords'] == "TRUE" and config['repeat_discovery'] == "TRUE" and config['probe_cand_binding'] == "TRUE") or (config['defined_coords'] == "FALSE" and config['repeat_discovery'] == "FALSE" and config['probe_cand_binding'] == "FALSE"):

    print("Repeat_Discovery Mode, Probe_Design Mode, and Probe Cand Binding Mode  can only be run independently. Please select TRUE for desired workflow and FALSE for the other process. Exiting ...")
    exit()

#If jellyfish hash table is not given, as specified in config file, Tigerfish will generate a genome wide jellyfish hash table
if config['jf_hash_given'] == "FALSE":
    rule generate_jf_count:
        input:
            fasta_file = config["fasta_file"]
        conda:
            "../../shared_conda_envs/tigerfish.yml"
        params:
            mer = config["mer_val"],
            mfree="60G",
            h_rt = "20:0:0"
        benchmark:
            'pipeline_output/benchmarks/01_reference_files/01_generate_jf_count/jf_count_log.log'
        output:
            'pipeline_output/01_reference_files/01_generate_jf_count/genome_query.jf'
        shell:
            "jellyfish count -s 3300M -m {params.mer} -o {output} -C {input.fasta_file}"

#function provides output if bowtie2 indices are given or if they need to be produced
def input_for_bowtie(wildcards):
    # requires a config containing switches for the whole workflow
    if config["bowtie2_indices_given"]=="TRUE":
        return BOWTIE2_DIR
    elif config["bowtie2_indices_given"]=="FALSE":
        return rules.generate_bt2_indices.output

#function provides output if jellyfish indices need to be produced or are already provided with a directory
def input_for_jf_idx(wildcards):
    #requires a config containing switches for the workflow
    if config['jf_hash_given'] == "TRUE":
        return JF_HASH
    elif config['jf_hash_given'] == "FALSE":
        return rules.generate_jf_count.output

#rule used to generate bowtie2 indices if they are not already provided
if config['bowtie2_indices_given'] == "FALSE":
    BOWTIE2_DIR = f'pipeline_output/01_reference_files/04_generate_bt2_indices/{ASSEMBLY}'
    rule generate_bt2_indices:
        input:
            fasta_file = config["fasta_file"]
        conda:
            "../../shared_conda_envs/tigerfish.yml"
        params:
            mfree="60G",
            h_rt = "20:0:0"
        benchmark:
            'pipeline_output/benchmarks/01_reference_files/04_generate_bt2_indices/bt2_idx_log.log'
        output:
            f'{BOWTIE2_DIR}/{ASSEMBLY}.1.bt2',
            f'{BOWTIE2_DIR}/{ASSEMBLY}.2.bt2',
            f'{BOWTIE2_DIR}/{ASSEMBLY}.3.bt2',
            f'{BOWTIE2_DIR}/{ASSEMBLY}.4.bt2',
            f'{BOWTIE2_DIR}/{ASSEMBLY}.rev.1.bt2',
            f'{BOWTIE2_DIR}/{ASSEMBLY}.rev.2.bt2',
        shell:
            'bowtie2-build --threads 4 {input} {BOWTIE2_DIR}/{ASSEMBLY}'

#function provides output if jellyfish count files need to be produced or if path is given
def input_for_jf_count_file(wildcards):
    #requires config containing switches for the workflow
    if config['jf_count_given'] == "TRUE":
        return JF_COUNT
    elif config['jf_count_given'] == "FALSE":
        return rules.generate_jf_idx.output.jf_count

#function provides output if scaffold indices need to be produced or if path is given
def input_for_chrom_idx_file(wildcards):
    #requires config containing switches for the workflow
    if config['chrom_idx_given']  == "TRUE":
        return CHROM_IDX
    elif config['chrom_idx_given'] == "FALSE":
        return rules.generate_jf_idx.output.chrom_idx

#function provides output if scaffold FASTA files are given in dir or if they need to be created
def input_for_chrom_fasta_file(wildcards):
    #requires config containing switches for the workflow
    if config['chrom_fasta_given']  == "TRUE":
        return CHROM_FASTA
    elif config['chrom_fasta_given'] == "FALSE":
        return rules.generate_jf_idx.output.chrom_fa

#rule directs if jellyfish count, scaffold index, and scaffold FASTA files need to be produced if no jellyfish hash was generated previously
if config['jf_hash_given'] == 'FALSE':
    rule generate_jf_idx:
        input:
            fasta_file = config["fasta_file"],
            jf = input_for_jf_idx
        conda:
            "../../shared_conda_envs/tigerfish.yml"
        params:
            chrom_name = "{sample}",
            mer = config["mer_val"],
            mfree="60G",
            h_rt = "20:0:0"
        benchmark:
            'pipeline_output/benchmarks/01_reference_files/03_generate_jf_idx/{sample}_log.log'
        output:
            jf_count = 'pipeline_output/01_reference_files/03_generate_jf_idx/{sample}_jf_temp.txt',
            chrom_idx = 'pipeline_output/01_reference_files/03_generate_jf_idx/{sample}_index.txt',
            chrom_fa = 'pipeline_output/01_reference_files/03_generate_jf_idx/repeat_fasta/{sample}.fa'
        shell:
            "python ../../workflow/scripts/generate_jf_idx.py -f {input.fasta_file} -j {input.jf} -c {params.chrom_name} -m {params.mer} -f_o {output.chrom_fa} -j_o {output.jf_count} -i {output.chrom_idx}"

#rule dictates behavior if user provides BED file and specifies defined_coords run mode
if config['defined_coords'] == "TRUE":
    rule split_bed:
        input:
            bed_file = config["bed_file"]
        conda:
            "../../shared_conda_envs/tigerfish.yml"
        params:
            chrom_name = "{sample}",
            mfree="10G",
            h_rt = "3:0:0"
        benchmark:
            'pipeline_output/benchmarks/02_intermediate_files/01_split_bed/{sample}_log.log'
        output:
            chrom_region = 'pipeline_output/02_intermediate_files/01_split_bed/{sample}_regions.bed'
        shell:
            "python ../../workflow/scripts/split_bed.py -b {input.bed_file} -c {params.chrom_name} -o {output.chrom_region}"

#rule dictates behavior if users define repeat_discovery mode
if config['repeat_discovery'] == "TRUE":
    rule repeat_ID:
        input:
            jf_count = input_for_jf_count_file,
            chrom_index = input_for_chrom_idx_file
        conda:
            "../../shared_conda_envs/tigerfish.yml"
        params:
            window = config["window"],
            threshold = config["threshold"],
            composition = config["composition"],
            file_start = config["file_start"],
            chrom_name = "{sample}",
            mer = config["mer_val"],
            mfree="85G",
            h_rt="200:0:0"
        benchmark:
            "pipeline_output/benchmarks/02_intermediate_files/01_repeat_ID/{sample}_log.log"
        output:
            out_bed = "pipeline_output/02_intermediate_files/01_repeat_ID/{sample}_regions.bed",
        shell:
            "python ../../workflow/scripts/repeat_ID.py -j {input.jf_count} -i {input.chrom_index} -m {params.mer} -w {params.window} -t {params.threshold} -c {params.composition} -chr {params.chrom_name} -st {params.file_start} -o_b {output.out_bed}"

#function defines which files are to be returned to initiate probe design
def input_for_design_probes(wildcards):
    # requires a config containing switches for the whole workflow
    if config["defined_coords"]=='TRUE':
        return 'pipeline_output/02_intermediate_files/01_split_bed/{sample}_regions.bed'
    elif config["repeat_discovery"]=='TRUE':
        return 'pipeline_output/02_intermediate_files/01_repeat_ID/{sample}_regions.bed'

#rule takes output created from input_for_design_probes and designs probes against those BED regions
rule design_probes:
    input:
        region_bed = input_for_design_probes,
        chr_path = input_for_chrom_fasta_file
    conda:
        "../../shared_conda_envs/tigerfish.yml"
    params:
        mfree="60G",
        h_rt="200:0:0",
        chrom_name = "{sample}",
        min_length = config["min_length"],
        max_length = config["max_length"],
        min_temp = config["min_temp"],
        max_temp = config["max_temp"]
    benchmark:
        "pipeline_output/benchmarks/02_intermediate_files/02_design_probes/{sample}_log.log"
    output:
        designed_probes = "pipeline_output/02_intermediate_files/02_design_probes/{sample}_blockParse_probe_df.bed",
        probe_fa = "pipeline_output/02_intermediate_files/02_design_probes/{sample}_probe_regions.fa"
    shell:
        "python ../../workflow/scripts/design_probes.py -b {input.region_bed} -c {params.chrom_name} -g {input.chr_path} -p_o {output.designed_probes} -r_o {output.probe_fa} -l {params.min_length} -L {params.max_length} -t {params.min_temp} -T {params.max_temp}"

#rule performs k-mer filtering on designed probes based on user defined parameters
rule kmer_filter:
    input:
        jf = input_for_jf_count_file,
        probes = rules.design_probes.output.designed_probes,
        region_fa = rules.design_probes.output.probe_fa
    conda:
        "../../shared_conda_envs/tigerfish.yml"
    params:
        mfree="50G",
        h_rt="200:0:0",
        mer = config["mer_val"],
        chrom_name = "{sample}",
        c1 = config["c1_val"],
        c2 = config["c2_val"]
    benchmark:
        "pipeline_output/benchmarks/02_intermediate_files/03_kmer_filter/{sample}_log.log"
    output:
        "pipeline_output/02_intermediate_files/03_kmer_filter/{sample}_probes_pre_filter.txt"
    shell:
        "python ../../workflow/scripts/kmer_filter.py -p {input.probes} -o {output} -j {input.jf} -f {input.region_fa} -m {params.mer} -c1 {params.c1} -c2 {params.c2} -c {params.chrom_name}"

#rule filters probes further and rank sorts them based on user defined parameters
rule probe_mer_filter:
    input:
        probes = rules.kmer_filter.output
    conda:
        "../../shared_conda_envs/tigerfish.yml"
    params:
        mfree="50G",
        h_rt="200:0:0",
        mer = config["mer_val"],
        enrich = config["enrich_score"],
        copy_num = config["copy_num"],
        mer_cutoff = config["mer_cutoff"]
    benchmark:
        "pipeline_output/benchmarks/02_intermediate_files/04_probe_mer_filter/{sample}_log.log"
    output:
        "pipeline_output/02_intermediate_files/04_probe_mer_filter/{sample}_probes_mer_filter.txt"
    shell:
        "python ../../workflow/scripts/probe_mer_filter.py -f {input.probes} -o {output} -e {params.enrich} -cn {params.copy_num} -m {params.mer_cutoff} -k {params.mer}"

#rule generates genomic bins using BEDtools which will be used to validate alignment and candidate probe binding specificity
rule generate_genome_bins:
    input:
        sizes = config["chrom_sizes_file"]
    output:
        alignment_bins = "pipeline_output/01_reference_files/02_generate_genome_bins/genome_bins_tigerfish_alignment.bed",
        threshold_bins = "pipeline_output/01_reference_files/02_generate_genome_bins/genome_bins_tigerfish_threshold.bed"
    conda:
        "../../shared_conda_envs/tigerfish.yml"
    params:
        mfree="20G",
        h_rt = "10:0:0",
        genome = config['assembly'],
        window = config['genome_windows'],
        thresh_window = config['thresh_window']
    benchmark:
        "pipeline_output/benchmarks/01_reference_files/02_generate_genome_bins/genome_bins.txt"
    shell:
        """
        bedtools makewindows -g {input.sizes} -w {params.window} > {output.alignment_bins}
        bedtools makewindows -g {input.sizes} -w {params.thresh_window} > {output.threshold_bins}
        """

#rule is used to split all repeat regions found within one scaffold into independent repeat region specific files that will undergo alignment
checkpoint make_chrom_dir:
    input:
       probes = rules.probe_mer_filter.output
    conda:
        "../../shared_conda_envs/tigerfish.yml"
    params:
        mfree="5G",
        h_rt="350:0:0"
    benchmark:
        "pipeline_output/benchmarks/02_intermediate_files/05_make_chrom_dir/{sample}.log"
    output:
        region_split = directory("pipeline_output/02_intermediate_files/05_make_chrom_dir/{sample}/")
    shell:
        """
        python ../../workflow/scripts/split_filter.py -f {input.probes} -o {output}
        """
#rule takes each repeat region created from the checkpoint and performs alignment-based filtering analysis to isolate final candidate probes
rule alignment_filter:
    input:
        probe_files = "pipeline_output/02_intermediate_files/05_make_chrom_dir/{sample}/{region}.txt",
        genome_bins = rules.generate_genome_bins.output.alignment_bins,
        BOWTIE2_DIR = input_for_bowtie
    conda:
        "../../shared_conda_envs/tigerfish.yml"
    params:
        mfree="25G",
        h_rt="350:0:0",
        region_thresh = config['target_sum'], 
        k_val = config['bt2_alignments'],
        max_pdups_binding = config['max_pdups_binding'],
        seed_length = config["seed_length"],
        model_temp = config["model_temp"],
        min_on_target = config["min_on_target"],
        max_probe_return = config["max_probe_return"],
        off_bin_thresh = config["off_bin_thresh"],
        binding_prop = config['binding_prop'],
        ref_flag = config['ref_flag']
    benchmark:
        "pipeline_output/benchmarks/02_intermediate_files/06_alignment_filter/{sample}/{region}.log"
    output:
        "pipeline_output/02_intermediate_files/06_alignment_filter/{sample}/{region}_alignment.txt"
    shell:
        "python ../../workflow/scripts/alignment_filter.py -f {input.probe_files} -b {BOWTIE2_DIR}/{ASSEMBLY} -o {output} -r {params.region_thresh} -p {params.binding_prop} -k {params.k_val} -pb {params.max_pdups_binding} -l {params.seed_length} -t {params.model_temp} -moT {params.min_on_target} -Mr {params.max_probe_return} -gb {input.genome_bins} -th {params.off_bin_thresh} -rf {params.ref_flag}"

#function will aggregate all repeat regions that complete the alignment_filter process
def aggregate_alignment_input(wildcards):
    checkpoint_output = checkpoints.make_chrom_dir.get(**wildcards).output[0]
    return expand("pipeline_output/02_intermediate_files/06_alignment_filter/{sample}/{region}_alignment.txt",
           sample=wildcards.sample,
           region=glob_wildcards(os.path.join(checkpoint_output, "{region}.txt")).region)

#function aggregates all file paths collected by function to merge back into a single scaffold
rule merge_alignment_filter:
    input:
        aggregate_alignment_input
    output:
        'pipeline_output/02_intermediate_files/07_merge_alignment_filter/{sample}_alignments.tsv'
    conda:
        "../../shared_conda_envs/tigerfish.yml"
    params:
        mfree='3G',
        h_rt='3:0:0'
    benchmark:
        "pipeline_output/benchmarks/02_intermediate_files/07_merge_alignment_filter/{sample}.log"
    shell:
        "cat {input} > {output}"

#checkpoint then splits all regions remaining into seperate files in new directory
checkpoint split_rm_alignments:
    input:
        rules.merge_alignment_filter.output
    output:
        region_split = directory("pipeline_output/02_intermediate_files/08_split_rm_alignments/{sample}/")
    conda:
        "../../shared_conda_envs/tigerfish.yml"
    params:
        mfree='3G',
        h_rt='3:0:0'
    benchmark:
        "pipeline_output/benchmarks/02_intermediate_files/08_split_rm_alignments/{sample}.log"
    shell:
        "python ../../workflow/scripts/split_rm_alignments.py -f {input} -o {output}"

#rule proceeds with independent region probe alignment for all final candidate probes contained within each repeat region
rule align_probes:
    input:
        "pipeline_output/02_intermediate_files/08_split_rm_alignments/{sample}/{region}_alignments.txt"
    output:
        "pipeline_output/02_intermediate_files/09_align_probes/{sample}/{region}_probe_alignment.txt"
    conda:
        "../../shared_conda_envs/tigerfish.yml"
    params:
        mfree="20G",
        h_rt = "10:0:0",
        k_val = config["bt2_alignments"],
        seed_length = config["seed_length"],
        model_temp = config["model_temp"]
    benchmark:
        "pipeline_output/benchmarks/02_intermediate_files/09_align_probes/{sample}/{region}_alignment.txt"
    shell:
        'python ../../workflow/scripts/generate_alignments.py -f {input} -o {output} -b {BOWTIE2_DIR}/{ASSEMBLY} -k {params.k_val} -l {params.seed_length} -t {params.model_temp}'

#rule taked SAM output to identify where reported alignments are located as BED regions
rule derived_beds:
    input:
        rules.align_probes.output
    output:
        "pipeline_output/02_intermediate_files/10_derived_beds/{sample}/{region}_derived.bed"
    conda:
        "../../shared_conda_envs/tigerfish.yml"
    params:
        mfree="20G",
        h_rt = "10:0:0"
    benchmark:
        "pipeline_output/benchmarks/02_intermediate_files/10_derived_beds/{sample}/{region}_derived.txt"
    shell:
        "python ../../workflow/scripts/make_derived_beds.py -f {input} -o {output}"

#rule takes alignment file and creates BED file from target repeat region
rule get_region_bed:
    input:
        rules.alignment_filter.output
    output:
        "pipeline_output/02_intermediate_files/11_get_repeat_bed/{sample}/{region}.bed"
    conda:
        "../../shared_conda_envs/tigerfish.yml"
    params:
        mfree="20G",
        h_rt = "10:0:0"
    benchmark:
        "pipeline_output/benchmarks/02_intermediate_files/11_get_repeat_bed/{sample}/{region}.txt"
    shell:
        "python ../../workflow/scripts/get_region_bed.py -i {input} -o {output}"

#rule intersects derived alignments and repeat region with genomic bins to identify where reported alignments map (within target repeat vs outside of target repeat)
rule bedtools_intersect:
    input:
        derived_bed = rules.derived_beds.output,
        genome_bin = rules.generate_genome_bins.output.threshold_bins,
        repeat_bed = rules.get_region_bed.output
    output:
        alignments_out = "pipeline_output/02_intermediate_files/12_bedtools_intersect/{sample}/{region}_intersect.txt",
        repeat_out = "pipeline_output/02_intermediate_files/12_bedtools_intersect/{sample}/{region}_repeat_intersect.txt"
    conda:
        "../../shared_conda_envs/tigerfish.yml"
    params:
        mfree="20G",
        h_rt = "320:0:0"
    benchmark:
        "pipeline_output/benchmarks/02_intermediate_files/12_bedtools_intersect/{sample}/{region}_derived.txt"
    shell:
        "bedtools intersect -wa -wb -a {input.derived_bed} -b {input.genome_bin} > {output.alignments_out} |"
        "bedtools intersect -wa -wb -a {input.repeat_bed} -b {input.genome_bin} > {output.repeat_out}"

#rule to report where repeat region probes bind in the genome, provides thresholded binding based on user parameters to isolate imaging target
rule get_alignments:
    input:
        alignment_intersect = rules.bedtools_intersect.output.alignments_out,
        region_intersect = rules.bedtools_intersect.output.repeat_out,
        probes_alignment = rules.align_probes.output,
        genome_bin = rules.generate_genome_bins.output.threshold_bins
    output:
        target_binding = "pipeline_output/04_supplementary_output/01_genome_wide_binding/{sample}/{region}_alignment_binding.txt",
        thresh_binding = "pipeline_output/04_supplementary_output/02_threshold_binding/{sample}/{region}_thresh_binding.txt",
        binding_maps = "pipeline_output/04_supplementary_output/03_genome_wide_binding_plots/{sample}/{region}_genome_view.png"
    conda:
        "../../shared_conda_envs/tigerfish.yml"
    benchmark:
        "pipeline_output/benchmarks/04_supplementary_output/02_get_alignments/{sample}/{region}_alignments.txt"
    params:
        mfree="20G",
        h_rt = "320:0:0",
        thresh = config["align_thresh"]    
    shell:
        "python ../../workflow/scripts/get_alignments.py -c_t {input.genome_bin} -c_o {input.alignment_intersect} -p {input.probes_alignment} -r_o {input.region_intersect} -pl {output.binding_maps} -t {params.thresh} -t_s {output.thresh_binding} -c_s {output.target_binding}"

#function will take all probes after their imaging target region has been appended to aggregate their paths
def all_mapped_output(wildcards):
    checkpoint_output = checkpoints.split_rm_alignments.get(**wildcards).output[0]
    return expand("pipeline_output/04_supplementary_output/04_region_probes/{sample}/{region}_probes.txt",
           sample=wildcards.sample,
           region=glob_wildcards(os.path.join(checkpoint_output, "{region}_alignments.txt")).region)

#rule append imaging target regions and summarize binding information and probe quantity within each target repeat
rule map_region_coords:
    input:
        probe_file = rules.alignment_filter.output,
        thresh_file = rules.get_alignments.output.thresh_binding,
        alignment_file = rules.align_probes.output
    output:
        mod_probe_file = 'pipeline_output/04_supplementary_output/04_region_probes/{sample}/{region}_probes.txt',
        repeat_binding_summ = 'pipeline_output/04_supplementary_output/05_region_binding_summaries/{sample}/{region}_binding_summ.txt'
    conda:
        "../../shared_conda_envs/tigerfish.yml"
    params:
        mfree="20G",
        h_rt = "10:0:0"
    benchmark:
        "pipeline_output/benchmarks/04_supplementary_output/03_map_region/{sample}/{region}.txt"
    shell:
        "python ../../workflow/scripts/collapse_repeat.py -f {input.thresh_file} -a {input.alignment_file} -p {input.probe_file} -po {output.mod_probe_file} -ro {output.repeat_binding_summ}"

#rule will merge the individually-flattened chromosome annotation files
rule merge_mapping:
    input:
        all_mapped_output
    output:
        'pipeline_output/04_supplementary_output/06_chrom_candidate_probes/{sample}_final_probes.txt'
    conda:
        "../../shared_conda_envs/tigerfish.yml"
    params:
        mfree='3G',
        h_rt='3:0:0'
    shell:
        "cat {input} > {output}"

rule merge_chrom_mapping:
    input:
        expand("pipeline_output/04_supplementary_output/06_chrom_candidate_probes/{sample}_final_probes.txt", sample=SAMPLES)
    output:
        f'pipeline_output/03_core_output/{config["assembly"]}_final_probes.txt'
    conda:
        "../../shared_conda_envs/tigerfish.yml"
    params:
        mfree='3G',
        h_rt='3:0:0'
    shell:
        """
        cat {input} | sort -k1,1 -V -s > {output}
        """

#ruke performs a scaffold wide summary of total repeat regions, probes in each region, and on-target binding specificity
rule summary:
    input:
        rules.merge_chrom_mapping.output
    output:
        f'pipeline_output/03_core_output/{config["assembly"]}_final_probe_summary.txt'
    conda:
        "../../shared_conda_envs/tigerfish.yml"
    params:
        mfree="3G",
        h_rt="200:0:0"
    shell:
        "python ../../workflow/scripts/finish_summary.py -f {input} -o {output}"

#rule takes output of summary to finish as final file that ends Tigerfish pipeline

if config['probe_cand_binding'] != "TRUE":

    rule finish:
        input:
            rules.summary.output
        output:
            'pipeline_output/finished/DONE.txt'
        params:
            mfree = '3G',
            h_rt = '3:0:0'
        shell:
            'touch {output}'

########################################################################################################

if config['probe_cand_binding'] == "TRUE": 

    rule gather_repeat_regions:
        input:
            config['probe_cand_file']
        conda:
            "../../shared_conda_envs/tigerfish.yml"
        params:
            mfree="10G",
            h_rt="200:0:0",
            chrom_name = "{sample}"
        benchmark:
            "pipeline_output/benchmarks/02_intermediate_files/01_split_regions/{sample}_log.log"
        output:
            "pipeline_output/02_intermediate_files/01_split_regions/{sample}_probes.txt"
        shell:
            "python ../../workflow/scripts/split_filter_region.py -f {input} -o {output} -c {params.chrom_name}"

    rule align_cand_probes:
        input:
            rules.gather_repeat_regions.output
        output:
            "pipeline_output/02_intermediate_files/02_align_probes/{sample}_probe_alignment.txt"
        conda:
            "../../shared_conda_envs/tigerfish.yml"
        params:
            mfree="20G",
            h_rt = "10:0:0",
            k_val = config["bt2_alignments"],
            seed_length = config["seed_length"],
            model_temp = config["model_temp"]
        benchmark:
            "pipeline_output/benchmarks/02_intermediate_files/02_align_probes/{sample}_alignment.txt"
        shell:
            'python ../../workflow/scripts/generate_alignments.py -f {input} -o {output} -b {BOWTIE2_DIR}/{ASSEMBLY} -k {params.k_val} -l {params.seed_length} -t {params.model_temp}'

    rule derived_cand_beds:
        input:
            rules.align_cand_probes.output
        output:
            "pipeline_output/02_intermediate_files/03_derived_beds/{sample}_derived.bed"
        conda:
            "../../shared_conda_envs/tigerfish.yml"
        params:
            mfree="20G",
            h_rt = "10:0:0"
        benchmark:
            "pipeline_output/benchmarks/02_intermediate_files/03_derived_beds/{sample}_derived.txt"
        shell:
            "python ../../workflow/scripts/make_derived_beds.py -f {input} -o {output}"

    rule get_cand_region_bed:
        input:
            config['probe_cand_file']
        output:
            "pipeline_output/02_intermediate_files/04_get_repeat_bed/{sample}.bed"
        conda:
            "../../shared_conda_envs/tigerfish.yml"
        params:
            mfree="20G",
            h_rt = "10:0:0"
        benchmark:
            "pipeline_output/benchmarks/02_intermediate_files/04_get_repeat_bed/{sample}.txt"
        shell:
            "python ../../workflow/scripts/get_region_bed.py -i {input} -o {output}"

    rule bedtools_intersect_cands:
        input:
            derived_bed = rules.derived_cand_beds.output,
            genome_bin = rules.generate_genome_bins.output.threshold_bins,
            repeat_bed = rules.get_cand_region_bed.output
        output:
            alignments_out = "pipeline_output/02_intermediate_files/05_bedtools_intersect/{sample}_intersect.txt",
            repeat_out = "pipeline_output/02_intermediate_files/05_bedtools_intersect/{sample}_repeat_intersect.txt"
        conda:
            "../../shared_conda_envs/tigerfish.yml"
        params:
            mfree="20G",
            h_rt = "320:0:0"
        benchmark:
            "pipeline_output/benchmarks/02_intermediate_files/05_bedtools_intersect/{sample}_derived.txt"
        shell:
            "bedtools intersect -wa -wb -a {input.derived_bed} -b {input.genome_bin} > {output.alignments_out} |"
            "bedtools intersect -wa -wb -a {input.repeat_bed} -b {input.genome_bin} > {output.repeat_out}"

    rule get_cand_alignments:
        input:
            alignment_intersect = rules.bedtools_intersect_cands.output.alignments_out,
            region_intersect = rules.bedtools_intersect_cands.output.repeat_out,
            probes_alignment = rules.align_cand_probes.output,
            genome_bin = rules.generate_genome_bins.output.threshold_bins
        output:
            target_binding = "pipeline_output/04_supplementary_output/01_genome_wide_binding/{sample}_alignment_binding.txt",
            thresh_binding = "pipeline_output/04_supplementary_output/02_threshold_binding/{sample}_thresh_binding.txt",
            binding_maps = "pipeline_output/04_supplementary_output/03_genome_wide_binding_plots/{sample}_genome_view.png"
        conda:
            "../../shared_conda_envs/tigerfish.yml"
        benchmark:
            "pipeline_output/benchmarks/04_supplementary_output/02_get_alignments/{sample}_alignments.txt"
        params:
            mfree="20G",
            h_rt = "320:0:0",
            thresh = config["align_thresh"]
        shell:
            "python ../../workflow/scripts/get_alignments.py -c_t {input.genome_bin} -c_o {input.alignment_intersect} -p {input.probes_alignment} -r_o {input.region_intersect} -pl {output.binding_maps} -t {params.thresh} -t_s {output.thresh_binding} -c_s {output.target_binding}"

    rule generate_cand_chromomap:
        input:
            chrom_sizes = config["chrom_sizes_file"],
            region_bed = rules.get_cand_region_bed.output,
            binding_maps = rules.get_cand_alignments.output.binding_maps
        output:
            "pipeline_output/04_supplementary_output/06_generate_chromomap/{sample}_chromomap.html"
        conda:
            "../../shared_conda_envs/chromomap_env.yml"
        params:
            mfree="20G",
            h_rt = "10:0:0"
        benchmark:
            "pipeline_output/benchmarks/03_output_files/02_supplementary_output/06_generate_chromomap/{sample}_plots.txt"
        shell:
            "Rscript --vanilla ../../workflow/scripts/make_chromomap.R -c {input.chrom_sizes} -r {input.region_bed} -o {output}"

    rule map_cand_region_coords:
        input:
            probe_file = rules.gather_repeat_regions.output,
            thresh_file = rules.get_cand_alignments.output.thresh_binding,
            alignment_file = rules.align_cand_probes.output,
            chromomap = rules.generate_cand_chromomap.output
        output:
            mod_probe_file = 'pipeline_output/04_supplementary_output/04_region_probes/{sample}_probes.txt',
            repeat_binding_summ = 'pipeline_output/04_supplementary_output/05_repeat_binding_summaries/{sample}_thresh_summ.txt'
        conda:
            "../../shared_conda_envs/tigerfish.yml"
        params:
            mfree="20G",
            h_rt = "10:0:0"
        benchmark:
            "pipeline_output/benchmarks/04_supplementary_output/03_map_region/{sample}.txt"
        shell:
            "python ../../workflow/scripts/collapse_repeat.py -f {input.thresh_file} -a {input.alignment_file} -p {input.probe_file} -po {output.mod_probe_file} -ro {output.repeat_binding_summ}"

    rule merge_cand_mapping:
        input:
            rules.map_cand_region_coords.output.mod_probe_file
        output:
            'pipeline_output/04_supplementary_output/06_chrom_candidate_probes/{sample}_final_probes.txt'
        conda:
            "../../shared_conda_envs/tigerfish.yml"
        benchmark:
            "pipeline_output/benchmarks/03_core_output/{sample}_probes.txt"
        params:
            mfree='3G',
            h_rt='3:0:0'
        shell:
            "cat {input} > {output}"

    rule merge_cand_chrom_mapping:
        input:
            expand("pipeline_output/04_supplementary_output/06_chrom_candidate_probes/{sample}_final_probes.txt", sample=SAMPLES)
        output:
            f'pipeline_output/03_core_output/{config["assembly"]}_final_probes.txt'
        conda:
            "../../shared_conda_envs/tigerfish.yml"
        params:
            mfree='3G',
            h_rt='3:0:0'
        shell:
            """
            cat {input} | sort -k1,1 -V -s > {output}
            """

    rule summary_cand_analysis:
        input:
            mapping_out = rules.merge_cand_chrom_mapping.output
        output:
            f'pipeline_output/03_core_output/{config["assembly"]}_final_probe_summary.txt'
        conda:
            "../../shared_conda_envs/tigerfish.yml"
        params:
            mfree="3G",
            h_rt="200:0:0"
        shell:
            "python ../../workflow/scripts/finish_summary.py -f {input.mapping_out} -o {output}"

    rule finish_cand_analysis:
        input:
            rules.summary_cand_analysis.output
        output:
            'pipeline_output/finished/DONE.txt'
        params:
            mfree = '3G',
            h_rt = '3:0:0'
        shell:
            'touch {output}'
```
