## Supplementary Software for "Tigerfish designs oligonucleotide-based *in situ* hybridization probes targeting intervals of highly repetitive DNA at the scale of genomes": README.md

<div align="center">
    <a href="#readme"><img src="../docs/source/imgs/tigerfish_logo.png" width="200"></a>
</div>

## Snakefiles, Scripts, and Conda Environments

Here, each directory contains specific file paths to Snakemake files for each implementation of Tigerfish in the main workflow, conda environments used to run scripts, and detailed scripts used in each pipeline.

* [scripts used in Snakefiles](scripts/)
* [Snakemake file](main_pipeline/Snakefile)
* [conda environments used](../shared_conda_envs)
