## Supplementary Software for "Tigerfish designs oligonucleotide-based *in situ* hybridization probes targeting intervals of highly repetitive DNA at the scale of genomes": alignment_filter.py

```
#!/usr/bin/env python3
# -*- coding: utf-8 -*-
##############################################################################
"""
Created on Tues March 29
@author: Robin Aguilar
Beliveau and Noble Labs
University of Washington | Department of Genome Sciences
"""
##############################################################################

import argparse
import time
import pandas as pd
import nupack
from nupack import *
import tempfile
from Bio.Seq import reverse_complement as rev_comp
import subprocess
from subprocess import Popen
import itertools
from Bio.Seq import Seq
import os

##############################################################################

def read_probe_filter(p_file):
    """
    Function takes probe file, adjusts probe start and end to coordinates
    that match the repeat region syntax. Then removes redundant columns 
    before returning the file as a dataframe.
    Parameters
    ----------
    p_file : file
        contains the probe information after filtering
    Returns
    -------
    probe_df : dataframe
        returns probe information as a pandas dataframe
    """

    #read in the file with the appropriate column names
    colnames = ["chrom","p_start","p_end","probe","Tm","region",
                "r_count","h_count","k_score","k_norm"]

    #read as a dataframe
    probe_df = pd.read_csv(p_file, delimiter = '\t', names = colnames)

    probe_df[['chrom','region']] = probe_df['region'].str.split(':',
                expand=True)

    probe_df[['r_start','r_end']] = probe_df['region'].str.split('-',
                expand=True)

    #cast r columns as ints
    probe_df[['r_start','r_end']] = probe_df[['r_start',
                'r_end']].astype(int)

    # column for chrom:start-stop that matches true genomic coords
    probe_df['start'] = probe_df['p_start'] + probe_df['r_start']
    probe_df['end'] = probe_df['p_end'] + probe_df['r_start']

    probe_coords_list = []
    #make lists to make a new region
    chrom_list = probe_df['chrom'].tolist()
    start_list = probe_df['start'].tolist()
    end_list = probe_df['end'].tolist()

    #make the probe coords column
    for chrom,start,end in zip(chrom_list,start_list,end_list):
        probe_coords_list.append('%s:%s-%s' % (chrom,start,end))

    probe_df['probe_coords'] = probe_coords_list

    #now do the same for region again
    r_start_list = probe_df['r_start'].tolist()
    r_end_list = probe_df['r_end'].tolist()

    r_coords_list = []
    for chrom,r_start,r_end in zip(chrom_list,r_start_list,r_end_list):
        r_coords_list.append('%s:%s-%s' % (chrom,r_start,r_end))

    probe_df['region'] = r_coords_list

    #cull columns
    columns = ['chrom','p_start','p_end','r_start','r_end','start','end']
    probe_df = probe_df.drop(columns,axis=1)

    #rearrange cols
    probe_df = probe_df[['probe_coords','region','probe','Tm','r_count',
                         'h_count','k_score','k_norm']]

    return probe_df

##############################################################################

def filter_thresh(probe_df,strand_conc_a,strand_conc_b,
                               bowtie_idx,r_thresh,bowtie_string,
                               NUPACK_MODEL,bt2_k_val,max_pdups_binding,
                               seed_length,max_probe_return,min_on_target,
                               genomic_bins,thresh,pdups_p,ref_flag):
    """
    Function implements filter by returning probe cands until on target sum
    for a target region is reached.
    Parameters
    ----------
    probe_df : dataframe
        contains probes that have been filtered by mer count and uniqeness
    Returns
    -------
    on_target_dict : dictionary
        the on target alignment pdups aggregate sum for any probe within
        its repeat region target
    off_target_dict : dictionary
        the off target alignment pdups aggregate sum for any probe outside of 
        its target repeat reguin.
    prop_target_dict : dictionary
        the proportion of on_target pdups binding sum/ all pdups binding
        for all probes
    """

    #need to make lists to store the probe names, on target, off target, prop
    keep_probe_names_list = []
    keep_on_target_list = []
    keep_off_target_list = []
    keep_probe_prop_list = []
    keep_probe_seq = []
    skip_probe_names_list = []
    fail_probe_names_list = []
    all_run_probes_names_list = []
    probe_times = []

    loop_count = 0

    #initiate groupby

    #this takes into account single instances of probe in repeat
    probe_list = probe_df['probe'].tolist()
    probe_coords_list = probe_df['probe_coords'].tolist()

    #make a list of the repeat regions
    probe_regions_list = probe_df['region'].tolist()

    #sets probe coords per region as a dictionary
    region_dict = dict(zip(probe_coords_list,probe_regions_list))

    #keeps track of the on target threshold sum for the region
    threshold_count = 0

    #while the threshold count is below the param requested and 
    #the length of the probe list is greater than 1
    while (threshold_count <= r_thresh and 
           len(keep_probe_names_list) < int(max_probe_return) and 
           len(probe_list) >= 1):

        t0 = time.time()

        #make the call for the top probe to get the pairwise df
        top_probe_al = generate_pairwise_df(probe_list[0],
                                                  probe_coords_list[0],
                                                  bowtie_idx,
                                                  bowtie_string,
                                                  strand_conc_a,
                                                  strand_conc_b,
                                                  NUPACK_MODEL,
                                                  bt2_k_val,seed_length,ref_flag)

        #compute the on target sum for the top probe
        prop_dict,on_target_dict,off_target_dict = nupack_sum(top_probe_al,
                                                                  region_dict,ref_flag)

        #function that makes a temp .bed of alignments and gets overlap
        #embedded function that takes both overlap inputs to return pileup
        #returns: agg by chrom, and pileup subset if pileup ok, then proceed
        signal_pileup_subset,chrom_summ_df = get_bedtools_map(probe_regions_list[0],probe_coords_list[0],
                                                                  genomic_bins,
                                                                  top_probe_al,
                                                                  thresh,ref_flag)
        loop_count += 1

        #checks the items in the proportion dictionary
        #and this is the first item  to be added into the empty list
        #evals if any bins are off target
        for key,val in prop_dict.items():
            if (val >= float(pdups_p) and ((signal_pileup_subset['bin_type'] == 1).all() == True) and
                (on_target_dict[key] >= int(min_on_target))):

                #append probe and it's relevant on target and off
                #target information
                keep_probe_names_list.append(key)
                keep_on_target_list.append(on_target_dict[key])
                keep_off_target_list.append(off_target_dict[key])
                keep_probe_prop_list.append(prop_dict[key])
                keep_probe_seq.append(probe_list[0])

                #add the on target aggregate pdups sum to update 
                #the threshold count
                threshold_count += on_target_dict[key]

            else:

                top_probe_list = []
                pdups_val_list = []

                #stores the probe that failed into the failed list                    
                fail_probe_names_list.append(probe_coords_list[0])

                #takes the length of the probe list to populate it with failed cand
                for i in range(len(probe_list)-1):
                    top_probe_list.append(probe_list[0])

                #computes pdups between failed cand and all other probes following it
                for top,cand,coord in zip(top_probe_list,probe_list[1:],probe_coords_list[1:]):
                    pdups_val = pdups(top,cand,strand_conc_a,strand_conc_b,NUPACK_MODEL)
                    pdups_val_list.append(pdups_val)

                    #removes any following probe cands with pdups of this value or higher
                    if pdups_val >= 0.50:
                        #if true they are removed from the list to survey
                        probe_list.remove(cand)
                        #if true they are added to the skipped list
                        skip_probe_names_list.append(cand)
                        probe_coords_list.remove(coord)
                    else:
                        pdups_forward_val = pdups_forward(top,cand,strand_conc_a,strand_conc_b,NUPACK_MODEL)
                        if pdups_forward_val >= 0.50:
                            probe_list.remove(cand)
                            skip_probe_names_list.append(cand)
                            probe_coords_list.remove(coord)


        t1 = time.time()
        all_run_probes_names_list.append(probe_coords_list[0])

        #pop the head probe being compared from the top of the list
        probe_list.pop(0)
        probe_coords_list.pop(0)

        #records the total run-time for probe in loop
        total_n = t1-t0
        probe_times.append(total_n)

    #make dictionaries of the on target, off target, prop
    on_target_dict = dict(zip(keep_probe_names_list,keep_on_target_list))
    off_target_dict = dict(zip(keep_probe_names_list,keep_off_target_list))
    prop_target_dict = dict(zip(keep_probe_names_list,keep_probe_prop_list))
    probe_run_times_dict = dict(zip(all_run_probes_names_list,probe_times))

    return on_target_dict,off_target_dict,prop_target_dict,probe_run_times_dict,loop_count,keep_probe_names_list,skip_probe_names_list,fail_probe_names_list

##############################################################################

def generate_final_df(probe_df,on_target_dict,off_target_dict,
                      prop_target_dict,probe_run_times_dict,
                      loop_count,keep_probe_names_list,skip_probe_names_list,
                      fail_probe_names_list,o_file,ref_flag):
    """
    Function will make the dictionaries into pandas dataframes where the 
    cols are read as probe, on target, off target, on target pdups prop
    Parameters
    ----------
    probe_df :  dataframe
        DESCRIPTION.
    on_target_dict : dictionary
        probes are key and on target pdups aggregate alignment binding sum
        is the value
    off_target_dict : dictionary
        probes are key and off target pdups aggregate alignment binding sum
        is the value    
    prop_target_dict : dictionary
        probes are key and on target pdups  binding proportion
        is the value
        
    Returns
    -------
    None. Writes output file with probes to keep and relevant info
    """

    #make the three dicts into a dataframe
    on_target_df = pd.DataFrame(on_target_dict.items(),
                                columns=['probe_coords',
                                         'on_target_sum'])
    off_target_df = pd.DataFrame(off_target_dict.items(),
                                 columns=['probe_coords',
                                          'off_target_sum'])
    prop_target_df = pd.DataFrame(prop_target_dict.items(),
                                  columns=['probe_coords',
                                           'on_target_prop'])

    probe_coords_list = probe_df['probe_coords'].tolist()
    probe_list = probe_df['probe'].tolist()

    coords_probe_dict = dict(zip(probe_coords_list,probe_list))

    skip_coords_list = []
    for key,val in coords_probe_dict.items():
        if val in skip_probe_names_list:
            skip_coords_list.append(key)

    #label each type of probe
    label_list = [] 
    for probe in probe_coords_list:
        if probe in keep_probe_names_list:
            label_list.append("Run,kept")
        elif probe in fail_probe_names_list:
            label_list.append("Run,failed")
        elif probe in skip_coords_list:        
            label_list.append("Skipped")
        else:
            label_list.append("Not run")

    for coords in probe_coords_list:
        if coords not in probe_run_times_dict:
            probe_run_times_dict[coords] = 0

    #total loop_count
    total_loop_count = []
    for i in range(len(probe_list)):
        total_loop_count.append(loop_count)

    #create a dict to get the 
    probe_run_times_df = pd.DataFrame(probe_run_times_dict.items(),
                                  columns=['probe_coords',
                                           'run_time'])        

    #then merge the dicts into the main probe df (lengths should match)
    on_off_merged = pd.merge(on_target_df, off_target_df,
                             on="probe_coords")

    pdups_binding_df = pd.merge(on_off_merged, prop_target_df,
                                on="probe_coords")

    #write final df
    keep_probes_df = pd.merge(probe_df, pdups_binding_df, on="probe_coords")
    keep_probes_df.to_csv(o_file, header=False, index=False, sep="\t")

    #write reference file
    repeat_name = probe_df['region'].iloc[0]

    if int(ref_flag) == 1:

        ref_file_dir = "pipeline_output/02_intermediate_files/06_alignment_filter/log/"
        create_dir(ref_file_dir)
        ref_file_name = ref_file_dir + str(repeat_name) + ".txt"
        log_probes_df = pd.merge(probe_df,probe_run_times_df, on="probe_coords")
        log_probes_df['label'] = label_list
        log_probes_df['loop_count'] = total_loop_count

        log_probes_df.to_csv(ref_file_name, header=False, index=False, sep="\t")


##############################################################################

def generate_pairwise_df(probe_seq,probe_coords,bowtie_idx,bowtie_string,
                         strand_conc_a,strand_conc_b,NUPACK_MODEL,
                         bt2_k_val,seed_length,ref_flag):
    """
    Function will take a probe sequence and generate an alignment against
    the entire genome with given bowtie2 string and indices. Then, 
    sam2pairwise is called and sequences are removed of N bases and the 
    RC of the derived alignment to the probe sequence is used to compute
    the NUPACK pdups score. This is then used to compute on target and 
    off target aggregate pdups scores to predict oligo probe binding
    to target sequences.
    
    Parameters
    ----------
    probe_seq : string
        a probe sequence of interest from the filtered probe dataframe
    probe_coords : string
        the coordinates of the probe sequence
    Returns
    -------
    pairwise_df : dataframe
        includes parent seq, derived seq, and pdups score
    """

    #begin looping over each RR to write a tmp file
    with tempfile.TemporaryDirectory() as tmpdir:

        #make temporary file name if ref_flag != 1 to save memory
        #intermediate files saved to demonstrate test functionality
        fastq_filename = tmpdir + '/region.fastq'

        #reads the file to start writing
        fastq = open(fastq_filename,'w')

        #writes the file as a temp fastq
        quals = '~' * len(probe_seq)
        fastq.write('@'+ str(probe_coords) + '\n%s\n+\n%s' %
                        (probe_seq, quals))
        fastq.close()

        #now you want to take the fastq and I think run bt2
        sam_file = tmpdir + '/derived.sam'

        #the value 300000 is = k, or the max number of alignments returned
        sam_file = bt2_call(fastq_filename,sam_file,bt2_k_val,
                            bowtie_idx,bowtie_string,seed_length)

        if int(ref_flag) == 1:
            bam_dir = "pipeline_output/02_intermediate_files/06_alignment_filter/bam/"
            create_dir(bam_dir)
            bam_file = bam_dir + str(probe_coords) + ".bam" 
        else:
            bam_file = tmpdir + '/derived.bam'

        bam_file = samtools_call(sam_file,bam_file)

        #now make the pairwise file
        pairwise_file = tmpdir + '/derived.out'
        s2p = (sam2pairwise_call(bam_file).decode('utf-8'))

        #writes a tmp pairwise file
        with open(pairwise_file,'w') as pf:
            for line in s2p:
                pf.write(line)
            pf.close()

        #write a function to clean the pairwise sequences
        pairwise_df,unique_df = process_pairwise(pairwise_file)

        #for any probes after it, nupack to see if pdups < 0.50
        parent_seq = unique_df["parent"].tolist()
        derived_seq = unique_df["derived"].tolist()

        #the pdups values and the names of the probe sequences
        pdups_vals_list = []
        probe_names_list = []

        #zip the parent seq and the derived seq
        for ps,ds in zip(parent_seq,derived_seq):

            #add names and compute pdups to list
            probe_names_list.append(ps + "_" + ds)

            #compute the pdups values
            pdups_val = pdups(ps,ds,strand_conc_a,strand_conc_b,NUPACK_MODEL)

            #add the values into a list
            pdups_vals_list.append(pdups_val)

        #add names and pdups into zipped dict
        probe_pdups_dict = dict(zip(probe_names_list,pdups_vals_list))

        #make a list for all pdups scores
        all_pdups_list = []

        #now do this for the larger df to append vals
        all_parent_seq = pairwise_df["parent"].tolist()
        all_derived_seq = pairwise_df["derived"].tolist()

        #zip the two lists together and append value to key in dict
        for ps,ds in zip(all_parent_seq,all_derived_seq):
            all_dict_key = ps + "_" + ds
            all_pdups_list.append(probe_pdups_dict[all_dict_key])

        pairwise_df['pdups'] = all_pdups_list

        return pairwise_df

##############################################################################

def create_dir(dir):
   """
   Function creates directories without throwing an error
   """
   if not os.path.exists(dir):
      os.makedirs(dir)
   return dir

##############################################################################

def bt2_call(fq_file,sam_file,k_val,bowtie_idx,bowtie_string,seed_length):
    """
    Function runs bowtie2
    Parameters
    ----------
    fq_file : fastq file 
     fastq file for a particular probe sequence
    sam_file : sam file
        the temp sam file to be generated
    k_val : int
        the value of k is the max number of alignments to be returned
        by bowtie2
    bowtie_idx: path
        the file path to the bt2 idx for a particular genome
    Returns
    -------
    sam_file : sam file
        the temp sam file to be generated
    """

    #use subprocess to call bowtie
    subprocess.call(['bowtie2', '-x', bowtie_idx,
         '-U', str(fq_file),
         '-k',str(k_val),
         bowtie_string,
         '-S', str(sam_file),
         '-L', str(seed_length)])

    return sam_file

##############################################################################

def samtools_call(sam_file,bam_file):
    """
    Function takes the sam file derived and generates a bam file to run
    sam2pairwise
    Parameters
    ----------
    sam_file : sam file
        sam file contains the alignment sequences
    bam_file : name of output bam file
        bam contains the appropriate format to parse out alignment
        sequences
    Returns
    -------
    bam_file : name of output bam file
        bam contains the appropriate format to parse out alignment
        sequences
    """

    #call subprocess to cast sam file into bam file
    subprocess.call(["samtools", "view", "-bS", sam_file],
                    stdout=open(bam_file,'w'))

    return bam_file

##############################################################################
def sam2pairwise_call(bam_file):
    """
    Function calls sam2pairwise to generate derived sequences for each
    recorded alignment from bowtie2 
    Parameters
    ----------
    bam_file : bam file
        contains sequences in the appropriate format
    Returns
    -------
    s2p : sam to pairwise output
        contains probe sequence and derived alignment sequence
    """

    #calls on subprocess to generate sam2pairwise output file
    sv = subprocess.Popen(['samtools', 'view', bam_file], \
    stdout=subprocess.PIPE)

    #allows sam2pairwise output file to be readable
    s2p = (subprocess.check_output('sam2pairwise', stdin=sv.stdout))

    return s2p


##############################################################################
def process_pairwise(pairwise_file):
    """
    Function takes the sam2pairwise output file and parses out the derived
    sequences into a pandas dataframe. Returns a compact version where
    sequences are collapsed into unique pairs as well as the version
    that is not condensed into unique pairs
    Parameters
    ----------
    pairwise_file : sam2pairwise output
        output of running sam2pairiwise
    Returns
    -------
    pairwise_df : dataframe
        the full pairwise dataframe of alignments
    unique_probes_df : dataframe
        the collapsed dataframe with unique pairs of alignments
    """

    #create list to add dictionary of rows
    rows_list = []

    with open(pairwise_file) as f:
        for probe_ID,parent,aligns,reference in itertools.zip_longest(*[f]*4):

            #create dict to store the row
            row_dict = {}
            row_dict.update(probe_ID = probe_ID.split('\t')[0],
                            parent = str(parent.replace('-','').strip('N\n')),
                            derived = str(Seq(reference.replace('-','').
                                              strip('N\n'))),
                            align_chr = probe_ID.split('\t')[2],
                            align_start = probe_ID.split('\t')[3])

            rows_list.append(row_dict)

        #append the row into an appropriate datafrane
        pairwise_df = pd.DataFrame(rows_list, columns = ['probe_ID',
                                        'parent',
                                        'derived',
                                        'align_chr',
                                        'align_start'])

        pairwise_df=pairwise_df[~pairwise_df.parent.str.contains("N")]
        pairwise_df=pairwise_df[~pairwise_df.derived.str.contains("N")]


    #collapse duplicates
    unique_probes_df = pairwise_df.drop_duplicates(subset=['parent',
                                                  'derived'],
                                          keep='first')

    return pairwise_df, unique_probes_df

##############################################################################
def pdups(seq1,seq2,strand_conc_a,strand_conc_b,NUPACK_MODEL):
    """
    Function implements NUPACK by invoking the NUPACK model and computing
    pdups over two sequences of interest, taking the RC of seq2
    Parameters
    ----------
    seq1 : string
        sequence string 1 (parent probe)
    seq2 : string
        sequence string 2 (derived alignment)
    Returns
    -------
    pdups_score : float
        the computed pdups score by nupack
    """

    a = nupack.Strand(seq1, name='a')
    b = nupack.Strand(rev_comp(seq2), name='b')
    t = nupack.Tube({a: strand_conc_a, b: strand_conc_b},
                    complexes=nupack.SetSpec(max_size=2), name='t')

    # calculate partition function and resulting concentrations
    tube_result = nupack.tube_analysis(tubes=[t], model=NUPACK_MODEL)

    # calculate duplex probability
    conc = tube_result.tubes[t].complex_concentrations[nupack.Complex([a, b])]
    pdups_score = conc / strand_conc_b

    return pdups_score

##############################################################################

def pdups_forward(seq1,seq2,strand_conc_a,strand_conc_b,NUPACK_MODEL):
    """
    Function implements NUPACK by invoking the NUPACK model and computing
    pdups over two sequences of interest, taking the RC of seq2
    Parameters
    ----------
    seq1 : string
        sequence string 1 (parent probe)
    seq2 : string
        sequence string 2 (derived alignment)
    Returns
    -------
    pdups_score : float
        the computed pdups score by nupack
    """

    a = nupack.Strand(seq1, name='a')
    b = nupack.Strand(seq2, name='b')
    t = nupack.Tube({a: strand_conc_a, b: strand_conc_b},
                    complexes=nupack.SetSpec(max_size=2), name='t')

    # calculate partition function and resulting concentrations
    tube_result = nupack.tube_analysis(tubes=[t], model=NUPACK_MODEL)

    # calculate duplex probability
    conc = tube_result.tubes[t].complex_concentrations[nupack.Complex([a, b])]
    pdups_score = conc / strand_conc_b

    return pdups_score

##############################################################################

def nupack_sum(pairwise_df,probe_region_dict,ref_flag):
    """
    Function computes the sum of nupack scores that correspond to on target
    and off target binding
    Parameters
    ----------
    pairwise_df :  dataframe
        contains probes, corresponding alignment, derived from sam2pairwise.
    probe_region_dict : dictionary
        contains information about the probe and repeat region in a dict
    Returns
    -------
    total_prop_dict : dictionary
        probe is key and on target binding proportion is value
    total_on_target_dict : dictionary
        probe is key and on target binding sum is value.
    total_off_target_dict : dictionary
        probe is key and off target binding sum is value.
    """

    #make a col that includes the derived end
    dp_lengths=[]
    derived_probe_list = pairwise_df['derived'].tolist()
    for probe in derived_probe_list:
        dp_lengths.append(len(probe))

    #obtains the end coordinates of the length of the derived alignment seq
    derived_end_list =[]
    pairwise_df.align_start = pairwise_df.align_start.astype(int)

    #takes the derived alignment start to a lit
    derived_starts_list = pairwise_df['align_start'].tolist()

    #creates a list for the derived end values
    for dpl,ds in zip(dp_lengths,derived_starts_list):
        derived_end_list.append(dpl+ds)

    #adds the derived ends to a pairwise df
    pairwise_df['align_end'] = derived_end_list

    #repeat coords for the probe are added to a list
    repeat_coords_list = []
    probe_coords_list = pairwise_df['probe_ID'].tolist()

    #obtain the repeat coords for each probe coord
    for pc in probe_coords_list:
        if pc in probe_region_dict:
            repeat_coords_list.append(probe_region_dict[pc])

    #append repeat coords as a col in df
    pairwise_df['repeat_coords'] = repeat_coords_list

    #format columns with the repeat_coords
    pairwise_df[['parent_chrom','region']] = pairwise_df['repeat_coords'].str.split(':',
                expand=True)

    pairwise_df[['r_start','r_end']] = pairwise_df['region'].str.split('-',
                expand=True)

    #cast r columns as ints
    pairwise_df[['r_start','r_end']] = pairwise_df[['r_start',
                'r_end']].astype(int)

    #make dicts to store the on_target_sum, off_target, total
    prop_dict = {}
    on_target_dict = {}
    off_target_dict = {}

    on_total = 0
    off_total = 0

    probe_chrom = probe_coords_list[0].split(":")[0]

    #probe list
    derived_chrom_list = pairwise_df['align_chr'].tolist()
    parent_chrom_list=[probe_chrom for i in range(len(derived_chrom_list))]

    #repeat coords list
    repeat_start_list = pairwise_df['r_start'].tolist()
    repeat_end_list = pairwise_df['r_end'].tolist()

    #derived probe starts and ends
    derived_start_list = pairwise_df['align_start'].tolist()
    derived_end_list = pairwise_df['align_end'].tolist()

    #pdups list
    pdups_list = pairwise_df['pdups'].tolist()

    #run the relevant probe chrom, derived chrom, and starts and stops
    #for the probe region and the derived alignment
    for pc,dc,ds,de,rs,re,pdups in zip(parent_chrom_list,
                                       derived_chrom_list,
                                       derived_start_list,
                                       derived_end_list,
                                       repeat_start_list,
                                       repeat_end_list,pdups_list):

        #checks if the derived alignment coordinates fall within the 
        #region of the repeat coordinates
        if pc == dc and int(ds) >= int(rs) and int(de) <= int(re):
            on_total += pdups
        else:
            off_total += pdups

    #instead of a list this should be a dictionary
    prop_dict[probe_coords_list[0]] = (on_total)/(on_total+off_total)
    on_target_dict[probe_coords_list[0]] = on_total
    off_target_dict[probe_coords_list[0]] = off_total

    if int(ref_flag) == 1:
        bam_dir = "pipeline_output/02_intermediate_files/06_alignment_filter/bam_pdups/"
        create_dir(bam_dir)
        bam_file = bam_dir + str(probe_coords_list[0]) + ".txt"
        save_pairwise_df = pairwise_df.drop(columns=['parent_chrom','align_end','region','repeat_coords','r_start','r_end'])
        save_pairwise_df.to_csv(bam_file, header=False, index=False, sep="\t")

    return prop_dict,on_target_dict,off_target_dict

##############################################################################

def get_bedtools_map(probe_region,probe_coords,genome_bin_file,top_probe_al,thresh,ref_flag):

    with tempfile.TemporaryDirectory() as tmpdir:

        #the file for the bed file overlap
        repeat_filename = tmpdir + '/region.bed'
        repeat_overlap_out = tmpdir + '/overlap_out.bed'
        probe_overlap_out = tmpdir + '/probe_overlap_out.bed'


        #write the repeat contents as a bed file
        repeat_bed = open(repeat_filename, 'w')

        #split the string on chars
        probe_region = probe_region.replace(":","\t")
        probe_region = probe_region.replace("-","\t")

        #write repeat into file format
        repeat_bed.write(str(probe_region))
        repeat_bed.close()

        #subprocess call to get overlap file
        subprocess.call(['bedtools', 'intersect', '-wa', '-wb', '-a',
                         repeat_filename, '-b', genome_bin_file],
                    stdout=open(repeat_overlap_out,'w'))

        #read repeat file as dataframe
        colnames = ["chrom", "start", "stop", "chrom_b","start_b","stop_b"]
        repeat_overlap_out_df = pd.read_csv(repeat_overlap_out,
                                            names=colnames,header=None,
                            sep='\t')


        #the tmp file for the alignments
        alignment_file = tmpdir + '/probe_alignments.tsv'


        #drop cols
        columns = ['probe_ID', 'parent', 'derived', 'pdups', 
                   'repeat_coords', 'parent_chrom', 'region',
                   'r_start', 'r_end']
        
        top_probe_al_coords = top_probe_al.drop(columns,axis=1)

        #write contents of df to file
        top_probe_al_coords.to_csv(alignment_file, header=False,
                                   index=False, sep="\t")

        #subprocess call to get overlap file
        subprocess.call(['bedtools', 'intersect', '-wa', '-wb', '-a',
                         alignment_file, '-b', genome_bin_file],
                    stdout=open(probe_overlap_out,'w'))

        #read probe overlap filer as dataframe
        colnames = ["chrom", "start", "stop", "chrom_b","start_b","stop_b"]
        probe_overlap_out_df = pd.read_csv(probe_overlap_out,
                                           names=colnames,
                                           header=None,sep='\t')

        #format pDups file correctly for map function
        columns = ['align_end','repeat_coords','parent_chrom',
                   'region','r_start','r_end']
        top_probe_al_map = top_probe_al.drop(columns,axis=1)

        top_probe_al_map = top_probe_al_map[["probe_ID","parent","derived",
                "align_chr","align_start","pdups"]]

        #read repeat file as dataframe
        colnames = ["chrom","bin_start","bin_stop"]
        genome_bin_file_df = pd.read_csv(genome_bin_file, names=colnames,
                                         header=None,sep='\t')

        signal_pileup_subset,chrom_summ_df = map_alignments_by_bin(genome_bin_file_df,
                                                                   probe_overlap_out_df,
                                                                   repeat_overlap_out_df,
                                                                   top_probe_al_map,thresh)

        
        return signal_pileup_subset,chrom_summ_df

##############################################################################

def map_alignments_by_bin(chr_track,chr_overlap,repeat_overlap,pairs_pdups,thresh):

    """
    Function takes the genome bin file, the bedtools intersect file, and
    the pairwise file containing pdups computes to process them as 
    dataframes.
    
    Parameters
    ----------
    chr_track_file : file
        file containing the 1Mb bins of the genome
    chr_overlap_bed : file
        bedtools intersect output over alignment coords
    repeat_overlap_bed: file
        bedtools intersect output over repeat region coords
    pDups_scores : file
        file from alignment output
    Returns
    -------
    chr_track : dataframe
        bed file containing the coordinates of 1Mb bins
    chr_overlap : dataframe
        contains the overlaps of bedtools intersect on alignments
    repeat_overlap: dataframe
        contains the overlaps of the bedtools intersect on repeat region
    pairs_pdups : dataframe
        contains the pairwise output
    """

    #label columns for ground file with the bed tracks
    chr_track=chr_track[~chr_track.chrom.str.contains("M")]

    #label columns for matching overlaps
    chr_overlap=chr_overlap[~chr_overlap.chrom.str.contains("M")]
    chr_overlap=chr_overlap[~chr_overlap.chrom_b.str.contains("M")]

    #label columns for matching overlaps
    repeat_overlap=repeat_overlap[~repeat_overlap.chrom.str.contains("M")]
    repeat_overlap=repeat_overlap[~repeat_overlap.chrom_b.str.contains("M")]

    #label columns for data where you have the nupack data
    pairs_pdups=pairs_pdups[~pairs_pdups.derived.str.contains("M")]
    pairs_pdups=pairs_pdups[~pairs_pdups.align_chr.str.contains("M")]

    pairs_pdups = pairs_pdups.drop(['parent', 'derived'], axis = 1)

    derived_coords_list = pairs_pdups['align_chr'].tolist()
    starts_list = pairs_pdups['align_start'].tolist()
    pdups_vals_list = pairs_pdups['pdups'].tolist()

    pairs_pdups_dict = {}
    bin_dict = {}
    bin_coords_list = []

    chrom_list = chr_overlap['chrom'].tolist()
    start_list = chr_overlap['start'].tolist()
    start_b_list = chr_overlap['start_b'].tolist()
    stop_b_list = chr_overlap['stop_b'].tolist()

    for chrom,start,pdups in zip(derived_coords_list,starts_list,pdups_vals_list):
        coord_key = str(chrom) + "_" + str(start)
        pairs_pdups_dict[coord_key]=pdups

    coord_key_list = []
    for chrom,start in zip(chrom_list,start_list):
        coord_key = str(chrom) + "_" + str(start)
        coord_key_list.append(coord_key)

    chr_overlap['coord_key'] = coord_key_list

    pdups_list = []
    for coord in coord_key_list:
        if coord in pairs_pdups_dict:
            pdups_list.append(pairs_pdups_dict[coord])
    
    chr_overlap['pdups'] = pdups_list

    #remove intermediate column
    chr_overlap = chr_overlap.drop(['chrom', 'start', 'stop','coord_key'], axis=1)

    chr_overlap.columns = ['chrom','bin_start','bin_stop','pdups']

    bin_sums = chr_overlap.groupby(['chrom','bin_start','bin_stop'])['pdups'].sum().reset_index()

    #subset items in chr_track not in bin sums

    #drop pdups for now 
    bins_w_vals = bin_sums.drop(['pdups'], axis = 1)

    chr_track['bin_start']=chr_track['bin_start'].astype(int)
    chr_track['bin_stop']=chr_track['bin_stop'].astype(int)
    bins_w_vals['bin_start']=bins_w_vals['bin_start'].astype(int)
    bins_w_vals['bin_stop']=bins_w_vals['bin_stop'].astype(int)

    #subset rows not in bins with vals
    common = bins_w_vals.merge(chr_track,on=['chrom', 'bin_start',
                                             'bin_stop'],how='left')

    not_overlapping = pd.merge(chr_track, common, how='left', indicator=True) \
           .query("_merge == 'left_only'") \
           .drop('_merge',1)

    not_overlapping['pdups'] = 0.0

    #now merge with nupack
    frames = [bin_sums,not_overlapping]

    merged = pd.concat(frames)

    merged = merged.sort_values(by=['chrom','bin_start','bin_stop'])

    #here you should read in the repeat dataframe to get the chrom_start_stops
    chrom_list = repeat_overlap['chrom_b'].tolist()
    start_list = repeat_overlap['start_b'].tolist()
    stop_list = repeat_overlap['stop_b'].tolist()

    #make a dictionary storing location as key lookup
    repeat_dict = {}

    for chrom,start,stop in zip(chrom_list,start_list,stop_list):
        repeat_key = str(chrom) + "_" + str(start) + "_" + str(stop)
        repeat_dict[repeat_key] = 1

    #store values of bin type (target = 1, non = 0)
    bin_type = []

    #make lists to generate key lookups of all bins
    bin_chrom_list = merged['chrom'].tolist()
    bin_start_list = merged['bin_start'].tolist()
    bin_stop_list = merged['bin_stop'].tolist()

    for chrom,start,stop in zip(bin_chrom_list,bin_start_list,bin_stop_list):
        location_key = str(chrom) + "_" + str(start) + "_" + str(stop)
        if location_key in repeat_dict:
            bin_type.append(1)
        else:
            bin_type.append(0)

    #annotates whether bin is considered target or non-target
    merged['bin_type'] = bin_type

    #subset the signal over particular regions of interest
    signal_pileup_subset = merged.loc[merged['pdups'] >= int(thresh)]

    #summarize all signals by chromosome in table form 
    chrom_summ_df = merged.groupby(['chrom'])['pdups'].sum().reset_index()

    return signal_pileup_subset,chrom_summ_df

##############################################################################
                                                                          

def main():

    start_time=time.time()

    userInput = argparse.ArgumentParser(description=\
        '%Requires a filtered probe file based on k-mer occurence'
        'Implements bt2 alignment to identify probes that have redundant'
        'binding and significant off targets in the human genome')

    requiredNamed = userInput.add_argument_group('required arguments')
    requiredNamed.add_argument('-f', '--probe_file', action='store',
                               required=True, help='The filtered probe file'
                               'that contains all sequences and regions')
    requiredNamed.add_argument('-o', '--out_file', action='store',
                               required=True, help='The filtered probe file'
                               'that contains all sequences and regions')
    userInput.add_argument('-r', '--region_threshold', action='store',
                           default=5000, type=int, help='The total'
                           'on-target binding sum to reach by repeat region')
    requiredNamed.add_argument('-b', '--bowtie_index', action='store',
                           required=True, help='The path to the bowtie'
                           'index to run the alignment algorithm')
    requiredNamed.add_argument('-k', '--bt2_max_align', action='store',
                           required=True, default = 300000,
                           help='The max number of alignments to be returned'
                           'by bowtie')
    requiredNamed.add_argument('-l', '--seed_length', action='store',
                           required=True, default = 20,
                           help='Seed length when returning bt2 alignments')
    requiredNamed.add_argument('-t', '--model_temp', action='store',
                           required=True, default = 74.5,
                           help='NUPACK model temp, (C)')
    requiredNamed.add_argument('-pb', '--max_pdups_binding', action='store',
                           required=True, default = 0.60,
                           help='In order for a probe to be added into a'
                           'list to be kept, its pdups value with all'
                           'probes must be below this max pdups binding val')
    requiredNamed.add_argument('-moT', '--min_on_target', action='store',
                           required=True, default = 1000,
                           help='The min on target aggregate pdups binding'
                           'sum to consider a probe')
    requiredNamed.add_argument('-Mr', '--max_probe_return', action='store',
                           required=True, default = 10,
                           help='Either aggregate on target is returned or'
                           'max probe number is returned first.')
    requiredNamed.add_argument('-gb', '--genomic_bin', action = 'store',
                           required = True, help = 'genome binned file')
    requiredNamed.add_argument('-th', '--thresh', action='store',
                               required=True, help='pdups >= to subset')
    requiredNamed.add_argument('-p', '--pdups_p', action='store',
                               required=True, help='pdups prop min')
    requiredNamed.add_argument('-rf', '--ref_flag', action='store',
                               required=True, help='reference flag',default = 0)

    args = userInput.parse_args()
    p_file = args.probe_file
    o_file = args.out_file
    r_thresh = args.region_threshold
    bowtie_idx = args.bowtie_index
    bt2_k_val = args.bt2_max_align
    max_pdups_binding = args.max_pdups_binding
    seed_length = args.seed_length
    model_temp = args.model_temp
    min_on_target = args.min_on_target
    max_probe_return = args.max_probe_return
    genomic_bins = args.genomic_bin
    thresh = args.thresh
    pdups_p = args.pdups_p
    ref_flag = args.ref_flag

    #the bowtie string settings used for running the alignment algorithm
    bowtie_string = "--local -N 1 -R 3 -D 20 -i C,4 --score-min G,1,4"

    # configure nupack model for use
    NUPACK_MODEL = nupack.Model(
        material = 'dna',
        celsius = float(model_temp),
        sodium = 0.39,
        magnesium = 0.0,
        ensemble = 'stacking')

    #configure the strand concentrations for nupack predicted duplex score
    #pDups
    strand_conc_a=1e-6
    strand_conc_b=1e-12


    probe_df = read_probe_filter(p_file)

    print("---%s seconds ---"%(time.time()-start_time))

    on_target_d,off_target_d,prop_target_d,probe_run_times_dict,loop_count,keep_probe_names_list,skip_probe_names_list,fail_probe_names_list = filter_thresh(probe_df,
                                                               strand_conc_a,
                                                               strand_conc_b,
                                                               bowtie_idx,
                                                               r_thresh,
                                                               bowtie_string,
                                                               NUPACK_MODEL,
                                                               bt2_k_val,
                                                               max_pdups_binding,
                                                               seed_length,
                                                               max_probe_return,
                                                               min_on_target,
                                                               genomic_bins,
                                                               thresh,pdups_p,ref_flag)

    print("---%s seconds ---"%(time.time()-start_time))

    generate_final_df(probe_df,on_target_d,off_target_d,prop_target_d,probe_run_times_dict,loop_count,keep_probe_names_list,skip_probe_names_list,fail_probe_names_list,o_file,ref_flag)

    print("---%s seconds ---"%(time.time()-start_time))

if __name__== "__main__":
    main()
```
