## Supplementary Software for "Tigerfish designs oligonucleotide-based *in situ* hybridization probes targeting intervals of highly repetitive DNA at the scale of genomes": check_repeatID.py

```
#import libraries
import time
import pandas as pd
import argparse

def read_test_run_files(chr4_t_file, chrx_t_file):
    
    """
    Function takes test probe files and reads them into
    pandas dataframes.
    
    Parameters
    ----------
    chr4_t_file : file path
        chr4 probe file path from user test output
    chrx_t_file: file path
        chrX probe file path from user test output

    Returns
    -------
    chr4_t_df: dataframe
        chr4 test output file organized as a pandas dataframe
    chrx_t_df: dataframe
        chrX test output file organized as a pandas dataframe
    """    
    
    colnames = ["probe_coords","repeat_coords","imaging_coords","probe", "Tm",
                "r_count", "h_count", "binding_prop", "ranking",
                "on_target","off_target","pdups_prop"]
    
    chr4_t_df = pd.read_csv(chr4_t_file, names=colnames,header=None,
                            sep='\t')

    chrx_t_df = pd.read_csv(chrx_t_file, names=colnames,header=None,
                            sep='\t')
        
    return(chr4_t_df,chrx_t_df)
    
##############################################################################
    
def read_expected_output(chr4_e_file, chrx_e_file):
    
    """
    Function take expected output probe files and reads them into 
    pandas dataframes. 
    
    Parameters
    ----------
    chr4_e_file : file path
        chr4 probe file path from expected output
    chrx_e_file: file path
        chrX probe file path from expected output

    Returns
    -------
    chr4_e_df: dataframe
        chr4 expected output file organized as a pandas dataframe
    chrx_e_df: dataframe
        chrX expected output file organized as a pandas dataframe
    """      
    
    colnames = ["probe_coords","repeat_coords","probe", "Tm",
                "r_count", "h_count", "binding_prop", "ranking",
                "on_target","off_target","pdups_prop"]
    
    chr4_e_df = pd.read_csv(chr4_e_file, names=colnames,header=None,
                            sep='\t')

    chrx_e_df = pd.read_csv(chrx_e_file, names=colnames,header=None,
                            sep='\t')
        
    return(chr4_e_df,chrx_e_df)
    
##############################################################################
    
def compare_file_contents(chr4_t_df, chr4_e_df, chrx_t_df, chrx_e_df):
    
    """
    Function takes test dataframes and expected output dataframes and compares
    probe contents as sets to see if contents in tests and expected outputs 
    match.
    
    Parameters
    ----------
    chr4_t_df: dataframe
        chr4 test output file organized as a pandas dataframe
    chrx_t_df: dataframe
        chrX test output file organized as a pandas dataframe
    chr4_e_df: dataframe
        chr4 expected output file organized as a pandas dataframe
    chrx_e_df: dataframe
        chrX expected output file organized as a pandas dataframe
        
    Returns
    -------
    None. Print statment dictating whether files match.
    
    """    
    
    chr4_t_set = set(chr4_t_df['probe'].tolist())
    chrx_t_set = set(chrx_t_df['probe'].tolist())

    chr4_e_set = set(chr4_e_df['probe'].tolist())
    chrx_e_set = set(chrx_e_df['probe'].tolist())
    
    if (chr4_t_set==chr4_e_set) and (chrx_t_set==chrx_e_set):
        
        print("Test run matches Tigerfish expected output!")
        
    else:
        
        print("Test run does not match expected output.")
    
    
##############################################################################

    
def main():
    
    start_time=time.time()
    
    """Reads input test files from repeat_ID test run output and compares
    expected output files from repeat_ID to validate successul test run."""
    
    #allow user inpir parameters on command line
    
    userInput = argparse.ArgumentParser(description=\
        '%Requires probe file output from Tigerfish test run')
        
    requiredNamed = userInput.add_argument_group('required arguments')
    
    requiredNamed.add_argument('-ft', '--chr4_t_file', action='store', 
                               required=True, 
                               help='chr4 probe output test file')
    requiredNamed.add_argument('-xt', '--chrx_t_file', action='store', 
                               required=True, 
                               help='chrX probe output test file')
    requiredNamed.add_argument('-fe', '--chr4_e_file', action='store', 
                               required=True, 
                               help='chr4 expected probe output file')
    requiredNamed.add_argument('-xe', '--chrx_e_file', action='store', 
                               required=True, 
                               help='chrX expected probe output file')

    # Import user-specified command line values.
    args = userInput.parse_args()
    
    ft = args.chr4_t_file
    xt = args.chrx_t_file
    fe = args.chr4_e_file
    xe = args.chrx_e_file
    
    chr4_t_df, chrx_t_df = read_test_run_files(ft,xt)

    print("---%s seconds ---"%(time.time()-start_time))

    
    chr4_e_df,chrx_e_df = read_expected_output(fe,xe)
    
    print("---%s seconds ---"%(time.time()-start_time))

    compare_file_contents(chr4_t_df, chr4_e_df, chrx_t_df, chrx_e_df)

    print("---%s seconds ---"%(time.time()-start_time))

    print("Done")

if __name__ == '__main__':
    main()
```
