## Supplementary Software for "Tigerfish designs oligonucleotide-based *in situ* hybridization probes targeting intervals of highly repetitive DNA at the scale of genomes": collapse_repeat.py

```
import pandas as pd
import time
import argparse
import os
import re
##############################################################################

def read_thresh_file(file_path):
    
    #read in the file with the appropriate column names
    colnames = ["chrom","start","end","score",'region']
    
    #read as a dataframe
    probe_df = pd.read_csv(file_path, delimiter = '\t',
                           names = colnames)
            
    return probe_df

##############################################################################

def read_align_file(align_path):

    colnames = ['probe_coords','probe','align','chrom','start','pdup']

    align_df = pd.read_csv(align_path,delimiter = '\t', names = colnames)

    return align_df

##############################################################################

def collapse_repeat(probe_df):
 
    #subset true region only
    probe_df=probe_df.loc[probe_df['region'] == 1]
        
    #delete score
    probe_df = probe_df.drop(['region'], axis=1)
    
    #sort ranges
    probe_df=probe_df.sort_values(by=['start', 'end'])
    
    #collapse ranges
    collapsed_df = probe_df.groupby(((probe_df.shift()["end"] - probe_df["start"]) < 0).cumsum()).agg({"start":"min",
                                                                                                       "end":"max","score":'sum'})    
    #if len collapse > 1
    if len(collapsed_df) > 1:
        
        #keep row with largest sum
        collapsed_df=collapsed_df[collapsed_df.score == collapsed_df.score.max()]

    #then append chrom
    chrom_val = (probe_df['chrom'].tolist())[0]
    
    collapsed_df['chrom'] = chrom_val
    
    #sort columns
    collapsed_df=collapsed_df[["chrom","start","end","score"]]
    
    #get name of collapsed
    chrom_name = collapsed_df['chrom'].tolist()
    start_val = collapsed_df['start'].tolist()
    end_val = collapsed_df['end'].tolist()
    
    for c,s,e in zip(chrom_name,start_val,end_val):
        region_coords = str(c) + ":" + str(s) + "-" + str(e)
    
    return region_coords

##############################################################################

def chart_alignment(align_df,repeat_name):

    repeat_name_list = re.split('/|:|-|_', repeat_name)
    chrom_val = repeat_name_list[0]
    start_val = repeat_name_list[1]
    stop_val = repeat_name_list[2]

    #keep rows where chrom_val matches in df
    align_df = align_df.loc[align_df['chrom'] == chrom_val]
    align_df = align_df.loc[align_df['start'] >= int(start_val)]
    align_df = align_df.loc[align_df['start'] < int(stop_val)]
 
    start_list = align_df['start'].tolist()
    
    true_start = (sorted(start_list)[0])

    true_end = (sorted(start_list)[-1])
    
    accurate_region = str(chrom_val) + ":" + str(true_start) + "-" + str(true_end)

    return accurate_region

##############################################################################

def append_repeat(region_coords,probe_file,probe_out_path,region_out_path):
    
    colnames = ["probe_coords","repeat_coords","probe","Tm","r_count",
           "h_count","k_prop","rank","on_t","off_t","on_prop"]

    probe_df = pd.read_csv(probe_file, delimiter = '\t',
                           names = colnames)
 
    #get length of master file containing probes
    region_coords_list = [region_coords for i in range(len(probe_df))]
    
    #add column in
    probe_df['align_region_coords'] = region_coords_list
    
    #sort columns
    probe_df = probe_df[["probe_coords","repeat_coords","align_region_coords",
                         "probe","Tm","r_count","h_count","k_prop","rank",
                         "on_t","off_t","on_prop"]]    
    
    #then write out probe file
    probe_df.to_csv(str(probe_out_path), header=False,index=False, sep="\t") 
        
    regions_df = probe_df.drop(["probe_coords","probe","Tm","r_count",
                                "h_count","k_prop","rank","on_t",
                                "off_t","on_prop"], axis=1)
    
    #then write out region compare file
    regions_df.to_csv(str(region_out_path), header=False,index=False, sep="\t")

##############################################################################


def main():
    
    start_time=time.time()

    userInput = argparse.ArgumentParser(description=\
        '%Requires a probe file containing filtered probes'
        'This splits this file into independent repeat regions to prepare'
        'for parallel alignment as final filter pass')

    requiredNamed = userInput.add_argument_group('required arguments')
    requiredNamed.add_argument('-f', '--file_path', action='store', 
                               required=True, help='The post probe file'
                               'that contains all sequences and regions')
    requiredNamed.add_argument('-a', '--align_path', action='store',
                               required=True, help='The alignment file'
                               'that contains all sequences and regions')
    requiredNamed.add_argument('-p', '--probe_file_path', action='store',
                               required=True, help='The post probe file'
                               'that contains all sequences and regions')
    requiredNamed.add_argument('-po', '--probe_out_path', action='store',
                               required=True, help='The directory where'
                               'the split files will be located by repeat')
    requiredNamed.add_argument('-ro', '--region_out_path', action='store',
                               required=True, help='The directory where'
                               'the split files will be located by repeat')

    args = userInput.parse_args()
    file_path = args.file_path
    probe_file_path = args.probe_file_path
    probe_out_path = args.probe_out_path
    region_out_path = args.region_out_path
    align_path = args.align_path

    probe_cols = ["probe_coords","region_coords", "probe","Tm","r_count","h_count",
                 "probe_binding","rank","on_target","off_target","ont_binding"]
    
    #the summary df
    summ_df = pd.read_csv(probe_file_path,sep='\t',names=probe_cols)

    repeat_name = summ_df['region_coords'].tolist()[0]
    
    repeat_name_list = re.split('/|:|-|_', repeat_name)
    chrom_val = repeat_name_list[0]
    start_val = repeat_name_list[1]
    stop_val = repeat_name_list[2]

    repeat_length = int(stop_val) - int(start_val)

    if repeat_length > 150000:

        range_df = read_thresh_file(file_path)

        print("---%s seconds ---"%(time.time()-start_time))

        region_coords = collapse_repeat(range_df)

        print("---%s seconds ---"%(time.time()-start_time))

        append_repeat(region_coords,probe_file_path,probe_out_path,region_out_path)

        print("done")

    elif repeat_length <= 150000:

        align_df = read_align_file(align_path)

        print("---%s seconds ---"%(time.time()-start_time))
    
        region_coords = chart_alignment(align_df, repeat_name)

        print("---%s seconds ---"%(time.time()-start_time))

        append_repeat(region_coords,probe_file_path,probe_out_path,region_out_path)

        print("---%s seconds ---"%(time.time()-start_time))

        print("done")
    
if __name__== "__main__":
    main()    
```
