## Supplementary Software for "Tigerfish designs oligonucleotide-based *in situ* hybridization probes targeting intervals of highly repetitive DNA at the scale of genomes": design_probes.py

```
#!/usr/bin/env python3
# -*- coding: utf-8 -*-
##############################################################################
"""
Created on Mon Jun 28 10:41:18 2021
@author: Robin Aguilar
Beliveau and Noble Labs
University of Washington | Department of Genome Sciences
"""
##############################################################################
#specific script name
script_name = "design_probes"

#import libraries
import time
import argparse
import subprocess
import numpy as np
import pandas as pd
import refactoredBlockparse as bp

#import biopython libraries
from Bio.SeqUtils import MeltingTemp as mt
from Bio import SeqIO

##############################################################################

def make_fasta_from_bed(bed, region_fa, genome_fa, name):
    """
    This function will run a bedtools process on a genomic fasta provided,
    from a bed file to return a fasta of the bed file regions listed.

    Parameters
    ----------
    bed: bed file
    File containing the genomic coordinates to return a multi-entry fasta.
    These genomic coordinates are the identified repeat regions from the
    repeat identification script.

    region_fa: multi-entry fasta file
    The output fasta file of sequences from the generated bed file from
    the repeat identification script.

    genome_fa: fasta file name
    Reference fasta file to be used to create fasta seqs against repeats.

    Returns
    -------
    genome_fa described above
    """

    if str(genome_fa[-3:]) != ".fa":
        genome_fa = str(genome_fa) + "/" + str(name) + ".fa"

    subprocess.call(['bedtools', 'getfasta', '-fi', genome_fa, '-bed',
                     bed, '-fo', region_fa], stderr=None, shell=False)

##############################################################################

def blockParse_run(region_fa,name,probe_out,min_len,min_temp,max_len,max_temp):
    """
    This function takes the provided fasta seqs derived from the bed file
    and passes each repeat seq into the refactoredBlockParse script from
    Oligominer. Here, probes are written and appended to an output dataframe.
    runSequenceCrawler is run using parameters described in Oligominer for 
    default probe generation. 

    Parameters
    ----------
    region_fa: fasta file
    Contains fasta sequences from derived repeat regions

    name: string
    The header of each fasta repeat becomes a column specifying repeat name
    that each probe was derived from.

    Returns
    -------
    probe_out: tsv file
    The output file name containing all designed probes for all provided
    repeat region fasta sequences.
    """

    name_list = []
    sequence_list = []

    fasta_sequences = list(SeqIO.parse(open(region_fa),'fasta'))
    #you want to parse each fasta sequence as a string and append
    #to a sequence list
    
    for fasta in fasta_sequences:
        name_list.append(fasta.id)
        sequence=(str(fasta.seq))
        sequence_list.append(sequence)
        
    #zip the names (headers of the fasta) and the string sequence into a list,
    #then make into a dict
    zipped_list=zip(name_list,sequence_list)
    dict_name_seq=dict(zipped_list)
    
    #then run each item in the dict into the refactored blockParse script
    #which can now handle multi-lined fastas
    for names,sequences in dict_name_seq.items():
        
        probes=bp.runSequenceCrawler(sequences,names,name, int(min_len), int(max_len), 20, 80, 
                                     mt.DNA_NN3, int(min_temp), int(max_temp),
                                     'AAAAA,TTTTT,CCCCC,GGGGG', 390, 50, 0,
                                     25, 25, None, True ,False, False, False,
                                     False, False,(str(probe_out)))
        
##############################################################################

def main():
    
    start_time=time.time()

    """Reads a jellyfish count file of a given scaffold, a chrom index
    file to account for base location, as well as the path to the 
    chromosome fasta to generate bed files of genomic regions that
    have been flagged as having elevated k-mer counts based on user
    parameters.
    """

    userInput = argparse.ArgumentParser(description=\
        '%Requires a jellyfish count file'
        'and chromosome index count generated by generate_jf_idx'
        'and scaffold fasta file derived from generate_jf_idx.') 
        
    requiredNamed = userInput.add_argument_group('required arguments')
    
    requiredNamed.add_argument('-b', '--bed_name', action='store', 
                               required=True, help='The valid regions'
                               'evaluated by Tigerfish as repetitive')
    requiredNamed.add_argument('-r_o', '--region_out', action='store', 
                               required=True, help='Multi-entry fasta of'
                               'mapped repetitive regions')
    requiredNamed.add_argument('-p_o', '--probes_out', action='store', 
                               required=True, help='Output probes from'
                               'blockparse. File is a dataframe.')
    requiredNamed.add_argument('-g', '--genome_fasta', action='store',
                               required=True, help='Genomic fasta reference'
                               'used to generate region_out file')
    requiredNamed.add_argument('-c', '--chrom_name', action='store',
                               required=True, help='The name of chromosome'
                               'undergoing probe design')
    requiredNamed.add_argument('-l', '--min_len', action='store',
                               required=True, help='min len of probe')
    requiredNamed.add_argument('-L', '--max_len', action='store',
                               required=True, help='max len of probe')
    requiredNamed.add_argument('-t', '--min_temp', action='store',
                               required=True, help='min Tm of probe')
    requiredNamed.add_argument('-T', '--max_temp', action='store',
                               required=True, help='max Tm of probe')
    
    args = userInput.parse_args()
    bed = args.bed_name
    region_fa = args.region_out
    probe_out = args.probes_out
    genome_fa = args.genome_fasta
    name = args.chrom_name
    min_len = args.min_len
    max_len = args.max_len
    min_temp = args.min_temp
    max_temp = args.max_temp
    
    
    make_fasta_from_bed(bed,region_fa,genome_fa,name)

    print("---%s seconds ---"%(time.time()-start_time))

    blockParse_run(region_fa,name,probe_out,min_len,min_temp,max_len,max_temp)
    
    print("---%s seconds ---"%(time.time()-start_time))

    print("Done")
    
    
if __name__ == '__main__':
    main()
```
