## Supplementary Software for "Tigerfish designs oligonucleotide-based *in situ* hybridization probes targeting intervals of highly repetitive DNA at the scale of genomes": finish_summary.py

```
#!/usr/bin/env python3
# -*- coding: utf-8 -*-
##############################################################################
"""
Created on Wed Jun 30 12:32:44 2021
@author: Robin Aguilar
Beliveau and Noble Labs
University of Washington | Department of Genome Sciences
"""
##############################################################################

#import libraries
import argparse
import time
import pandas as pd

##############################################################################

def read_probe_file(p_file):
    """
    Reads the final probe file as a pandas dataframe

    Parameters
    ----------
    probe_file : file
        file containing final probes that have passed and undergone 
        alignment

    Returns
    -------
    probe_df: dataframe
        dataframe containing final probe information

    """
    
    #read in the file with the appropriate column names
    colnames = ["probe_coords","repeat_coords","probe","Tm","r_count",
                "h_count","k_score","k_norm", "on_target_sum",
                "off_target_sum", "on_target_prop"]
    
    probe_df = pd.read_csv(p_file, delimiter = '\t', names = colnames)

    return probe_df

##############################################################################

def summarize_probe_data(probe_df,o_file):
    
    
    if len(probe_df) == 0:
        
        #if there were no probes designed in the region, return that 
        #none of the designed probes satisfied the alignment filter
        #so the file is empty
        

       colnames = ["probe_coords","repeat_coords","probe","Tm","r_count",
                "h_count","k_score","k_norm", "on_target_prop"]

       #drop redundant cols
       summ_df = probe_df.drop(colnames, axis=1)

       #fill missing vals with 0s
       summ_df['probe_count'] = 0
       summ_df['on_target_sum'] = 0.0
       summ_df['off_target_sum'] = 0.0
        
    else:
        
        #implement groupby to parse probes in repeats
        #output summary table that includes region and total probes in each
        #also highlights the total aggregate binding of all regions
        
        summ_df = (probe_df.groupby('repeat_coords')
         .agg({'probe':'count','on_target_sum': 'sum','off_target_sum':'sum'})
         .reset_index()
         .rename(columns={'probe':'probe_count'}))
            
    return summ_df
    
##############################################################################

def write_file(summ_df,o_file):
    
    summ_df.to_csv(o_file, header=True, index=False, sep="\t")

##############################################################################

def main():
    
    start_time=time.time()

    userInput = argparse.ArgumentParser(description=\
        '%Requires a final alignment file generated from Tigerfish'
        'takes all probes generated from that file and summarizes'
        'total probes in each repeat and aggregate on target binding')

    requiredNamed = userInput.add_argument_group('required arguments')
    requiredNamed.add_argument('-f', '--probe_file', action='store',
                               required=True, help='The filtered probe file'
                               'that contains all sequences and regions')
    requiredNamed.add_argument('-o', '--out_file', action='store',
                               required=True, help='The summary file'
                               'aggregating target binding by mapped repeat')
    
    args = userInput.parse_args()
    p_file = args.probe_file
    o_file = args.out_file


    probe_df = read_probe_file(p_file)
    
    print("---%s seconds ---"%(time.time()-start_time))
    
    summ_df = summarize_probe_data(probe_df,o_file)
    
    print("---%s seconds ---"%(time.time()-start_time))

    write_file(summ_df,o_file)
    
    print("---%s seconds ---"%(time.time()-start_time))

    print("done")

    
if __name__== "__main__":
    main()
```
