## Supplementary Software for "Tigerfish designs oligonucleotide-based *in situ* hybridization probes targeting intervals of highly repetitive DNA at the scale of genomes": generate_jf_idx.py

```
#!/usr/bin/env python3
# -*- coding: utf-8 -*-

##############################################################################

# Tigerfish
# generate_jf_idx.py

"""
Created on Wed Jun 23 16:56:01 2021
@author: Robin Aguilar
Beliveau and Noble Labs
University of Washington | Department of Genome Sciences
"""
##############################################################################

#specific script name
scriptName = "generate_jf_idxs"

#import libraries
import time
import argparse
from itertools import islice
from operator import itemgetter, attrgetter
import subprocess
import numpy as np
import pandas as pd
from itertools import groupby
import re

#import biopython libraries
from Bio.Seq import Seq
from Bio import SeqIO

##############################################################################

def find_scaffold(fa_file,chrom,scaffold_fa):
    
    """
    Generates fasta file of desired chromosome
    
    Parameters
    ----------
    fa_file : fasta file
        Multi lined Genome fasta file.
    chrom : string
        chromosome string to subset

    Returns
    -------
    scaffold_fa : fasta file
        fasta file of chromosome that was subset
    """

    #opens the fasta file
    fasta_sequences = SeqIO.parse(open(fa_file),'fasta')
    
    #parses the chromosome of interest from file
    for fasta in fasta_sequences:

        if chrom == fasta.id:

            SeqIO.write(fasta,scaffold_fa,"fasta")

    return scaffold_fa 

##############################################################################

def jf_query(jf_idx,scaffold_fa,jf_out):
    
    """
    Runs jellyfish to generate a query file from the jellyfish index provided

    Parameters
    ----------
    jf_idx : jellyfish index file
        genome wide jellyfish index file
    scaffold_fa : fasta file
        chromosome fasta file generated by find_scaffold()

    Returns
    -------
    jf_out : jellyfish count file
        file that describes k-mer counts in genome
    """
    
    #call subprocess to generate query count of k-mers for given fasta
    query_file=subprocess.call(['jellyfish', 'query', jf_idx, '-s',
                     scaffold_fa, '-o', jf_out], stderr=None, shell=False)
    
    return jf_out

##############################################################################
    
def map_coords(scaffold_fa):
    
    """
    This function will take a the chrom fasta file it and identify where 
    all N and non-N bases are located

    Parameters
    ----------
    scaffold_fa : fasta file
        chromosome fasta file generated by find_scaffold()

    Returns
    -------
    bases_dist_start : list
        list to populate with location of ATCG bases (ints)
    n_bases_start : list
        list to populate with location of ATCG bases (ints)
    """
    
    #lists to populate locations of base types, ATCG or N
    bases_dist_start=[]
    n_bases_start=[]
    
    #open fasta file
    fa_seq = list(SeqIO.parse(open(scaffold_fa),'fasta'))
    
    for fasta in fa_seq:
        sequence=str(fasta.seq).lower()
        #first identified where ATCG bases are located
        
        for match in re.finditer('[atcg]',sequence):
            bases_dist_start.append(int(match.start()))
                    
        #identifies the location of N bases
        for match in re.finditer('[n]', sequence):
            n_bases_start.append(int(match.start()))
    
    return bases_dist_start,n_bases_start

##############################################################################

def group_ranges(bases_dist_start,n_bases_start):
    
    """
    This function will return lists continous ranges of N and non-N bases.

    Parameters
    ----------
    bases_dist_start : list
        list containing location of ATCG bases as ints
    n_bases_start : list
        list containing locatio of N bases (if any) as ints

    Returns
    -------
    tuples of continuous ATCG bases (ranges)
    tuples of continuous N bases (n_ranges)
    """

    ranges = []
    n_ranges=[]
    
    #this collapses the set of the ATCG bases to provide  ranges in a list
    for k, g in groupby(enumerate(bases_dist_start), lambda x: x[1]-x[0]):
        group = list(map(itemgetter(1), g))
        ranges.append(str(group[0]) + "\t" + str(group[-1]+1))

    if n_bases_start:
        #this collapses the set of N bases to provide ranges in a list 
        for k, g in groupby(enumerate(n_bases_start), lambda x: x[1]-x[0]):
            group=list(map(itemgetter(1),g))
            n_ranges.append(str(group[0]) + "\t" + str(group[-1]+1))
        
    return ranges,n_ranges

##############################################################################

def create_df_ranges(ranges,n_ranges,n_bases_start):
    """
    This function is used to call on the compute ranges function and handle
    ranges if N bases are present in the fasta file
    
    Parameters
    ----------
    ranges : tuples
        tuples of continous ranges for ATCG basesw
    n_ranges : tuples
        tuples of continuous ranges for N bases
    n_bases_start : list
        list of start values if N bases are present in genome

    Returns
    -------
    probes_ranges : dataframe
        contacenated ranges that includes N if present in genome, else this
        returns a dataframe with continous coordinates of ATCG bases
    """

    #compute a dataframe for normal regions that are not N
    
    probes_ranges=compute_ranges(ranges,"R")
    
    #compute the dataframe for each respective range if N's are found in the file
    
    if n_bases_start:
        n_ranges_df=compute_ranges(n_ranges,"N")
        
        probes_ranges=merge_ranges(probes_ranges,n_ranges_df)
            
    #return that dataframe
    return probes_ranges
    
##############################################################################

def compute_ranges(base_ranges,label):
    
    """
    Helper function called in create_df_ranges()
    
    Parameters
    ----------
    base_ranges : tuples
        this list contains either the ATCG base list or N base list
    label : string
        "R" represents ATCG bases and "N" represents N bases

    Returns
    -------
    num_ranges : dataframe
        dataframe containing start and end of labeled regions, tab seperated

    """
    
    #make a dataframe of bases
    ranges_v=pd.DataFrame(list(base_ranges), columns=['ranges'])
    
    #split the start and end to be seperate columns
    ranges_v[['start','end']] = ranges_v.ranges.str.split(expand=True)
    num_start=ranges_v['start'].astype(int)
    num_end=ranges_v['end'].astype(int)
    num_ranges = pd.concat([num_start, num_end], axis=1)
    
    #label whether the range is of a normal base "R" or an N, "N"
    num_ranges['type']=str(label)
    num_ranges['size']=num_ranges['end'].astype(int)-num_ranges['start'].astype(int)
    
    return num_ranges

##############################################################################

def merge_ranges(norm_ranges,n_ranges):
    """
    Helper function called in create_df_ranges()
    
    Parameters
    ----------
    norm_ranges : dataframe
        dataframe of ranges for ATCG bases, cols are start, stop, range type
    n_ranges : dataframe
        dataframe of ranges for N bases, cols are start, stop, range type

    Returns
    -------
    reorder_df : dataframe
        merged dataframe that contains concatenated both N and R ranges
        if applicable.
    """
    
    #merges the two types of ranges if applicable
    merged_ranges = pd.concat([norm_ranges, n_ranges])
    merged_ranges.index = range(len(merged_ranges.index))
    
    #sorts the dataframes in ascending order
    sorted_df = merged_ranges.sort_values(by=['start'], ascending=True)
    reorder_df=sorted_df.set_index(np.arange(len(sorted_df.index)))
        
    return reorder_df

##############################################################################

def subtract_kmer_length(probes_ranges,mer_l):
    """
    This function will take each row before each N type and deducts from it's
    end value the length of the kmer, all R regions are then put in a new DF
    
    Parameters
    ----------
    probes_ranges : dataframe
        takes the ordered base ranges in a dataframe

    Returns
    -------
    normal_ranges : dataframe containing only index location values of 
    ATCG bases
    """
    
    #for each row contianing an N
    for index, row in probes_ranges.iloc[1:].iterrows():
        if row['type']=="N":
            prev=probes_ranges.index.get_loc(index-1)
            parse_rows=(probes_ranges.iloc[prev])
            
            #take into account the different, the start of the next k-mer
            probes_ranges.loc[prev,['end']]=parse_rows["end"] - int(mer_l)

    #subset all the R's into a seperate dataframe
    normal_ranges=probes_ranges.loc[(probes_ranges.type == "R")]
    
    return normal_ranges

##############################################################################
    
def generate_index_file(normal_ranges,index_out):
    """
    This function will take the two columns from the normal ranges DF and
    then append these ranges to a list. This list is then written out
    as a file.
    
    Parameters
    ----------
    normal_ranges : dataframe containing the adjusted integer locations of 
    ATCG bases with respect to k-mer value.

    Returns
    -------
    None. Writes file for continuous ATCG base location in the genome.
    """

    #list to store the ranges in the dataframe
    kmer_indices=[]

    #make the dataframes into lists
    start_ranges_list = normal_ranges['start'].tolist()
    end_ranges_list = normal_ranges['end'].tolist()

    #then zip the two lists together
    for start,end in zip(start_ranges_list,end_ranges_list):
        for x in range (start, end+1):
            kmer_indices.append(x)

    #write the indices to file
    with open(index_out,"w") as k_file:
        for i in kmer_indices:
            k_file.write(str(i) + "\n")

##############################################################################


def main():
    
    start_time=time.time()

    """Reads a fasta file and jellyfish genome wide index to generate
    jellyfish count files, scaffold index files,
    and seperated chromosome scaffolds. 
    The jellyfish count file takes the given genome file and identifies
    count occurences of all k-mers within the .fa sequence file.
    The scaffold index file contains the position of each k-mer as
    ints, where N-bases are skipped to account for genomics coords
    accurately.
    The jellyfish genome index is created from a genome wide fasta. Selected
    chromosomes of interested are individually separated into chr.fa
    files."""
    
    #allow user inpir parameters on command line
    
    userInput = argparse.ArgumentParser(description=\
        '%Requires a genome FASTA file as input'
        'And Jellyfish Index file of genome'
        'single-entry or multi-lined FASTA files are supported.')
        
    requiredNamed = userInput.add_argument_group('required arguments')
    
    requiredNamed.add_argument('-f', '--fasta_file', action='store', 
                               required=True, 
                               help='The genomic fasta file')
    
    requiredNamed.add_argument('-j', '--jf_indexfile', action='store',
                               required=True,
                               help='The jf file of a given genome')
    
    requiredNamed.add_argument('-c', '--chr_name', action='store', 
                               required=True,
                               help='Define the scaffold being queried')
    
    requiredNamed.add_argument('-f_o', '--scaffold_fa_out', action='store',
                               required=True,
                               help='scaffold fasta output file')
    
    requiredNamed.add_argument('-j_o', '--jf_out', action='store', 
                               required=True,
                               help='jellyfish query file output')
    
    requiredNamed.add_argument('-i', '--j_index_out', action='store',
                               required=True,
                               help='jellyfish index output file')
    
    requiredNamed.add_argument('-m', '--mer_val', action='store',
                               required=True,
                               help='length of k-mers queried')

    args = userInput.parse_args()
    fa_file = args.fasta_file
    jf_idx = args.jf_indexfile
    chrom= args.chr_name
    scaffold_fa = args.scaffold_fa_out
    jf_out = args.jf_out
    index_out = args.j_index_out
    mer_l = args.mer_val


    scaffold_fa = find_scaffold(fa_file,chrom,scaffold_fa)

    print("---%s seconds ---"%(time.time()-start_time))

    jf_out = jf_query(jf_idx,scaffold_fa,jf_out)
    
    print("---%s seconds ---"%(time.time()-start_time))

    bases_dist_start,n_bases_start = map_coords(scaffold_fa)

    print("---%s seconds ---"%(time.time()-start_time))
    
    ranges,n_ranges=group_ranges(bases_dist_start,n_bases_start)
    
    print("---%s seconds ---"%(time.time()-start_time))
    
    probes_ranges = create_df_ranges(ranges,n_ranges,n_bases_start)

    print("---%s seconds ---"%(time.time()-start_time))

    normal_ranges=subtract_kmer_length(probes_ranges,mer_l)
    
    print("---%s seconds ---"%(time.time()-start_time))

    kmer_indices=generate_index_file(normal_ranges,index_out)
    
    print("---%s seconds ---"%(time.time()-start_time))
    
    print("Done")


if __name__ == '__main__':
    main()
    
```
