## Supplementary Software for "Tigerfish designs oligonucleotide-based *in situ* hybridization probes targeting intervals of highly repetitive DNA at the scale of genomes": get_alignments.py

```
#!/usr/bin/env python3
# -*- coding: utf-8 -*-
##############################################################################
"""
Created on Thu Jul  1 19:09:52 2021
@author: Robin Aguilar
Beliveau and Noble Labs
University of Washington | Department of Genome Sciences
"""
##############################################################################

#import libraries
import pandas as pd
import matplotlib.pyplot as plt
import re
from sklearn import preprocessing
import time
import argparse

##############################################################################

def generate_dfs(chr_track_file,chr_overlap_bed,repeat_overlap_bed,
                 pDups_scores):
    
    """
    Function takes the genome bin file, the bedtools intersect file, and
    the pairwise file containing pdups computes to process them as 
    dataframes.
    
    Parameters
    ----------
    chr_track_file : file
        file containing the 1Mb bins of the genome
    chr_overlap_bed : file
        bedtools intersect output over alignment coords
    repeat_overlap_bed: file
        bedtools intersect output over repeat region coords
    pDups_scores : file
        file from alignment output
    Returns
    -------
    chr_track : dataframe
        bed file containing the coordinates of 1Mb bins
    chr_overlap : dataframe
        contains the overlaps of bedtools intersect on alignments
    repeat_overlap: dataframe
        contains the overlaps of the bedtools intersect on repeat region
    pairs_pdups : dataframe
        contains the pairwise output
    """

    #label columns for ground file with the bed tracks
    colnames = ["chrom","bin_start","bin_stop"]
    chr_track = pd.read_csv(chr_track_file, names=colnames,header=None,
                            sep='\t')
    
    chr_track=chr_track[~chr_track.chrom.str.contains("M")]

    #label columns for matching overlaps
    colnames = ["chrom", "start", "stop", "chrom_b","start_b","stop_b"]
    chr_overlap = pd.read_csv(chr_overlap_bed, names=colnames,header=None,
                          sep='\t')
    
    chr_overlap=chr_overlap[~chr_overlap.chrom.str.contains("M")]
 
    #label columns for matching overlaps
    colnames = ["chrom", "start", "stop", "chrom_b","start_b","stop_b"]
    repeat_overlap = pd.read_csv(repeat_overlap_bed, names=colnames,
                                 header=None, sep='\t')
    
    repeat_overlap=repeat_overlap[~repeat_overlap.chrom.str.contains("M")]

    #label columns for data where you have the nupack data
    colnames = ["align_coords","parent","derived",
                "derived_chrom","align_start","pdups"]

    pairs_pdups = pd.read_csv(pDups_scores, names=colnames,header=None,
                          sep='\t')

    pairs_pdups=pairs_pdups[~pairs_pdups.derived_chrom.str.contains("M")]

    return chr_track,chr_overlap,repeat_overlap,pairs_pdups

##############################################################################

def map_columns(pairs_pdups,chr_overlap):
    """
    Function maps the bedtools intersect overlaps with the pdups scorese
    Parameters
    ----------
    pairs_pdups : dataframe
        contains the pairwise probes with pdups computes
    chr_overlap : dataframe
        contains the genomic overlaps from bedtools intersect
    Returns
    -------
    bin_sums : dataframe
        contains the derived chrom, start, stop, and pdups score
    """

    pairs_pdups = pairs_pdups.drop(['parent', 'derived'], axis = 1)

    derived_chrom_list = pairs_pdups['derived_chrom'].tolist()
    starts_list = pairs_pdups['align_start'].tolist()
    pdups_vals_list = pairs_pdups['pdups'].tolist()

    pairs_pdups_dict = {}
    bin_dict = {}
    bin_coords_list = []

    chrom_list = chr_overlap['chrom'].tolist()
    start_list = chr_overlap['start'].tolist()
    start_b_list = chr_overlap['start_b'].tolist()
    stop_b_list = chr_overlap['stop_b'].tolist()

    for chrom,start,pdups in zip(derived_chrom_list,starts_list,pdups_vals_list):
        coord_key = str(chrom) + "_" + str(start)
        pairs_pdups_dict[coord_key]=pdups

    coord_key_list = []
    for chrom,start in zip(chrom_list,start_list):
        coord_key = str(chrom) + "_" + str(start)
        coord_key_list.append(coord_key)

    chr_overlap['coord_key'] = coord_key_list

    pdups_list = []
    for coord in coord_key_list:
        if coord in pairs_pdups_dict:
            pdups_list.append(pairs_pdups_dict[coord])

    chr_overlap['pdups'] = pdups_list

    chr_overlap = chr_overlap.drop(['chrom', 'start', 'stop','coord_key'], axis=1)

    chr_overlap.columns = ['chrom','bin_start','bin_stop','pdups']

    bin_sums = chr_overlap.groupby(['chrom','bin_start','bin_stop'])['pdups'].sum().reset_index()

    return bin_sums

##############################################################################

def intersect_chr_tracks(bin_sums,chr_track,repeat_overlap):
    """
    Function takes the where probes map to genomic bins and their pdups
    scores and merges this to all other genomic bins. All other bins that
    do not have pdups scores mapping to them receive a 0.
    Parameters
    ----------
    bin_sums : dataframe
        contains chromosome tracks and corresponding pdups scores
    chr_track : dataframe
        contains the genomic chr, start, stop for 1Mb bins
    Returns
    -------
    merged : dataframe
        contains all genomic bins with corresponding mapped scores
    """
   
    #subset items in chr_track not in bin sums
 
    #drop pdups for now 
    bins_w_vals = bin_sums.drop(['pdups'], axis = 1)

    chr_track['bin_start']=chr_track['bin_start'].astype(int)
    chr_track['bin_stop']=chr_track['bin_stop'].astype(int)
    bins_w_vals['bin_start']=bins_w_vals['bin_start'].astype(int)
    bins_w_vals['bin_stop']=bins_w_vals['bin_stop'].astype(int)
    
    #subset rows not in bins with vals
    common = bins_w_vals.merge(chr_track,on=['chrom', 'bin_start','bin_stop'],how='left')
    
    not_overlapping = pd.merge(chr_track, common, how='left', indicator=True) \
           .query("_merge == 'left_only'") \
           .drop('_merge',1)
           
    not_overlapping['pdups'] = 0.0
    
    #now merge with nupack
    frames = [bin_sums,not_overlapping]
    
    merged = pd.concat(frames)
    
    merged = merged.sort_values(by=['chrom','bin_start','bin_stop'])

    #here you should read in the repeat dataframe to get the chrom_start_stops
    chrom_list = repeat_overlap['chrom_b'].tolist()
    start_list = repeat_overlap['start_b'].tolist()
    stop_list = repeat_overlap['stop_b'].tolist()

    #make a dictionary storing location as key lookup
    repeat_dict = {}

    for chrom,start,stop in zip(chrom_list,start_list,stop_list):
        repeat_key = str(chrom) + "_" + str(start) + "_" + str(stop)
        repeat_dict[repeat_key] = 1

    #store values of bin type (target = 1, non = 0)
    bin_type = []

    #make lists to generate key lookups of all bins
    bin_chrom_list = merged['chrom'].tolist()
    bin_start_list = merged['bin_start'].tolist()
    bin_stop_list = merged['bin_stop'].tolist()

    for chrom,start,stop in zip(bin_chrom_list,bin_start_list,bin_stop_list):
        location_key = str(chrom) + "_" + str(start) + "_" + str(stop)
        if location_key in repeat_dict:
            bin_type.append(1)
        else:
            bin_type.append(0)

    #annotates whether bin is considered target or non-target
    merged['bin_type'] = bin_type

    return merged

##############################################################################
#function to return binding summary of all bins >= input_val (~120)

def generate_summary_table(merged,thresh,thresh_summ,chrom_summ):

    #subset the signal over particular regions of interest

    signal_pileup_subset = merged.loc[merged['pdups'] >= float(thresh)]

    #writes row(s) to output
    signal_pileup_subset.to_csv(thresh_summ,index = False,
                               index_label=False,
                               header = None,
                               sep = '\t')

    #summarize all signals by chromosome in table form 
    chrom_summ_df = merged.groupby(['chrom'])['pdups'].sum().reset_index() 

    chrom_summ_df.to_csv(chrom_summ,index = False,
                               index_label=False,
                               header = None,
                               sep = '\t')

##############################################################################

def generate_plot(merged,out_plot):
    """
    Function takes the tracks of genomic regions and maps them
    accordingly based on scaffold.
    Parameters
    ----------
    merged : dataframe
        chrom, start, stop, and pdups sum
    Returns
    -------
    .png of in silico predicted binding of oligo signal
    """
    
    #sort merged
    chrom_list = merged['chrom'].tolist()
    
    #implement sort nicely
    def tryint(s):
        try:
            return int(s)
        except ValueError:
            return s

    def alphanum_key(s):
        return [tryint(c) for c in re.split('([0-9]+)', s)]

    def sort_nicely(l):
        return sorted(l, key=alphanum_key)
    
    chrom_list = sort_nicely(chrom_list)
    
    #keep the first instanc of chrom
    condensed_list = list(dict.fromkeys(chrom_list))
        
    chrom_nums = []
    for i in range(0,len(condensed_list)):
        chrom_nums.append(int(i))
        
    #make a dict that binds the chrom to the num
    chrom_dict = dict(zip(condensed_list, chrom_nums))
    
    merged['chrom_num'] = merged['chrom'].map(chrom_dict)
        
    merged = merged.sort_values(by=['chrom_num','bin_start','bin_stop'])
        
    #scales the values of pdups between 0 - 255 using normalization
    min_max_scaler = preprocessing.MinMaxScaler(feature_range=(0, 255))
    y_transformed = min_max_scaler.fit_transform(merged[['pdups']])
    y_transformed_list = y_transformed.ravel()
    merged['nupack_trans'] = list(y_transformed_list)
    
    merged = merged.drop(['pdups'], axis=1)
    
    #generate linked lists for the chrom bins
    bins_ll=[]
    #values for transformed pdups
    y_vals_trans = []
    #chrom name
    standard_chrom = []
    
    #iterate by chromosome to append y transform values to scaffold
    grouped_probes=merged.groupby('chrom',sort=False)
    for name,group in grouped_probes:
        
        y_list = group['nupack_trans'].tolist()
        y_vals_trans.append(y_list)
        
        bins_list = group['bin_start'].tolist()
        bins_ll.append(bins_list)
        standard_chrom.append(name)

    #plot figure size
    fig = plt.figure(figsize=(24, 8))
    
    #generate figure width
    fig_width_list = []
    for ll in bins_ll:
        fig_width_list.append((ll[-1]))
    
    #scale figure height
    height_list = []
    for i in range(len(fig_width_list)):
        height_list.append(24)
    
    #zip the bins and y vals
    for i,j,chrom,hw in zip(list(range(0,len(bins_ll))),
                  list(range(1,(len(y_vals_trans))+1)),
                  standard_chrom,fig_width_list):
        
        #make each chrom a subplot
        ax = fig.add_subplot(6,4,j)

        #for each figure make plotting params
        params = {"ytick.color" : "w",
          "xtick.color" : "w",
          "axes.labelcolor" : "w",
          "axes.edgecolor" : "w"}
        
        #update plot params
        plt.rcParams.update(params)
        
        #title of each plot is chrom name
        plt.title(str(chrom),color='w',fontsize=14,fontweight='bold')
        
        #set cmap colors
        cmap = plt.cm.Blues
        cmap.set_bad(color='w')

        #remove axes for easy viewing
        ax.spines['left'].set_visible(False)
        ax.spines['top'].set_visible(False)
        ax.spines['right'].set_visible(False)
        ax.get_yaxis().set_visible(False)
        ax.get_xaxis().set_visible(False)
        ax.spines['bottom'].set_linewidth(1.0)
        
        #plot as pileup of signal (histogram)
        plt.hist(bins_ll[i], weights= y_vals_trans[i],color = "magenta")
        ax.set_ylim([0,300])
        ax.set_xlim(0,hw)
                
    #organize subplots
    fig.subplots_adjust(bottom=0.10, right=0.6,top=0.9,hspace = 2.0,
                        wspace = 0.8)
    
    plt.savefig(out_plot,transparent=True,dpi=300)
    
    plt.show()
    
    
##############################################################################

def main():
    
    start_time=time.time()

    userInput = argparse.ArgumentParser(description=\
        'Requires the genome bin file, derived sequence bedtools intersect'
        'and the file containing pariwise sequences')
        
    requiredNamed = userInput.add_argument_group('required arguments')
    requiredNamed.add_argument('-c_t', '--chrom_track', action='store', 
                               required=True, help='The genome file'
                               'containing chrom sizes')
    requiredNamed.add_argument('-c_o', '--chrom_overlaps', action='store',
                               required=True, help='bedtools intersect'
                               'output for alignments')
    requiredNamed.add_argument('-r_o', '--repeat_overlap', action='store',
                               required=True, help='bedtools intersect'
                               'output for repeat regions')
    requiredNamed.add_argument('-p', '--pairwise_pdups', action='store',
                               required=True, help='file with pairwise'
                               'pdups vals')
    requiredNamed.add_argument('-pl', '--out_plot', action='store',
                               required=True, help='output genome plot')
    requiredNamed.add_argument('-t', '--thresh', action='store',
                               required=True, help='pdups >= to subset')
    requiredNamed.add_argument('-t_s', '--thresh_summ', action='store',
                               required=True, help='threshold summ file')
    requiredNamed.add_argument('-c_s', '--chrom_summ', action='store',
                               required=True, help='chrom pdups summ file')

    args = userInput.parse_args()
    chr_track_file = args.chrom_track
    chr_overlap_bed = args.chrom_overlaps
    repeat_overlap_bed = args.repeat_overlap
    pdups_scores = args.pairwise_pdups
    out_plot = args.out_plot
    thresh = args.thresh
    thresh_summ = args.thresh_summ
    chrom_summ = args.chrom_summ
 
    chr_track,chr_overlap,repeat_overlap, pairs_pdups = generate_dfs(chr_track_file,
                                                     chr_overlap_bed,
                                                     repeat_overlap_bed,
                                                     pdups_scores)

    print("---%s seconds ---"%(time.time()-start_time))
    
    bin_sums = map_columns(pairs_pdups,chr_overlap)

    print("---%s seconds ---"%(time.time()-start_time))
    
    merged = intersect_chr_tracks(bin_sums,chr_track,repeat_overlap)

    print("---%s seconds ---"%(time.time()-start_time))

    generate_summary_table(merged,thresh,thresh_summ,chrom_summ)

    print("---%s seconds ---"%(time.time()-start_time))

    generate_plot(merged,out_plot)

    print("---%s seconds ---"%(time.time()-start_time))
    
    print("done")
    
if __name__== "__main__":
    main()
```
