## Supplementary Software for "Tigerfish designs oligonucleotide-based *in situ* hybridization probes targeting intervals of highly repetitive DNA at the scale of genomes": get_region_bed.py

```
#!/usr/bin/env python3
# -*- coding: utf-8 -*-
"""
Created on Tue Mar 15 16:21:37 2022

@author: Robin Aguilar

Purpose: To take the repeat region and split it into a bed file
"""


#import libraries
import time
import pandas as pd
import argparse


##############################################################################

def parse_probe_file(in_file):
    """
    This function reads in the probe file and returns a df of probes
    
    Parameters
    ----------
    in_file : file
        file containing probe_information
    Returns
    -------
    probe_df : dataframe
    """

    colnames = ["probe_coords","repeat_coords","probe","Tm","repeat_count",
                "genome_count","k-binding","k_norm","on_target_sum",
                "off_target_sum","on_target_prop"]
    
    probe_df = pd.read_csv(in_file, delimiter = '\t', names = colnames)
    
    return probe_df

##############################################################################

def get_repeat_bed(probe_df,out_file):

    probe_df = probe_df.drop(columns=["probe_coords",
                                      "probe",
                                      "Tm",
                                      "repeat_count",
                                      "genome_count",
                                      "k-binding",
                                      "k_norm",
                                      "on_target_sum",
                                      "off_target_sum",
                                      "on_target_prop"])
    
    #format columns with the repeat_coords
    probe_df[['parent_chrom','region']] = probe_df['repeat_coords'].str.split(':',
                expand=True)

    probe_df=probe_df.drop_duplicates()

    probe_df[['r_start','r_end']] = probe_df['region'].str.split('-',
                expand=True)

    #cast r columns as ints
    probe_df[['r_start','r_end']] = probe_df[['r_start',
                'r_end']].astype(int)
    

    probe_df = probe_df.drop(columns=["region",
                                      "repeat_coords"])
    
    #writes row(s) to output bed file
    probe_df.to_csv(out_file,index = False,
                               index_label=False,
                               header = None,
                               sep = '\t')
    

##############################################################################

def main():

    start_time=time.time()

    """Reads in a probe file and remove redundant columns that would have 
    been generated during the alignment phase. The output of this can then
    be run using alignments."""
    
    #allow user inpir parameters on command line
    
    userInput = argparse.ArgumentParser(description=\
        '%Requires a probe file')
        
    requiredNamed = userInput.add_argument_group('required arguments')
    
    requiredNamed.add_argument('-i', '--in_file', action='store', 
                               required=True, 
                               help='The input probe file')
    
    requiredNamed.add_argument('-o', '--out_file', action='store', 
                               required=True, 
                               help='a cleaned probe file without redundant'
                               'cols')

    # Import user-specified command line values.
    args = userInput.parse_args()
    in_file = args.in_file
    out_file = args.out_file
    
    
    probe_df = parse_probe_file(in_file)
    
    print("---%s seconds ---"%(time.time()-start_time))
    
    get_repeat_bed(probe_df,out_file)
    
    print("---%s seconds ---"%(time.time()-start_time))

    print("Done")
    

if __name__ == '__main__':
    main()
```
