## Supplementary Software for "Tigerfish designs oligonucleotide-based *in situ* hybridization probes targeting intervals of highly repetitive DNA at the scale of genomes": kmer_filter.py

```
#!/usr/bin/env python3
# -*- coding: utf-8 -*-
##############################################################################

"""
Created on Mon Jun 28 11:09:32 2021
@author: Robin Aguilar
Beliveau and Noble Labs
University of Washington | Department of Genome Sciences
"""
##############################################################################

#specific script name
script_name = "kmer_filter"

#first you should load the libraries
import time
import argparse
import pandas as pd
from itertools import groupby
from collections import Counter
from Bio import SeqIO
from Bio.Seq import reverse_complement as rev_comp

##############################################################################

def read_probe_file(probe_file):
    """
    This function reads the probe file containing designed probes into a 
    pandas dataframe
    
    Parameters
    ----------
    probe_file : file
        file containing probes designed from the design_probes script

    Returns
    -------
    probe_data : dataframe
        dataframe of probes designed with labeled columns
    """
    #read probes csv
    colnames = ["chrom","p_start","p_stop","probe","Tm", "regions"]

    probe_df=pd.read_csv(probe_file, delimiter = '\t',names=colnames)
 
    probe_df = probe_df.drop_duplicates(subset=['probe'], keep='first')

    return probe_df

##############################################################################

def read_fasta_dict(fasta_file):
    """
    This script reads the fasta file generated for all repeat regions that
    underwent probe design and adds them into a dictionary for easy region
    lookup.
    
    Parameters
    ----------
    fasta_file : fasta file
        multi entry fasta contains sequence of each repeat region, generated
        in the design probes script

    Returns
    -------
    seq_dict : dictionary
        contains the probe region header as key and sequence as value
    """

    #parse out the probe file from blockparse
    seq_dict = {rec.id : rec.seq.upper() for rec in SeqIO.parse(fasta_file,
                                                                "fasta")}

    return seq_dict

##############################################################################

def repeat_count(probe_df,seq_dict,jf_file,MERLENGTH,chrom):
    
    """
    This function takes the probes from probes data, and generates indep.
    dictionaries accountaing for the total sum of k-mer count is found 
    within a probes target repeat vs the entire human genome.
    
    Parameters
    ----------
    probe_data : dataframe
        dataframe of designed probes
    seq_dict : dictionary
        dictionary of repeat region and repeat sequence
    jf_file : file
        file containing jellyfish k-mer and k-mer counts

    Returns
    -------
    all_repeat_counts : dictionary
        stores the probe seq as key, and sum of all k-mer counts in repeat
        as val
    all_hg_counts : dictionary
        stores the probe seq as key, and sum of all k-mer counts in genome
        as val
    """    

    #make dictionary ot read jellyfish file into k-mer (key) and count (val)
    jf_dict = {}

    if str(jf_file[-4:]) != ".txt":

        jf_file = str(jf_file) + "/" + str(chrom) + "_jf_temp.txt"

    #read jellyfish file into two lists
    with open(jf_file) as jf:
        rows = (line.split() for line in jf)
        for row in rows:
            jf_dict[row[0]] = int(row[1])

    #make dictionaries for repeat counts and genome counts
    #probe (key) and sum of all k-mer counts (val)
    all_hg_counts = {}
    all_repeat_counts = {}

    #group over each repeat region
    grouped_regions = probe_df.groupby(['regions'],sort = False)
    
    #iterate over each repeat region(name)
    for name,group in grouped_regions:
        
        #make a list of the probes in this group
        probes_list = group["probe"].tolist()
        
        #implement function to split region from the fasta dict(seq_dict)
        #into mers of a given size
        #region_mers is a list
        region_mers = generate_kmers(str(seq_dict[name]),int(MERLENGTH))
        
        #Counter collapses list into dictionary where mer (key) and Counter
        #appends count (val)
        region_mers = Counter(region_mers)
        
        #iterates over probes within a repeat region
        for probe in probes_list:
            
            #creates lists for each probe's k-mer counts in the list
            probe_region_count_list = []
            probe_genome_count_list = []
            
            #implement function to split probe into its k-mers of specified
            #length, returns the mers as a list
            probe_mers = generate_kmers(probe,int(MERLENGTH))
            
            #iterate over each mer in the probe_mers list for each probe
            for mer in probe_mers:
                
                #implement function to generate RC
                #if there are RC matches in repeat or genome, those counts
                #are also considered
                rev_mer = rev_comp(mer)
                
                #then check if mer or mer RC in region
                #append that count value
                if mer in region_mers:
                    probe_region_count_list.append(region_mers[mer])
                if rev_mer in region_mers:
                    probe_region_count_list.append(region_mers[rev_mer])
                    
                #then check if the mer or mer RC is in genome
                #append that count value
                if mer in jf_dict:
                    probe_genome_count_list.append(jf_dict[mer])
                if rev_mer in jf_dict:
                    probe_genome_count_list.append(jf_dict[rev_mer])
            
            #takes the sum of all k-mer counts identified for each probe
            #adds probe (key) and total sum of k-mer counts (val)
            all_repeat_counts[probe] = sum(probe_region_count_list)
            all_hg_counts[probe] = sum(probe_genome_count_list)

    return all_repeat_counts,all_hg_counts

##############################################################################

def generate_kmers(sequence,k_size):
    """
    This function takes a sequence, splits it into mers based on user k_size,
    returns mers as a list

    Parameters
    ----------
    sequence : string
        sequence to be split into mers of specified size
    k_size : int
        the size of k-mer desired

    Returns
    -------
    kmers : list
        list of k-mers depending on specified size

    """

    #make a list to store the kmers
    kmers = []
    
    #compute size of kmer window
    n_kmers = len(sequence) - int(k_size) + 1

    #generate kmer within window
    for i in range(n_kmers):
        kmer = sequence[i:i + int(k_size)]
        kmers.append(kmer)

    return kmers

##############################################################################
    
def append_probe_df(probe_df,all_repeat_counts,all_hg_counts):
    """
    This function takes a probe dataframe, and the dictionary for repeat
    and genome counts and appends the counts according to probes in cols.

    Parameters
    ----------
    probe_df : dataframe 
        dataframe that contains probe information and sequences
    all_repeat_counts : dictionary
        contains probe seq as key and total sum of all k-mer counts found in
        target repeat region as val
    all_hg_counts : dictionary
        contains probe seq as key and total sum of all k-mer counts found in
        human genome.

    Returns
    -------
    probe_df : dataframe
        contains probe infornation with addition of repeat count, hg count, 
        and the k_score which is the r_count/hg_count which determines how
        many of the probes k-mers exclusively target the repeat region of
        interest

    """

    #list to use to append r_count and hg_count as columns in the probe
    #dataframe
    repeat_count_list = []
    hg_count_list = []

    #list of probes
    probe_list = probe_df['probe'].tolist()
    
    #iterate that for probe in the list, append the correct total k-mer sum
    #associated with that particular probe
    for probe in probe_list:
        repeat_count_list.append(all_repeat_counts[probe])
        hg_count_list.append(all_hg_counts[probe])

    #add the lists as columns and cast as ints
    probe_df["r_count"] = repeat_count_list
    probe_df['r_count']=probe_df['r_count'].astype(int)
    probe_df["hg_count"] = hg_count_list
    probe_df['hg_count']=probe_df['hg_count'].astype(int)

    #compute k_score, which is the k-mer binding proportion of the total
    #sum of k-mer counts within the target repeat vs the whole genome
    probe_df['k_score'] = probe_df["r_count"]/probe_df["hg_count"]
    
    #cast k_score column as float
    probe_df['k_score']=probe_df['k_score'].astype(float)

    return probe_df

##############################################################################

def compute_normalized_binding(probe_df, c1_val, c2_val):
    """
    Function groups probes by repeat region and implements weights to rank
    order probes. Higher c1_values indicate that probes are more likely to
    be ranked by higher repeat count, higher c2 values are to implement 
    ranking mostly driven by k-mer binding proportion.
    
    Parameters
    ----------
    probe_df : datframe
        contains probe information in a dataframe

    Returns
    -------
    probe_df : dataframe
        contains probe information with updated ranking scheme based on 
        user input of c1 and c2 values
    """

    #list for the normalized binding value
    norm_val_list = []
    
    #dictionary to contain probe sequence as key and normalized value as val
    all_probe_norm_dict = {}
    
    #est weights, as users params
    c1 = c1_val
    c2 = c2_val

    #iterate by repeat region
    grouped_regions = probe_df.groupby(['regions'],sort = False)
    for name,group in grouped_regions:
        
        #store the value to update and add to the dictionary
        val_list = []
        
        #make a list for the probes
        probe_list = group['probe'].tolist()

        #make a list of the repeat count total and k_binding score
        repeat_counts = group["r_count"].tolist()
        k_binding = group["k_score"].tolist()

        #take the max of the group for the repeat count and k_binding score
        max_r_sum = max(repeat_counts)
        max_k_binding = max(k_binding)

        #do math to produce normalized value
        #in order of probes list
        for rc,kb in zip(repeat_counts,k_binding):
            
            #takes k-mer count in repeat and divides by max k-mer count in
            #entire repeat region and multiplies by the c1 weight.
            #This value is added to the k-mer binding score which is divided
            #by the max k-mer binding score in the repeat region and is
            #multiplied by the c2 weight. The sum of these two normalized
            #values drives ranking of each of these probes within each
            #region
            
            norm_binding_val = (((rc/max_r_sum)*c1) +
                                ((kb/max_k_binding)*c2))
            
            #append this normalized value to the list
            val_list.append(norm_binding_val)

        #zip the probe to the value as a dictionary
        probe_norm_val_dict = dict(zip(probe_list,val_list))
        
        
        #update this dictionary for the whole df
        all_probe_norm_dict.update(probe_norm_val_dict)

    #get a total probe list of normalized values
    probe_list = probe_df['probe'].tolist()
    for probe in probe_list:
        norm_val_list.append(all_probe_norm_dict[probe])

    #append this list to the aggregate df of probe information
    probe_df['norm_vals'] = norm_val_list

    #sort probes by the normalized values
    probe_df = probe_df.sort_values(by=['norm_vals'], ascending=False)

    return probe_df

##############################################################################

def write_file(probe_df, o_path):
    """
    Function takes the dataframe with probe data and writes to a specified
    user output file
    
    Parameters
    ----------
    probe_data : dataframe
        dataframe containing probe information in a ranked order
    o_path : file 
        path for the dataframe to be written as an output file
        

    Returns
    -------
    None. Returns a file with the dataframe information
    """

    #write dataframe to file
    probe_df.to_csv(str(o_path), header= False, index=False, sep="\t")

##############################################################################

def main():
    
    start_time=time.time()

    userInput = argparse.ArgumentParser(description=\
        '%Requires a probe file, repeat region fasta, and jellyfish'
        'k-mer count for a specific scaffold, the output is a filtered file'
        'where probes are ranked in order of normalized binding score val')

    requiredNamed = userInput.add_argument_group('required arguments')
    requiredNamed.add_argument('-p', '--probe_file', action='store',
                               required=True, help='The file probes were'
                               'designed in')
    requiredNamed.add_argument('-j', '--jf_file', action='store', 
                               required=True,help='The kmer count file'
                               'from jellyfish')
    requiredNamed.add_argument('-f','--fasta', action='store',required=True,
                               help = 'The fasta file for all repeat regions')          
    requiredNamed.add_argument('-m', '--merlength', action='store',
                               default=18, type=int, help='The size of'
                               'k-mers, should be same as jf build k-mer'
                               'value; default is 18')
    requiredNamed.add_argument('-o', '--out_path', action='store',
                               required=True, help='Output path of the'
                               'filtered probe file')
    requiredNamed.add_argument('-c1', '--c1_value', action='store',
                               required=True,type=int,help='weight 1 used'
                               'to compute normalized binding score')
    requiredNamed.add_argument('-c2', '--c2_value', action='store', 
                               required=True, type=int,help='weight 2 used'
                               'to compute normalized binding score')
    requiredNamed.add_argument('-c', '--chrom', action='store',
                               required=True,help='scaffold name')

    args = userInput.parse_args()
    
    probe_file=args.probe_file
    jf_file=args.jf_file
    fasta_file=args.fasta
    MERLENGTH=args.merlength
    o_path = args.out_path
    c1_val = args.c1_value
    c2_val = args.c2_value
    chrom = args.chrom


    probe_df = read_probe_file(probe_file)

    print("---%s seconds ---"%(time.time()-start_time))
    
    seq_dict = read_fasta_dict(fasta_file)

    print("---%s seconds ---"%(time.time()-start_time))
    
    all_repeat_counts,all_genome_counts = repeat_count(probe_df,
                                                             seq_dict,
                                                             jf_file,
                                                             MERLENGTH,chrom)

    print("---%s seconds ---"%(time.time()-start_time))
    
    probe_df = append_probe_df(probe_df,all_repeat_counts,
                                 all_genome_counts)

    print("---%s seconds ---"%(time.time()-start_time))

    probe_df = compute_normalized_binding(probe_df, c1_val, c2_val)

    print("---%s seconds ---"%(time.time()-start_time))

    write_file(probe_df, o_path)

    print("---%s seconds ---"%(time.time()-start_time))

if __name__== "__main__":
    main()
```
