## Supplementary Software for "Tigerfish designs oligonucleotide-based *in situ* hybridization probes targeting intervals of highly repetitive DNA at the scale of genomes": make_chromomap.R

```
#!/usr/bin/Rscript

install.packages("chromoMap",repos="http://cran.us.r-project.org")

library(chromoMap) 
library(htmlwidgets)
library(optparse)
library(tidyverse)
library(splitstackshape)

option_list = list(
  make_option(c("-c", "--chrom_sizes"), type="character", default=NULL, 
              help="dataset file name", metavar="character"),
  make_option(c("-r", "--repeat_annotation"), type="character", default=NULL,
              help="dataset file name", metavar="character"),
    make_option(c("-o", "--out"), type="character", default="out.txt", 
              help="output file name [default= %default]", metavar="character")
); 
 
opt_parser = OptionParser(option_list=option_list);
opt = parse_args(opt_parser);

if (is.null(opt$repeat_annotation)){
  print_help(opt_parser)
  stop("At least one argument must be supplied (input file).n", call.=FALSE)
}


#read in probe file
probe_repeat <- read.table(opt$repeat_annotation, 
                           sep = "\t", header=FALSE)

print(probe_repeat)

renamed_regions <- probe_repeat %>% 
  rename(
    chrom = V1,
    start = V2,
    stop = V3
    
  )

renamed_regions$annot <- "tigerfish"
#drop and reorder cols
renamed_regions<-renamed_regions[,c("annot","chrom","start","stop")]
renamed_regions <- as.data.frame(renamed_regions)

print(renamed_regions)

renamed_regions$start[renamed_regions$start == 0] <- 1

print(renamed_regions)

#read in chrom length file
chrom_sizes <- read.table(opt$chrom_sizes, 
                           sep = "\t",header=FALSE)

print(chrom_sizes)

#subset where chrom in regions is row in chrom_sizes

select_chrom <- subset(chrom_sizes, V1 %in% renamed_regions$chrom)

#name columns
colnames(select_chrom)<-c("chrom","stop")

#add start of chrom
select_chrom$start<-1

select_chrom<-select_chrom[,c("chrom","start","stop")]

#generate chromomap
map<-chromoMap(list(select_chrom),list(renamed_regions),
          anno_col = c("magenta"),
          segment_annotation = T,
          data_type = "categorical",
          chr_length = 6,
          chr_width = 25,
          canvas_width = 800,
          label_font = 12,
          chr_color = c("#abdbe3"))

#isave as html
saveWidget(map,opt$out)
```
