## Supplementary Software for "Tigerfish designs oligonucleotide-based *in situ* hybridization probes targeting intervals of highly repetitive DNA at the scale of genomes": make_derived_beds.py

```
#!/usr/bin/env python3
# -*- coding: utf-8 -*-
##############################################################################
"""
Created on Thu Jul  1 18:28:03 2021
@author: Robin Aguilar
Beliveau and Noble Labs
University of Washington | Department of Genome Sciences
"""
##############################################################################

#specific script name
script_name = "make_derived_bed"

#load libraries used
import argparse
import time
import pandas as pd

##############################################################################

def read_probe_file(file_path):
    """
    This function reads the parsed file from sam2pairwise alignments
    
    Parameters
    ----------
    file_path : file
        file containing alignments from sam2pairwise with computed pdups
    Returns
    -------
    probe_data : dataframe
        dataframe of probe alignments and pdups values
    """
    
    colnames = ["probe_coords","probe_seq","derived_seq","chrom","start",
                "pdups"]
    
    probe_df = pd.read_csv(file_path, delimiter = '\t', names = colnames)

    return probe_df

##############################################################################

def generate_bed(probe_df,out_path):
    """
    This function generates a bed file of all alignments provided in probe_df
    
    Parameters
    ----------
    probe_df : dataframe
        contains probe alignments and sequence info
    Returns
    -------
    None. 
        Output file contains bed coordinates of alignments identified.
    """

    #make the start and sequences into lists
    derived_seq_list = probe_df['derived_seq'].tolist()
    derived_start_list = probe_df['start'].tolist()
    
    #make a list for the end values
    end_list = []
    
    #take the length of the sequence and add to start
    for start,seq in zip(derived_start_list,derived_seq_list):
        end_list.append(int(start) + len(seq))

    #make the end the column
    probe_df['end'] = end_list

    #remove additional columns
    columns = ["probe_coords","probe_seq","derived_seq","pdups"]
    probe_df = probe_df.drop(columns,axis=1)
    
    #rearrange cols
    probe_df = probe_df[['chrom','start', 'end']]
    
    #write as output file
    probe_df.to_csv(out_path, header=False, index=False, sep="\t")
        

##############################################################################

def main():
    
    start_time=time.time()

    userInput = argparse.ArgumentParser(description=\
        'Takes pairwise alignment file and generates bed file'
        'from location of derived alignment pairs')
        
    requiredNamed = userInput.add_argument_group('required arguments')
    requiredNamed.add_argument('-f', '--file_path', action='store', 
                               required=True, help='The genome file'
                               'containing chrom sizes')
    requiredNamed.add_argument('-o', '--out_path', action='store',
                               required=True, help='The directory where'
                               'the bin file should be kept')
    
    args = userInput.parse_args()
    file_path = args.file_path
    out_path = args.out_path
    
    probe_df = read_probe_file(file_path)
    print("---%s seconds ---"%(time.time()-start_time))

    generate_bed(probe_df,out_path)
    print("---%s seconds ---"%(time.time()-start_time))
    
    print("done")

if __name__== "__main__":
    main()
```
