## Supplementary Software for "Tigerfish designs oligonucleotide-based *in situ* hybridization probes targeting intervals of highly repetitive DNA at the scale of genomes": probe_mer_filter.py

```
#!/usr/bin/env python3
# -*- coding: utf-8 -*-
##############################################################################
"""
Created on Mon Jun 28 15:47:32 2021
@author: Robin Aguilar
Beliveau and Noble Labs
University of Washington | Department of Genome Sciences
"""
##############################################################################

#specific script name
script_name = "probe_mer_filter"

#load libraries used
import time
import argparse
import pandas as pd
from itertools import groupby

##############################################################################

def read_region(file_path, ENRICH, COPY_NUM):
    """
    
    Function reads in probe file with normalized filter scores that are
    pre-filter. This takes the ENRICH and COPY_NUM values to filter probes.
    
    Parameters
    ----------
    file_path : file
        probe file that contains pre-filter k_score and k_norm values.

    Returns
    -------
    region_df : dataframe
        filtered dataframe that contains the filtered probes based on user
        specified arguments

    """
    
    #read in the file with the appropriate column names
    colnames = ["chrom","p_start","p_end","probe","Tm","region",
                "r_count_total","h_count_total","k_score","k_norm"]
    
    #read as a dataframe
    region_df = pd.read_csv(file_path, delimiter = '\t', names = colnames)
        
    region_df['k_score']=region_df['k_score'].astype(float)

    #filters dataframe based on user parameters
    region_df=region_df[(region_df['k_score'] >= float(ENRICH)) &
                        (region_df['r_count_total'] >= int(COPY_NUM))]
    
    #sorts the pandas dataframe based on descending order
    region_df=region_df.sort_values(by='k_norm', ascending=False)

    return region_df

##############################################################################

def split_mers(region_df,MERLENGTH):
    """
    Implements the generate_kmers function to add a column to the probe
    dataframe that decomposes each probe into it's k-mers (length specified
    by user)
    
    Parameters
    ----------
    region_df : dataframe
        dataframe that contains the probe sequences

    Returns
    -------
    region_df : dataframe
        dataframe that now contains a column that is a list of k-mers
        for the probe sequence
    """

    #make a list of the probes in the dataframe that have been filtered
    probe_list = region_df['probe'].tolist()

    #make a list for the k-mers
    all_mers_list = []
    
    #iterate over each probe in the list
    for probe in probe_list:
        #decompose the probe into k-mers of given length
        mer_list = generate_kmers(probe,MERLENGTH)
        #append list to list
        all_mers_list.append(mer_list)

    #add list of lists to appropriate dataframe rows
    region_df['mers_list'] = all_mers_list

    return region_df

##############################################################################

def generate_kmers(sequence,k_size):
    """
    Function take a string sequence and splits it into k-mers of a specified
    size in the form of a list.

    Parameters
    ----------
    sequence : string
        sequence to be decomposed into k-mers
    k_size : int
        length of k-mers to be decomposed

    Returns
    -------
    kmers : list
        list of k-mers of specified size

    """
    #make a list to store the kmers
    kmers = []

    #compute size of kmer window
    n_kmers = len(sequence) - int(k_size) + 1

    #generate kmer within window
    for i in range(n_kmers):
        kmer = sequence[i:i + int(k_size)]
        kmers.append(kmer)

    return kmers

##############################################################################

def rm_shared_mer_probes(region_df,MER_CUTOFF):
    """
    For all repeat regions that have more than one probe in them, this
    function will take the list of k-mers and add the top ranking probe
    based on k_norm into a set. For all probes within the same repeat region,
    if, it shares at least >= specified mer_cutoff (user specified prop), 
    the probe is removed from the final probe set. This list column of each
    probes k-mers is then dropped.

    Parameters
    ----------
    region_df : dataframe
        contains relevant probes before mer filtering

    Returns
    -------
    region_df : dataframe
        contains relevant probes following filtering based on mer
        proportion

    """
    
    #make a list for probes to be flagged that will be removed
    probe_to_remove = []
    
    #do a groupby over the repeat regions
    grouped_regions=region_df.groupby('region',sort=False)
    
    for name,group in grouped_regions:
        #if there is more than one probe in a given repeat region
        if len(group) > 1:
            
            #gets a list of all k-mers in each probe
            max_mers_list = group['mers_list'].tolist()
    
            #defines top ranking "keep set of 18-mers"
            keep_mers_set = set(max_mers_list[0])
        
            #get a list of all probes
            probes_list = group['probe'].tolist()
            
            for i in range(1,len(max_mers_list)):
                #will keep track of number of 18-mers within a given mers
                #list in the keep set
                
                in_count = 0
                #iterates starting with max_mers_list[1]
                for mer in max_mers_list[i]:
                    #checks if a mer is in the keep set
                    if mer in keep_mers_set:
                        #if so tally count
                        in_count += 1
                    if mer not in keep_mers_set:
                        #if mer is unique add it to the keep set
                        keep_mers_set.add(mer)
        
                #take the lengh of the given max_mer_list[i]
                list_len = len(max_mers_list[i])

                #compute proportion of shared 18-mers
                #if proportion is >= cutoff, then you want to cull
                if float(in_count/list_len) >= MER_CUTOFF:
                    probe_to_remove.append(probes_list[i])
    
    #remove rows of the dataframe that have probes that should be filtered
    region_df = region_df[~region_df['probe'].isin(probe_to_remove)]

    #drop the mers column
    region_df.drop(['mers_list'], axis=1, inplace=True)
            
    return region_df

##############################################################################

def write_file(region_df,out_path):
    """
    This function writes the filtered probe dataframe into a file

    Parameters
    ----------
    region_df : dataframe
        dataframe containing filtered probes

    Returns
    -------
    None. Probe file is generated from dataframe

    """

    region_df.to_csv(str(out_path), header=False, index=False, sep="\t")

##############################################################################

def main():
    
    start_time=time.time()

    userInput = argparse.ArgumentParser(description=\
        '%Requires a probe file containing filter params'
        'Returns filtered file that also accounts for filtering probes'
        'based on sharing a particular proportion of a top ranking probes'
        'k-mers.')

    requiredNamed = userInput.add_argument_group('required arguments')
    requiredNamed.add_argument('-f', '--file_path', action='store', 
                               required=True, help='The prefilter probe file'
                               'that contains all sequences and regions')
    requiredNamed.add_argument('-o', '--out_path', action='store',
                               required=True, help='The filtered probe file'
                               'that contains all sequences and regions')
    userInput.add_argument('-e', '--enrich_score', action='store', 
                           default=0.50, required = True, type=float, 
                           help='The max # of times a given k-mer is found'
                           'in a repeat region/entire HG, default is 0.50')
    userInput.add_argument('-cn', '--copy_num', action='store', default=10, 
                       required = True, type=int, help='The minimum allowed'
                       'number of times a k-mer is identified to be added'
                       'to the final probe list, default is 10')
    userInput.add_argument('-m', '--mer_cutoff', action='store', default=0.95, 
                       required = True, type=float, help='The proportion of'
                       'shared mers used to filter probes')
    userInput.add_argument('-k', '--merlength', action='store', default=18, 
                       required = True, type=int, help='The kmer length')
    
    args = userInput.parse_args()
    file_path = args.file_path
    out_path = args.out_path
    ENRICH=args.enrich_score
    COPY_NUM=args.copy_num
    MER_CUTOFF = args.mer_cutoff
    MERLENGTH = args.merlength


    region_df = read_region(file_path,ENRICH,COPY_NUM)
    print("---%s seconds ---"%(time.time()-start_time))

    region_df = split_mers(region_df,MERLENGTH)
    print("---%s seconds ---"%(time.time()-start_time))

    region_df = rm_shared_mer_probes(region_df,MER_CUTOFF)
    print("---%s seconds ---"%(time.time()-start_time))
        
    write_file(region_df,out_path)
    print("---%s seconds ---"%(time.time()-start_time))
    
    print("done")
    
if __name__== "__main__":
    main()
```
