## Supplementary Software for "Tigerfish designs oligonucleotide-based *in situ* hybridization probes targeting intervals of highly repetitive DNA at the scale of genomes": refactoredBlockparse.py

```
#!/usr/bin/env python

# Specific script name.
scriptName = 'refactoredBlockparse'

# Specify script version.
Version = '1.7'

# Import module for handling input arguments.
import argparse
import numpy as np
import pandas as pd

# Import timeit module and record start time. This provides a rough estimate of
# the wall clock time it takes to run the script.
import timeit

# Import the math module.
import math

# Import Biopython modules.
from Bio.SeqUtils import MeltingTemp as mt
from Bio.Seq import Seq
from Bio.Alphabet import IUPAC
from Bio.SeqUtils import GC
from Bio import SeqIO
from Bio.Alphabet import generic_dna

# Import regex module.
import re

class SequenceCrawler:
    def __init__(self, input_string,fasta_scaffold,chr_name, l, L, gcPercent, GCPercent, nn_table, tm, TM,
                 X, sal, form, sp, conc1, conc2, headerVal, bedVal,
                 OverlapModeVal, verbocity, reportVal, debugVal, metaVal, out_path):
        """Initializes a SequenceCrawler, which is used to efficiently scan a
        large sequence for satisfactory probe sequences."""

        """Change into an input_string"""
        self.input_string = input_string
        self.fasta_scaffold=fasta_scaffold
        self.chr_name=chr_name
        self.out_path=out_path

        self.l = l
        self.L = L
        self.gcPercent = gcPercent
        self.GCPercent = GCPercent

        self.tm = tm
        self.TM = TM
        self.X = X
        self.sal = sal
        self.form = form
        self.sp = sp
        self.conc1 = conc1
        self.conc2 = conc2
        self.headerVal = headerVal
        self.bedVal = bedVal
        self.OverlapModeVal = OverlapModeVal
        self.verbocity = verbocity
        self.reportVal = reportVal
        self.debugVal = debugVal
        self.metaVal = metaVal

        # Build the variables required for efficient melting temperature
        # checking. For melting temperature calculations, the nearest neighbor
        # values are stored as the algorithm crawls along a sequence to improve
        # efficiency.
        self.dH = 0
        self.dS = 1
        self.numGC = -999
        self.currdH = None
        self.currdS = None
        self.currInd = None
        self.currLen = None
        self.hQueue = [0] * L
        self.sQueue = [0] * L
        self.frontH = None
        self.backH = None
        self.frontS = None
        self.backS = None
        self.queueInd = None
        self.noGC = False

        # Declare complementary relationships.
        self.comps = {'A': 'T', 'T': 'A', 'C': 'G', 'G': 'C'}
        self.stackTable = self.reformatTable(nn_table)

        # Build parser for FASTA sequence block.
        """Change this into a generic seq object that expects a DNA string"""
        #for seq_record in Seq(self.input_string, generic_dna):
        #    self.block = str(seq_record).upper()
        #    print(self.block)

        self.block=str(Seq(self.input_string,generic_dna)).upper()
        #print(self.block)


    def reformatTable(self, table):
        """Given a NN table of the format in Bio.SeqUtils.MeltingTemp,
        constructs a dictionary that can handle arbitrary nearest neighbor
        identities without having to compute the complement later."""
        newTable = {}
        for inter in table:
            if inter[2] == '/':
                newTable[inter[0:2]] = table[inter]
                newTable[self.comps[inter[1]] + self.comps[inter[0]]] = table[inter]
            else:
                newTable[inter] = table[inter]
        return newTable


    def getFrontVals(self, letter):
             """Get the energetic contributions based on the beginning of the
             sequence. These are based on the 'init_X/Y' values stored in the
             nearest neighbor table."""
             if letter == 'G' or letter == 'C':
                 return (self.stackTable['init_G/C'][self.dH],self.stackTable['init_G/C'][self.dS])
             elif letter == 'A':
                 return (self.stackTable['init_A/T'][self.dH],self.stackTable['init_A/T'][self.dS])
             return (self.stackTable['init_A/T'][self.dH] + self.stackTable['init_5T/A'][self.dH],self.stackTable['init_A/T'][self.dS] + self.stackTable['init_5T/A'][self.dS])


    def getBackVals(self, letter):
          """Get the energetic contributions based on the end of the sequence."""

          if letter == 'G' or letter == 'C':
              return (self.stackTable['init_G/C'][self.dH],self.stackTable['init_G/C'][self.dS])
          elif letter == 'T':
              return (self.stackTable['init_A/T'][self.dH],self.stackTable['init_A/T'][self.dS])
          return (self.stackTable['init_A/T'][self.dH]+ self.stackTable['init_5T/A'][self.dH],self.stackTable['init_A/T'][self.dS]+ self.stackTable['init_5T/A'][self.dS])


    def resetTmVals(self, startInd, startLen):
          """Update the Tm calculation variables, by repopulating the queue. This
          happens when the crawler jumps ahead by more than a single base.."""

          self.currInd = startInd

          # Initialize values.
          self.numGC = 0
          (self.frontH, self.frontS) = self.getFrontVals(self.block[self.currInd])
          (self.backH, self.backS) = self.getBackVals(self.block[self.currInd + startLen - 1])

          # Iterate through the block and compute the nearest neighbor
          # contributions to deltaH and deltaH.
          for i in range(min(self.L, len(self.block) - self.currInd - 2)):
              neighbors = self.block[self.currInd + i: self.currInd + i + 2]
              if i < startLen and self.block[self.currInd + i] in 'GCgc':
                  self.numGC += 1
              if neighbors in self.stackTable:
                  self.hQueue[i] = self.stackTable[neighbors][self.dH]
                  self.sQueue[i] = self.stackTable[neighbors][self.dS]

          # Sum the nearest neighbor and edge contributions.
          self.currdH = sum(self.hQueue[:startLen - 1])+ self.stackTable['init'][self.dH]+ self.frontH + self.backH
          self.currdS = sum(self.sQueue[:startLen - 1])+ self.stackTable['init'][self.dS]+ self.frontS + self.backS

          # Handle the GC content cases.
          self.noGC = self.numGC == 0
          if self.noGC:
              self.currdH += self.stackTable['init_allA/T'][self.dH]
              self.currdS += self.stackTable['init_allA/T'][self.dS]
          else:
              self.currdH += self.stackTable['init_oneG/C'][self.dH]
              self.currdS += self.stackTable['init_oneG/C'][self.dS]
          self.currLen = startLen
          self.queueInd = 0


    def computeGCDiffs(self, diff):
         """Update the energy based on the change in GC content. Basically,
         considers the cases where previously the block had no GC and now has
         one added or previously there was a G or C and it is no longer in the
         sliding window.."""

         self.numGC += diff
         if self.numGC > 0 and self.noGC:
             # subtract the init for no GC
             self.currdH -= self.stackTable['init_allA/T'][self.dH]
             self.currdS -= self.stackTable['init_allA/T'][self.dS]
             # add the init for one GC
             self.currdH += self.stackTable['init_oneG/C'][self.dH]
             self.currdS += self.stackTable['init_oneG/C'][self.dS]
             self.noGC = False
         elif self.numGC == 0 and not self.noGC:
             # subtract the init for one GC
             self.currdH -= self.stackTable['init_oneG/C'][self.dH]
             self.currdS -= self.stackTable['init_oneG/C'][self.dS]
             # add the init for no GC
             self.currdH += self.stackTable['init_allA/T'][self.dH]
             self.currdS += self.stackTable['init_allA/T'][self.dS]
             self.noGC = True


    def probeTmOpt(self, seq1, ind, i, j):
            """Calculate the melting temperature more efficiently, by not
            recomputing stack sums for every possible oligo. This method is based
            on the Tm_NN function in the Bio.SeqUtils.MeltingTemp library. Logic
            for mismatches and other unnecessary parts have been stripped. This
            algorithm uses a sliding window strategy to keep track of deltaH and 
            deltaS contributions, as well as values based on GC content and the
            identities of the bases on the edges of strands."""

            # If we are just looking at a longer sequence, this will extend the
            # considered energy window.
            if ind == self.currInd and self.currLen != len(seq1):
                # Subtract the value for the previous end
                # Add the new base stacks
                # Add new end value
                (newBackH, newBackS) = self.getBackVals(seq1[-1])
                self.currdH = self.currdH - self.backH + newBackH
                self.currdS = self.currdS - self.backS + newBackS
                (self.backH, self.backS) = (newBackH, newBackS)
                diffGC = 0
                for j in range(self.currLen - 1, len(seq1) - 1):
                    self.currdH += self.hQueue[(self.queueInd + j) % self.L]
                    self.currdS += self.sQueue[(self.queueInd + j) % self.L]
                    if seq1[j] in 'GCgc':
                        diffGC += 1
                if seq1[self.currLen - 1] in 'GCgc':
                    diffGC -= 1
                if seq1[-1] in 'GCgc':
                    diffGC += 1
                self.computeGCDiffs(diffGC)

                self.currLen = len(seq1)

            # If we jumped the window forward too far, all Tm values get reset.
            elif ind - self.currInd >= self.L - self.l or self.currLen < len(seq1) + (ind - self.currInd):
                self.resetTmVals(ind, len(seq1))

            # Here, we have moved forward and need to shorten the front and back.
            elif self.currLen > len(seq1):
                # Subtract the value for the previous start and end.
                # Subtract the first base stack(s) and last base stack(s).
                # Add new start and end values.
                (newFrontH, newFrontS) = self.getFrontVals(seq1[0])
                (newBackH, newBackS) = self.getBackVals(seq1[-1])
                self.currdH = self.currdH - self.backH + newBackH - self.frontH + newFrontH
                self.currdS = self.currdS - self.backS + newBackS - self.frontS + newFrontS
                (self.frontH, self.frontS) = (newFrontH, newFrontS)
                (self.backH, self.backS) = (newBackH, newBackS)

                diffGC = 0
                # Subtract from front.
                for j in range(ind - self.currInd):
                    self.currdH -= self.hQueue[(self.queueInd + j) % self.L]
                    self.currdS -= self.sQueue[(self.queueInd + j) % self.L]
                    if self.block[ind - 1 - j] in 'GCgc':
                        diffGC -= 1

               # Subtract from back.
                for j in range(self.currInd + self.currLen - ind - len(seq1)):
                    self.currdH -= self.hQueue[(self.queueInd + self.currLen - 2 - j) % self.L]
                    self.currdS -= self.sQueue[(self.queueInd + self.currLen - 2 - j) % self.L]
                    if self.block[self.currInd + self.currLen - j - 2] in 'GCgc':
                        diffGC -= 1
                if self.block[self.currInd + self.currLen - 1] in 'GCgc':
                    diffGC -= 1

                if seq1[-1] in 'GCgc':
                    diffGC += 1

                self.queueInd = (self.queueInd + ind - self.currInd) % self.L
                for j in range(len(seq1), self.L):
                    if ind + j + 1 < len(self.block):
                        neighbors = self.block[ind + j] + self.block[ind + j + 1]
                        if neighbors in self.stackTable:
                            self.hQueue[(self.queueInd + j) % self.L] = self.stackTable[neighbors][self.dH]
                            self.sQueue[(self.queueInd + j) % self.L] = self.stackTable[neighbors][self.dS]

                # Adjust GC content count as necessary.
                self.computeGCDiffs(diffGC)

                self.currLen = len(seq1)
                self.currInd = ind

            # Adjust estimate based on salt concentration. Note that this logic
            # corresponds to saltcorr = 5 in the MeltingTemp library.
            concval = (self.conc1 - (self.conc2 / 2.0)) * 1e-9
            saltval = mt.salt_correction(Na=self.sal, K=0, Tris=0, Mg=0, dNTPs=0,method=5, seq=seq1)
            tmval = (1000.0 * self.currdH) /(self.currdS + saltval + (1.987 * math.log(concval))) - 273.15

            # ! return mt.chem_correction(tmval, fmd=self.form)
            approxtmval = float('%0.2f' % tmval)
            return mt.chem_correction(approxtmval, fmd=self.form)


    def tmCheck(self, seq2, ind, i, j):
           """Check if a candidate sequence has a melting temperature within
           range."""
           return float(self.tm) < self.probeTmOpt(seq2, ind, i, j) < float(self.TM)


    def gcCheck(self, seq3):
         """Check whether a candidate sequence has the right GC content."""
         return float(self.gcPercent) <= self.numGC * 100.0 / len(seq3) <= float(self.GCPercent)


    def prohibitCheck(self, seq4):
           """Check for prohibited sequence matches."""
           prohibList = str(self.X).split(',')
           for pro in prohibList:
               if re.search(pro, seq4, re.I) is not None:
                   return False
           return True


    def Ncheckopt(self, seq6):
          """Check for N bases in a sequence, searching from the back."""
          return seq6.rfind('N')


    def seqCheck(self, seq8, i):
          """Aggregate results from the N and prohibited sequences checks."""
          if self.Ncheckopt(seq8) == -1 and self.prohibitCheck(seq8):
              return True
     # Report reasons for failure if desired.
          if self.reportVal or self.debugVal:
              # Report on N-base check first.
              if self.Ncheckopt(seq8) != -1:
                  if self.reportVal:
                      self.reportList.append('Sequence window of %d bases '
                                             'beginning at %d failed due to the '
                                             'presence of an interspersed \'N\' '
                                             'base' \
                                             % (self.l, (self.start + i)))
                  self.N_int_fail_.append(1)
                  if self.debugVal:
                      print('Sequence window of %d bases beginning at %d failed '
                            'due to the presence of an interspersed \'N\' base' \
                            % (self.l, (self.start + i)))

              # Report if failure is due to the presence of prohibited sequences.
              if not self.prohibitCheck(seq8):
                  prohibList = str(self.X).split(',')
                  match_list = []
                  for pro in prohibList:
                      match_group = re.search(pro, seq8, re.I)
                      if match_group:
                          foundSeq = match_group.group(0)
                          match_list.append(foundSeq)
                          format_match = ', '.join('%s' % x for x in match_list)
                  if self.reportVal:
                      self.reportList.append('Sequence window of %d bases '
                                             'beginning at %d failed due to the '
                                             'presence of prohibited sequence(s) '
                                             '%s' \
                                             % (self.l, (self.start + i),
                                                format_match))
                      self.prohib_fail.append(1)
                  if self.debugVal:
                      print('Sequence window of %d bases beginning at %d failed '
                            'due to the presence of prohibited sequence(s) %s' \
                            % (self.l, (self.start + i), format_match))

    def probeCheck(self, seq5, ind, i, j):
         """Checks a probe properties based on the current sliding window."""
         # First check for N bases and prohibited sequences
         # in case the sequence window has been extended
         # and now includes them.
         # Next check Tm, % G+C
         # NOTE: Because of the variable setup, the tmCheck MUST come before the
         # gcCheck for this to work properly.
         if self.Ncheckopt(seq5) == -1 and self.prohibitCheck(seq5) and self.tmCheck(seq5, ind, i, j) and self.gcCheck(seq5):
             return True

         # Report reasons for failure if desired.
         if self.reportVal or self.debugVal:
             # Report on N-base check first
             if self.Ncheckopt(seq5) != -1:
                 if self.reportVal:
                     self.reportList.append('Sequence window of %d bases '
                                            'beginning at %d failed due to the '
                                            'presence of an interspersed \'N\' '
                                            'base' \
                                            % (self.l, (self.start + i)))
                     self.N_int_fail.append(1)
                 if self.debugVal:
                     print('Sequence window of %d bases beginning at %d '
                           'failed due to the presence of an interspersed \'N\' '
                           'base' % (self.l, (self.start + i)))

             # Report if failure is due to the presence of prohibited sequences.
             if not self.prohibitCheck(seq5):
                 prohibList = str(self.X).split(',')
                 match_list = []
                 for pro in prohibList:
                     match_group = re.search(pro, seq5, re.I)
                     if match_group:
                         foundSeq = match_group.group(0)
                         match_list.append(foundSeq)
                         format_match = ', '.join('%s' % x for x in match_list)
                 if self.reportVal:
                     self.reportList.append('Sequence window of %d bases '
                                            'beginning at %d failed due to the '
                                            'presence of prohibited sequence(s) '
                                            '%s' \
                                            % (self.l, (self.start + i),
                                               format_match))
                     self.prohib_fail.append(1)
                 if self.debugVal:
                     print('Sequence window of %d bases beginning at %d failed '
                           'due to the presence of prohibited sequence(s) %s' \
                           % (self.l, (self.start + i), format_match))

           # Report if Tm too low/high.
             if not self.tmCheck(seq5, ind, i, j):
                 if self.probeTmOpt(seq5, ind, i, j) < self.tm:
                     if self.reportVal:
                         self.reportList.append('Sequence window of %d bases '
                                                'beginning at %d failed due to '
                                                'Tm of %0.2f being below the '
                                                'allowed range of %d-%d' \
                                                 % ((self.l + j),
                                                    (self.start + i),
                                                    self.probeTmOpt(seq5, ind,
                                                                    i, j),
                                                    self.tm, self.TM))
                         self.Tm_fail_low.append(1)
                     if self.debugVal:
                         print('Sequence window of %d bases beginning at %d '
                               'failed due to Tm of %0.2f being below the '
                               'allowed range of %d-%d' \
                               % ((self.l + j), (self.start + i),
                                  self.probeTmOpt(seq5, ind, i, j),
                                  self.tm, self.TM))
                 if self.probeTmOpt(seq5, ind, i, j) > self.TM:
                     if self.reportVal:
                         self.reportList.append('Sequence window of %d bases '
                                                'beginning at %d failed due to '
                                                'Tm of %0.2f being above the '
                                                'allowed range of %d-%d' \
                                                % ((self.l + j),
                                                   (self.start + i),
                                                   self.probeTmOpt(seq5, ind,
                                                                   i, j),
                                                   self.tm, self.TM))
                         self.Tm_fail_high.append(1)
                     if self.debugVal:
                         print('Sequence window of %d bases beginning at %d '
                               'failed due to Tm of %0.2f being above the '
                               'allowed range of %d-%d' \
                               % ((self.l + j), (self.start + i),
                                  self.probeTmOpt(seq5, ind, i, j),
                                  self.tm, self.TM))

           # Report if %G+C too low/high.
             if not self.gcCheck(seq5):
                 if (self.numGC * 100.0 / len(seq5)) < self.gcPercent:
                     if self.reportVal:
                         self.reportList.append('Sequence window of %d bases '
                                                'beginning at %d failed due to '
                                                '%%G+C of %0.2f being below the '
                                                'allowed range of %d-%d' \
                                                % ((self.l + j),
                                                   (self.start + i),
                                                   (self.numGC * 100.0 \
                                                    / len(seq5)), \
                                                   self.gcPercent,
                                                   self.GCPercent))
                         self.gc_fail_low.append(1)
                     if self.debugVal:
                         print('Sequence window of %d bases beginning at %d '
                               
                               % ((self.l + j), (self.start + i), \
                                  (self.numGC * 100.0 / len(seq5)), \
                                  self.gcPercent, self.GCPercent))
                 if (self.numGC * 100.0 / len(seq5)) > self.GCPercent:
                     if self.reportVal:
                         self.reportList.append('Sequence window of %d bases '
                                                'beginning at %d failed due to '
                                                '%%G+C of %0.2f being below the '
                                                'allowed range of %d-%d' \
                                                % ((self.l + j),
                                                   (self.start + i), \
                                                   (self.numGC * 100.0 \
                                                    / len(seq5)), \
                                                   self.gcPercent,
                                                   self.GCPercent))
                         self.gc_fail_high.append(1)
                     if self.debugVal:
                         print('Sequence window of %d bases beginning at %d '
                               'failed due to %%G+C of %0.2f being below the '
                               'allowed range of %d-%d' \
                               % ((self.l + j), (self.start + i),
                                  (self.numGC * 100.0 / len(seq5)), \
                                  self.gcPercent, self.GCPercent))


    def BedprobeTm(self, seq7):
         """Tm calculation function for use with .bed output."""
         bedTmVal = float(('%0.2f' % mt.Tm_NN(seq7, Na=self.sal,
                                              dnac1=self.conc1,
                                              dnac2=self.conc2)))
         bed_fcorrected = ('%0.2f' % mt.chem_correction(bedTmVal, fmd=self.form))
         return bed_fcorrected

    def run(self):
         """Runs the crawler through the given block sequence to identify probes
         within the FASTA file satisfying the given constraints."""
         """
         Not needed because this modified version takes in a string and not a file object
         """
         # Parse out FASTA coordinate, scaffold info.
         if self.headerVal is None:
             headerParse = (self.fasta_scaffold).split(':')
             #print(headerParse)
             chrom=headerParse[0]
             if len(headerParse) != 1:
                 #chrom = headerParse.split('>')[1].split('\n')[0]
                 #print(chrom)
                 self.start = 1
                 stop = len(self.block)
             elif 'range=' in headerParse:
                 chrom = headerParse.split('=')[1].split(':')[0]
                 #print(chrom)
                 self.start = int(str(headerParse).split(':')[1].split('-')[0])
                 stop = str(headerParse).split('-')[1].split(' ')[0]

             else:
                 chrom = 'chrom'
                 self.start = 1
                 stop = len(self.block)
         else:
             chrom = self.headerVal.split(':')[0]
             #print(chrom)
             self.start = int(str(self.headerVal).split(':')[1].split('-')[0])
             stop = str(self.headerVal).split(':')[1].split('-')[1]
             #print(self.start)

         # Make lists to hold Report info if desired.
         if self.reportVal:
             self.reportList = []
             self.N_int_fail = []
             self.N_block_fail = []
             self.prohib_fail = []
             self.Tm_fail_low = []
             self.Tm_fail_high = []
             self.gc_fail_low = []
             self.gc_fail_high = []

         # Determine the size range the probe sequence can vary over.
         sizeRange = int(self.L) - int(self.l) + 1

         # Determine size of sequence block to mine.
         blockLen = len(self.block)

        # Make a list to store candidate probe coordinates and sequences.
         cands = []

         previousend = 0
         i = 0

         # Skip to first sequence without an unknown base.
         ncheckval = self.Ncheckopt(self.block[i:i + self.l])
         while ncheckval != -1:
             i += ncheckval + 1
             ncheckval = self.Ncheckopt(self.block[i:i + self.l])
             if self.reportVal:
                 self.reportList.append('Skipping %d base window %d-%d because '
                                        'it contains only \'N\' bases' \
                                        % (self.l, (self.start + i - self.l),
                                           (self.start + i - 1)))
                 self.N_block_fail.append(1)
             if self.debugVal:
                 print('Skipping %d base window %d-%d because it contains only '
                       '\'N\' bases' \
                       % (self.l, (self.start + i - self.l),
                          (self.start + i - 1)))
         self.resetTmVals(i, self.l)

         # Iterate over input sequence, vetting candidate probe sequences.
         while i < int(blockLen) - int(self.l):
             # Print status to terminal.
             #if i % 100000 == 0:
             #    print ('%d of %d' % (i, blockLen))

             # Find next sequence without an unknown base.
             ncheckval = self.Ncheckopt(self.block[i:i + self.l])
             while ncheckval != -1:
                 i += ncheckval + 1
                 ncheckval = self.Ncheckopt(self.block[i:i + self.l])
                 if self.reportVal:
                     self.reportList.append('Skipping %d base window %d-%d '
                                            'because it contains only \'N\' '
                                            'bases' \
                                            % (self.l, (self.start + i - self.l),
                                               (self.start + i - 1)))
                     self.N_block_fail.append(1)
                 if self.debugVal:
                     print('Skipping %d base window %d-%d because it contains '
                           'only \'N\' bases' \
                           % (self.l, (self.start + i - self.l),
                              (self.start + i - 1)))
             if self.seqCheck(self.block[i:i + self.l], i):

                 # Search for a sequence that starts at this index and satisfies
                 # all probe constraints.
                 j = 0
                 while i + j + self.l < int(blockLen) and j < sizeRange and not self.probeCheck(self.block[i:i + j + self.l],i, i, j):
                     j += 1

               # If a candidate sequence was found, then store it and write
                 # success to terminal if requested.
                 if not (i + j + self.l >= int(blockLen) or j >= sizeRange):
                     startPos = self.start + i
                     cands.append((str(startPos), str(startPos + j + self.l - 1),
                                   str(self.block[i:i + j + self.l])))
                     if self.verbocity:
                         print ('Picking a candidate probe of %d bases starting '
                                'at base %d' % (self.l + j, startPos))
                     if self.reportVal:
                         self.reportList.append('Picking a candidate probe of '
                                           '%d bases starting at base %d' \
                                           % (self.l + j, startPos))
                     if self.debugVal:
                         print('Picking a candidate probe of %d bases starting '
                               'at base %d' % (self.l + j, startPos))
                     previousend = i + j + self.l - 1

                 # Update the next index to search from. Probes must be
                 # non-overlapping.
                 if self.OverlapModeVal:
                     i += 1
                 else:
                     i = max(i + 1, previousend + 1) + self.sp
             else:
                 i += 1

         outList=[]

         for i, (start, end, seq) in enumerate(cands):
                 outList.append('%s\t%s\t%s\t%s\t%s\t%s' % (chrom, start, end, seq,
                                                        self.BedprobeTm(seq),self.fasta_scaffold))

         output_df = pd.DataFrame([sub.split("\t") for sub in outList])

         with open(str(self.out_path), 'a+') as f:
                 output_df.to_csv(f, sep='\t',header=None, index=False)

         # Write meta information to a .txt file if desired.
         if self.metaVal:
             metaText = open('%s_blockParse_meta.txt' % chrom, 'w')
             if probeNum == 0:
                 metaText.write('%s\t%s\t%d\t No candidate probes discovered' % (self.inputFile, Version, probeNum))
                 metaText.close()
             else:
                 metaText.write('%s\t%s\t%d\t%0.2f\t%0.2f' % (self.inputFile, Version, probeNum,
                                   probeWindow, probeDensity))
                 metaText.close()

        # If desired, create report file.
         if self.reportVal:
             self.N_int_failCount = len(self.N_int_fail)
             self.N_block_failCount = len(self.N_block_fail)
             self.prohib_failCount = len(self.prohib_fail)
             self.Tm_fail_lowCount = len(self.Tm_fail_low)
             self.Tm_fail_highCount = len(self.Tm_fail_high)
             self.gc_fail_lowCount = len(self.gc_fail_low)
             self.gc_fail_highCount = len(self.gc_fail_high)
             windowCount = (self.N_int_failCount + self.N_block_failCount + self.prohib_failCount + self.Tm_fail_lowCount + self.Tm_fail_highCount + self.gc_fail_lowCount+ self.gc_fail_highCount + probeNum)
             reportOut = open('%s_blockParse_log.txt' % chrom, 'w')
             self.reportList.insert(0, 'Results produced by %s %s' % (scriptName, Version))
             if probeNum == 0:
                 self.reportList.insert(1, 'No candidate probes discovered')
             else:
                 self.reportList.insert(1, '%d candidate probes identified in %0.2f '
                                      'kb yielding %0.2f candidates/kb'
                                      % (probeNum, probeWindow, probeDensity))
             self.reportList.insert(2, 'Note: only the first failure encountered is '
                                  'reported. The order of checks is \'N\' bases '
                                  '> prohib. sequences > Tm > %G+C')
             self.reportList.insert(3, '-' * 100)
             self.reportList.insert(4, '%d of %d / %0.4f%% of sequence windows '
                                  'examined resulted in candidate probes'
                                  % (probeNum, windowCount, \
                                     float(probeNum) / float(windowCount) * 100))
             self.reportList.insert(5, '%d of %d / %0.4f%% of sequence windows '
                                  'examined were skipped due to interspersed '
                                  '\'N\' bases' \
                                  % (self.N_int_failCount, windowCount,
                                     float(self.N_int_failCount) \
                                     / float(windowCount) * 100))
             self.reportList.insert(6, '%d of %d / %0.4f%% of sequence windows '
                                  'examined were skipped because they '
                                  'exclusively contained \'N\' bases' \
                                  % (self.N_block_failCount, windowCount,
                                     float(self.N_block_failCount) \
                                     / float(windowCount) * 100))
             self.reportList.insert(7, '%d of %d / %0.4f%% of sequence windows '
                                  'examined failed because they contained '
                                  'prohibited sequences' \
                                  % (self.prohib_failCount, windowCount,
                                     float(self.prohib_failCount) \
                                     / float(windowCount) * 100))
             self.reportList.insert(8, '%d of %d / %0.4f%% of sequence windows '
                                  'examined failed because the Tm was below %d' \
                                  % (self.Tm_fail_lowCount, windowCount,
                                     float(self.Tm_fail_lowCount) \
                                     / float(windowCount) * 100, self.tm))
             self.reportList.insert(9, '%d of %d / %0.4f%% of sequence windows '
                                  'examined failed because the Tm was above %d' \
                                  % (self.Tm_fail_highCount, windowCount,
                                     float(self.Tm_fail_highCount) \
                                     / float(windowCount) * 100, self.TM))
             self.reportList.insert(10, '%d of %d / %0.4f%% of sequence windows '
                                  'examined failed because the %%G+C was below '
                                  '%d' \
                                  % (self.gc_fail_lowCount, windowCount,
                                     float(self.gc_fail_lowCount) \
                                     / float(windowCount) * 100, self.gcPercent))
             self.reportList.insert(11, '%d of %d / %0.4f%% of sequence windows '
                                  'examined failed because the %%G+C was above '
                                  '%d' \
                                  % (self.gc_fail_highCount, windowCount,
                                     float(self.gc_fail_highCount) \
                                     / float(windowCount) * 100, self.GCPercent))
             self.reportList.insert(12, '-' * 100)
             reportOut.write('\n'.join(self.reportList))
             reportOut.close()


def runSequenceCrawler(input_string,fasta_scaffold,chr_name, l, L, gcPercent, GCPercent, nn_table, tm, TM, X, sal, form, sp, conc1, conc2, headerVal, bedVal,OverlapModeVal, verbocity, reportVal, debugVal, metaVal, out_path):
    """Creates and runs a SequenceCrawler instance."""

    sc = SequenceCrawler(input_string,fasta_scaffold,chr_name, l, L, gcPercent, GCPercent, nn_table, tm,
                           TM, X, sal, form, sp, conc1, conc2, headerVal, bedVal,
                           OverlapModeVal, verbocity, reportVal, debugVal,metaVal, out_path)
    sc.run()

def main():
    """Runs the crawler through the given block sequence to identify probes within the FASTA file satisfying the constraints provided through the commandline interface."""
    startTime = timeit.default_timer()
    # Allow user to input parameters on command line.
    userInput = argparse.ArgumentParser(description='%s version %s. Requires a FASTA file as input. Currently, only '
        'single-entry FASTA files are supported.  Returns a .fastq file, which '
        'can be inputted into short read alignment programs. Optionally, a '
        '.bed file can be outputted instead if \'-b\' is flagged. Tm values '
        'are corrected for [Na+] and [formamide].' % (scriptName, Version))
    requiredNamed = userInput.add_argument_group('required arguments')
    requiredNamed.add_argument('-f', '--file', action='store', required=True,
                                   help='The FASTA file to find probes in')
    userInput.add_argument('-l', '--minLength', action='store', default=36,
                               type=int,
                               help='The minimum allowed probe length; default is '
                                    '36')
    requiredNamed.add_argument('-chr', '--chr_name', action='store', required=True,
                                   help='The chromosome name')
    userInput.add_argument('-L', '--maxLength', action='store', default=41,
                               type=int,
                               help='The maximum allowed probe length, default is '
                                    '41')
    userInput.add_argument('-g', '--min_GC', action='store', default=20,
                               type=int,
                               help='The minimum allowed percent G + C, default is '
                                    '20')
    userInput.add_argument('-G', '--max_GC', action='store', default=80,
                               type=int,
                               help='The maximum allowed percent  G + C, default '
                                    'is 80')
    userInput.add_argument('-t', '--min_Tm', action='store', default=42,
                               type=int,
                               help='The minimum allowed Tm, default is 42')
    userInput.add_argument('-T', '--max_Tm', action='store', default=47,
                               type=int,
                               help='The maximum allowed Tm, default is 47')
    userInput.add_argument('-X', '--prohibitedSeqs', action='store',
                               default='AAAAA,TTTTT,CCCCC,GGGGG', type=str,
                               help='Prohibited sequence list (separated by commas '
                                    'with no spaces), default is '
                                    '\'AAAAA,TTTTT,CCCCC,GGGGG\'')
    userInput.add_argument('-s', '--salt', action='store', default=390,
                               type=int,
                               help='The mM Na+ concentration, default is 390')
    userInput.add_argument('-F', '--formamide', action='store', default=50,
                               type=float,
                               help='The percent formamide being used, default is '
                                    '50')
    userInput.add_argument('-S', '--Spacing', action='store', default=0,
                               type=int,
                               help='The minimum spacing between adjacent probes, '
                                    'default is 0 bases')
    userInput.add_argument('-c', '--dnac1', action='store', default=25,
                               type=float,
                               help='Concentration of higher concentration strand '
                                    '[nM] -typically the probe- to use for '
                                    'thermodynamic calculations. Default is 25')
    userInput.add_argument('-C', '--dnac2', action='store', default=25,
                               type=float,
                               help='Concentration of lower concentration strand '
                                    '[nM] -typically the target- to use for '
                                    'thermodynamic calculations. Default is 25')
    userInput.add_argument('-n', '--nn_table', action='store',
                               default='DNA_NN3',
                               type=str,
                               help='The nearest neighbor table of thermodynamic '
                                    'parameters to be used. See options in '
                                    'Bio.SeqUtils.MeltingTemp. Default is DNA_NN3')
    userInput.add_argument('-H', '--header', action='store', type=str,
                               help='Allows the use of a custom header in the '
                                    'format chr:start-stop. E.g. '
                                    '\'chr2:12500-13500\'')
    userInput.add_argument('-b', '--bed', action='store_true', default=False,
                               help='Output a .bed file of candidate probes '
                                    'instead of a .fastq file.')
    userInput.add_argument('-O', '--OverlapMode', action='store_true',
                               default=False,
                               help='Turn on Overlap Mode, which returns all '
                                    'possible candidate probes in a block of '
                                    'sequence including overlaps. Off by default. '
                                    'Note, if selecting this option, the '
                                    '-S/--Spacing value will be ignored')
    userInput.add_argument('-v', '--verbose', action='store_true',
                               default=False,
                               help='Turn on verbose mode to have probe mining'
                                    'progress print to Terminal. Off by default')
    userInput.add_argument('-R', '--Report', action='store_true', default=False,
                               help='Write a Report file detailing the results of '
                                    'each window of sequence considered by the '
                                    'script. The first set of lines give the '
                                    'occurrence of each possible failure mode for '
                                    'quick reference. Off by default. Note, '
                                    'selecting this option will slow the script '
                                    'considerably')
    userInput.add_argument('-D', '--Debug', action='store_true', default=False,
                               help='The same as -Report, but prints info to '
                                    'terminal instead of writing a log file. Off '
                                    'by default')
    userInput.add_argument('-M', '--Meta', action='store_true', default=False,
                               help='Write a text file containing meta information '
                                    'Off by default. Reports input file <tab> '
                                    'estimated runtime <tab> blockParse version '
                                    '<tab> candidates discovered <tab> span in kb '
                                    'covered by candidate probes <tab> candidate '
                                    'probes per kb')
    userInput.add_argument('-o', '--output', action='store', default=None,
                               type=str, help='Specify the stem of the output '
                                              'filename')

    # Import user-specified command line values.
    args = userInput.parse_args()
    l = args.minLength
    L = args.maxLength
    gcPercent = args.min_GC
    GCPercent = args.max_GC
    tm = args.min_Tm
    TM = args.max_Tm
    X = args.prohibitedSeqs
    sal = args.salt
    form = args.formamide
    sp = args.Spacing
    concA = args.dnac1
    concB = args.dnac2
    headerVal = args.header
    bedVal = args.bed
    OverlapModeVal = args.OverlapMode
    verbocity = args.verbose
    reportVal = args.Report
    debugVal = args.Debug
    metaVal = args.Meta
    out_path = args.output

    # Assign concentration variables based on magnitude.
    if args.dnac1 >= args.dnac2:
        conc1 = args.dnac1
        conc2 = args.dnac2

    else:
        conc1 = args.dnac2
        conc2 = args.dnac1

    # Retrieve the stack table. Note that this may give users the opportunity
    # to execute arbitrary code, so better security measures should be employed
    # if this code is ever hosted online.
    exec ('nn_table = mt.%s' % args.nn_table)
        
    runSequenceCrawler(input_string,fasta_scaffold, l, L, gcPercent, GCPercent, nn_table, tm, TM,
                           X, sal, form, sp, conc1, conc2, headerVal, bedVal,
                           OverlapModeVal, verbocity, reportVal, debugVal, metaVal, out_path)

    # Print wall-clock runtime to terminal.
    print ('Program took %f seconds' % (timeit.default_timer() - startTime))

if __name__== "__main__":
    main()
```
