## Supplementary Software for "Tigerfish designs oligonucleotide-based *in situ* hybridization probes targeting intervals of highly repetitive DNA at the scale of genomes": repeat_ID.py

```
#!/usr/bin/env python3
# -*- coding: utf-8 -*-

##############################################################################

"""
Created on Fri Jun 25 12:40:14 2021
@author: Robin Aguilar
Beliveau and Noble Labs
University of Washington | Department of Genome Sciences
"""
##############################################################################

#specific script name
script_name = "repeat_ID"

#import libraries
import time
import csv
import argparse
from collections import defaultdict
from itertools import islice
from operator import itemgetter, attrgetter
import subprocess
import numpy as np
import pandas as pd
import glob
import os
from Bio.Seq import Seq
from Bio import SeqIO
from itertools import groupby
from Bio.Alphabet import generic_dna, generic_protein
from collections import OrderedDict
import collections
from collections import Counter
import itertools
import re

##############################################################################

def open_index_file(index_file,chrom):
    """
    This function will take the two columns from the normal ranges DF and then 
    append these ranges to a list. This list is then written out as a file    

    Parameters
    ----------
    index_file : file
        file containing the index location of ATCG bases in the
        genome
        
    Returns
    -------
    kmer_indices : list
        file is read into a list
    """
    
    if str(index_file[-4:]) != ".txt":

        index_file = str(index_file) + "/" + str(chrom) + "_index.txt"

    #open index file and add to list
    with open(index_file, "r") as idx_f:
        kmer_indices = [line.rstrip('\n') for line in idx_f]

    if len(kmer_indices) == 0:
        print("Jellyfish count file provided not valid. Exiting...")

    return kmer_indices


##############################################################################

def generate_kmer_count_lists(jf_count,chrom):
    """
    This function will store the values from the jf query file into a list for 
    the k_mers and the corresponding count
    
    Parameters
    ----------
    jf_count : file
        file derived from jellyfish count providing count of k-mers in whole
        genome

    Returns
    -------
    k_mer : list
        list containing specific k-mers
    count : list
        list containing k-mer count values
        
    """
    
    k_mer=[]
    count=[]
    
    if str(jf_count[-4:]) != ".txt":

        jf_count = str(jf_count) + "/" + str(chrom) + "_jf_temp.txt"

    with open(jf_count, "r") as jf:
        for line in jf:
            k_mer.append(line.replace(" ","\t").split()[0])
            count.append(line.replace(" ","\t").split()[1])

    return k_mer,count

##############################################################################

def check_threshold(count,THRESHOLD):
    """
    This function will generate a binary list based on whether the count value 
    passes the defined threshold
    
    Parameters
    ----------
    count : list
        list containing k-mer count values
    THRESHOLD : user argument (int)
        min val of k-mer count to be flagged as enrchiched

    Returns
    -------
    pass_thresh_list : list
        binary list where (1) means k-mer count value is >= threshold,
        else assigned 0
    """
    
    pass_thresh_list=[]
    
    for i in count:
        if int(i)>=THRESHOLD:
            pass_thresh_list.append(1)
        else:
            pass_thresh_list.append(0)

    return pass_thresh_list

##############################################################################

def convolve_successes(pass_thresh_list,WINDOW,COMPOSITION):
    """
    The purpose of this function is to convolve over the array in the size of 
    the window you define, then generate a dataframe of all passing ranges   

    Parameters
    ----------
    pass_thresh_list : list
        binary list where (1) means k-mer count value is >= threshold,
        else assigned 0
    WINDOW : user argument (int)
        the length of the window to be searched in the pass_thresh_list
    COMPOSITION : user argument (float)
        the percentage of k-mers in the list that must be flagged as >=
        composition score

    Returns
    -------
    pass_w : tuple
        Windows of sequence to map repetitive region locations.

    """
    #convert list of binary k-mer count vals into array
    iter_data = np.array(pass_thresh_list)

    #iterate through array over length of specific WINDOW
    iter_vals_convolve=np.convolve(iter_data,np.ones(WINDOW,dtype=int),
                                   'valid')

    #list of values in sum values in sliding window casted as float
    iter_list=[(float(element)) for element in iter_vals_convolve]

    #this is the sum of all the counts of the k-mers
    #the most important column here is the row_span_start,
    #this will tell you which k-mer it is
    #this casts to dataframe
    w_sum=pd.DataFrame(list(iter_list),columns=['iter_sum'])

    #the length of the list represents the start values
    w_sum['sliding_win_start']=np.arange(len(w_sum))

    #the location of the end of each sliding window
    w_sum['sliding_win_end']=w_sum['sliding_win_start']+WINDOW-1
    
    #these are all the k-mers that pass the range you specified
    #note that these are not their true indices in sequence 
    #(basically which number k-mer in order they appear)
    pass_w = w_sum.loc[(w_sum['iter_sum']/WINDOW>=COMPOSITION)]

    return pass_w

##############################################################################

def obtain_repeat_indices(pass_w):
    """
    This function will collapse the dataframe into continuous ranges, and then 
    append those to a seperate dataframe, then stored as seperate lists
    
    Parameters
    ----------
    pass_w : dataframe
        subset of the dataframe containing only the rows that pass the 
        composition score from surveying the sliding window

    Returns
    -------
    indices_to_parse : dataframe
        
    """
    
    #lists for the continuous ranges of start and end values that will
    #be used to identify the repeat regions
    min_ranges=[]
    max_ranges=[]

    start_index=[]
    end_index=[]
    
    for k,g in pass_w.groupby(pass_w['sliding_win_start'] -
                              np.arange(pass_w.shape[0])):
            min_ranges.append(min(g['sliding_win_start']))
            max_ranges.append(max(g['sliding_win_end']))

    #add these two values into a dataframe where we can scan the
    #indices of jellyfish count
    indices_to_parse = pd.DataFrame(list(zip(min_ranges, max_ranges)),
                                    columns =['start_index_range',
                                              'end_index_range'])

    start_index = indices_to_parse['start_index_range'].tolist()
    end_index = indices_to_parse['end_index_range'].tolist()

    return indices_to_parse

##############################################################################

def find_repeats(indices_to_parse,kmer_indices,mer_length):
    """
    This function will refer to the indices to parse and append to a list, 
    then at that index, the start val in kmer_indices is appended, this is 
    also done for end sequences.
    
    Parameters
    ----------
    indices_to_parse : datafrane
        contains continous start and end coordinates
    kmer_indices : list
        contains the index derived from jellyfish files
        
    Returns
    -------
    kmer_start_seq : list
        list containing start coords of k-mer
    kmer_end_seq : list
        list containing ned coords of k-mer
    index_start : list
        list of where start indices were noted from parsing sliding window
    index_end : list
        list of where end indices were noted from parsing sliding window

    """

    #list of indices of all valid ATCG bases that represent k-mer location
    kmer_start=[]
    kmer_end=[]

    #index of regions found as valid in k-mer sliding window
    index_start=[]
    index_end=[]
    
    for item in indices_to_parse['start_index_range']:
        index_start.append(int(item))
        start=kmer_indices[item]
        kmer_start.append(int(start))

    repeat_starts_df=pd.DataFrame(list(zip(kmer_start,index_start)),
                                  columns=['r_start','r_index_start'])

    for item in indices_to_parse['end_index_range']:
        index_end.append(int(item))
        end=kmer_indices[item]
        kmer_end.append(int(int(end) + int(mer_length)))
     
    repeat_ends_df=pd.DataFrame(list(zip(kmer_end,index_end)),
                                columns=['r_end','r_index_end'])

    repeat_indices=pd.concat([repeat_starts_df, repeat_ends_df], axis=1)

    return kmer_start,kmer_end,index_start,index_end

##############################################################################

def nucleotide_range(kmer_start,kmer_end,index_start,index_end,bed_file,chrom):
    """
    This function will take the start and end lists and then map this back 
    to the regions in sequence
    
    Parameters
    ----------
    kmer_start : list
        location of start of k-mer
    kmer_end : list
        location of end of k-mer
    index_start : list
        start index of k-mer sliding window
    index_end : list
        end index of k-mer sliding window
    bed_file : file
        path and name of output bed file
    chrom : user arg string
        given scaffold to report bed file

    Returns
    -------
    None. The output bed file is written at specified path

    """

    #zip the two lists together and this should get you the start and end 
    #nucleotides of where that 18mer pattern was
    seq_range = pd.DataFrame(list(zip(kmer_start, kmer_end)), 
                                    columns =['r_start', 'r_end'])

    seq_index_range = pd.DataFrame(list(zip(index_start,index_end)),
                                   columns =['r_index_start','r_index_end'])

    #let's add a column to the dataframe that will contain the
    #chromosome information
    seq_range['chr']=chrom
    seq_index_range['chr']=chrom

    #let's now merge by the chromosome to collapse overlapping regions
    collapsed_nucleotide_range=seq_range.groupby((seq_range.r_end.shift() - 
                                                  seq_range.r_start).lt(0).cumsum()).agg(
        {'r_start': 'first','chr': 'first', 'r_end': 'last'})

    collapsed_nucleotide_range=collapsed_nucleotide_range[["chr","r_start","r_end"]]

    collapsed_nucleotide_range.to_csv(bed_file, header=None, index=None, sep='\t')


##############################################################################

def main():
    
    start_time=time.time()
    
    """Reads a jellyfish count file of a given scaffold, a chrom index
    file to account for base location, as well as the path to the 
    chromosome fasta to generate bed files of genomic regions that
    have been flagged as having elevated k-mer counts based on user
    parameters.
    """
    
    userInput = argparse.ArgumentParser(description=\
        '%Requires a jellyfish count file'
        'and chromosome index count generated by generate_jf_idx'
        'and scaffold fasta file derived from generate_jf_idx.') 
        
    requiredNamed = userInput.add_argument_group('required arguments')
        
    requiredNamed.add_argument('-j', '--jf_count', action='store',
                               required=True, help='The jf file of a'
                               'given genome')
    requiredNamed.add_argument('-i', '--index_file', action='store',
                                   required=True, help='The index file of'
                                   'a given genome')
    requiredNamed.add_argument('-chr', '--chr_name', action='store',
                                   required=True,
                               help='Define the scaffold being queried')
    requiredNamed.add_argument('-st','--start',action='store',
                                   required=True, help='The start sequence of'
                                   'the fasta region if not starting at' 
                                   'beginning of scaffold; default is ' '0')
    requiredNamed.add_argument('-w', '--window_length', action='store', 
                                   default=3000,type=int, help='The length of'
                                   'the scanning window for kmer enriched' 
                                   'regions; default is 3000')
    requiredNamed.add_argument('-t', '--threshold', action='store', 
                               default=10, type=int, help='The minimum number'
                               'of counts that defines a kmer enriched'
                               'region; default is 10')
    requiredNamed.add_argument('-c', '--composition_score', action='store',
                               default=0.5,type=float,help='The minimum'
                               'percentage of kmers that pass threshold' 
                               'in window; default is 0.5')
    requiredNamed.add_argument('-o_b', '--bed_file', action='store',
                               required=True, help='Used to generate fasta'
                               'file of kmer rich regions; default is')
    requiredNamed.add_argument('-m', '--mer_length', action='store',
                               required=True, help='Size of k-mers used')

    #Import user-specified command line values.
    args = userInput.parse_args()
    index_file = args.index_file
    jf_count = args.jf_count
    chrom= args.chr_name
    WINDOW = args.window_length
    THRESHOLD = args.threshold
    COMPOSITION = args.composition_score
    START=args.start
    bed_file = args.bed_file
    mer_length = args.mer_length

    #a list for the regions and scaffold sequences 
    name_list=[]
    sequence_list=[]
    zipped_list=[]
    bases_dist_start=[]

    #the names of output files to be generated
    
    kmer_indices=open_index_file(index_file,chrom)
    
    print("---%s seconds ---"%(time.time()-start_time))

    k_mer,count=generate_kmer_count_lists(jf_count,chrom)

    print("---%s seconds ---"%(time.time()-start_time))

    pass_thresh_list=check_threshold(count,THRESHOLD)

    print("---%s seconds ---"%(time.time()-start_time))

    pass_w=convolve_successes(pass_thresh_list,WINDOW,COMPOSITION)

    print("---%s seconds ---"%(time.time()-start_time))

    indices_to_parse=obtain_repeat_indices(pass_w)

    print("---%s seconds ---"%(time.time()-start_time))

    kmer_start,kmer_end,index_start,index_end=find_repeats(indices_to_parse,
                                                           kmer_indices,
                                                           mer_length)

    print("---%s seconds ---"%(time.time()-start_time))

    nucleotide_range(kmer_start,kmer_end,index_start,index_end,bed_file,chrom)

    print("---%s seconds ---"%(time.time()-start_time))
    
    print("Done")


    
if __name__ == '__main__':
    main()
```
