## Supplementary Software for "Tigerfish designs oligonucleotide-based *in situ* hybridization probes targeting intervals of highly repetitive DNA at the scale of genomes": split_bed.py

```
#!/usr/bin/env python3
# -*- coding: utf-8 -*-

##############################################################################

"""
Created on Fri Jun 25 09:54:15 2021
@author: Robin Aguilar
Beliveau and Noble Labs
University of Washington | Department of Genome Sciences
"""

##############################################################################

#specific script name
scriptName = "split_bed"

#import libraries
import time
import pandas as pd
import argparse

##############################################################################

def read_bed(bed_file):
    
    """
    Function takes the bed file and reads it into a pandas df.
    Parameters
    ----------
    bed_file : bed file
        file with region coords chr \t start \t stop

    Returns
    -------
    bed_df : dataframe
        dataframe of coordinates in file
    """
    
    colnames = ["chrom","start","stop"]
    
    bed_df = pd.read_csv(bed_file, names=colnames,header=None,
                            sep='\t')
    
    return bed_df
    
##############################################################################

def split_write_bed(bed_df,chrom,bed_out):
    """
    Function takes dataframe and subsets specific chrom into an output bed
    file with regions only corresponding to the chrom of choice.
    Parameters
    ----------
    bed_df : dataframe
        dataframe of region coordinates from file

    Returns
    -------
    None. Writes file with chromosome name as bed.
    """    
    
    #subsets rows containing specific chromosome
    chrom_df = bed_df[bed_df['chrom'].str.contains(str(chrom))]
    
    #writes row(s) to output bed file
    chrom_df.to_csv(bed_out,index = False,
                               index_label=False,
                               header = None,
                               sep = '\t')
    
    
##############################################################################

def main():
    
    start_time=time.time()

    """Reads a bed file provided by the user containing coordinates of
    regions for probe design. If regions on different chromosomes exist,
    this script will generate independent files for different regions based
    on chromosome."""
    
    #allow user inpir parameters on command line
    
    userInput = argparse.ArgumentParser(description=\
        '%Requires a bed file (tab sep, chrom \t start \t stop)'
        'And names of chromosomes to survey')
        
    requiredNamed = userInput.add_argument_group('required arguments')
    
    requiredNamed.add_argument('-b', '--bed_file', action='store', 
                               required=True, 
                               help='The region bed coordinates for probe'
                               'design')
    requiredNamed.add_argument('-c', '--chrom_name', action='store', 
                               required=True, 
                               help='The chrom to susbset from bed')
    requiredNamed.add_argument('-o', '--bed_out', action='store', 
                               required=True, 
                               help='path of output file')

    # Import user-specified command line values.
    args = userInput.parse_args()
    bed = args.bed_file
    chrom = args.chrom_name
    bed_out = args.bed_out
    
    
    bed_df = read_bed(bed)
    
    print("---%s seconds ---"%(time.time()-start_time))
    
    split_write_bed(bed_df,chrom,bed_out)
    
    print("---%s seconds ---"%(time.time()-start_time))
    

if __name__ == '__main__':
    main()
```
