## Supplementary Software for "Tigerfish designs oligonucleotide-based *in situ* hybridization probes targeting intervals of highly repetitive DNA at the scale of genomes": split_filter.py

```
#!/usr/bin/env python3
# -*- coding: utf-8 -*-
##############################################################################
"""
Created on Mon Jun 28 17:42:48 2021
@author: Robin Aguilar
Beliveau and Noble Labs
University of Washington | Department of Genome Sciences
"""
##############################################################################

#specific script name
script_name = "split_filter"

#load libraries used
import time
import argparse
import pandas as pd
from itertools import groupby
import os 

##############################################################################

def read_probe_file(file_path):
    """
    This function reads the probe file containing designed probes into a 
    pandas dataframe
    
    Parameters
    ----------
    file_path : file
        file containing probes that have been filtered
    Returns
    -------
    probe_df : dataframe
        returns a dataframe of probes that were filtered
    """
    
    #read in the file with the appropriate column names
    colnames = ["chrom","p_start","p_end","probe","Tm","region",
                "r_count_total","h_count_total","k_score","k_norm"]
    
    #read as a dataframe
    probe_df = pd.read_csv(file_path, delimiter = '\t', names = colnames)
    
    return probe_df

##############################################################################

def split_file(probe_df,out_path):
    """
    This function splits the dataframe into output files by repeat region as
    names.
    
    Parameters
    ----------
    probe_df : dataframe
        returns a dataframe of probes that were filtered
    out_path: file path
        file path of where split region files should be created.
    Returns
    -------
    None. Output files contain probes within each listed repeat region.
    """
    
    os.mkdir(out_path)

    #do a groupby over the repeat regions
    grouped_regions=probe_df.groupby('region',sort=False)
    
    #write each repeat region into a seperate file
    for name,group in grouped_regions:
        
        group.to_csv(str(out_path) + "/" + str(name) + ".txt", header=False,
                     index=False, sep="\t")

##############################################################################

def main():
    
    start_time=time.time()

    userInput = argparse.ArgumentParser(description=\
        '%Requires a probe file containing filtered probes'
        'This splits this file into independent repeat regions to prepare'
        'for parallel alignment as final filter pass')

    requiredNamed = userInput.add_argument_group('required arguments')
    requiredNamed.add_argument('-f', '--file_path', action='store', 
                               required=True, help='The post probe file'
                               'that contains all sequences and regions')
    requiredNamed.add_argument('-o', '--out_path', action='store',
                               required=True, help='The directory where'
                               'the split files will be located by repeat')
    
    args = userInput.parse_args()
    file_path = args.file_path
    out_path = args.out_path
    
    probe_df = read_probe_file(file_path)
    
    print("---%s seconds ---"%(time.time()-start_time))

    split_file(probe_df,out_path)
    
    print("---%s seconds ---"%(time.time()-start_time))
    
    print("done")
    
if __name__== "__main__":
    main()
```
