## Supplementary figures and images for "Tigerfish designs oligonucleotide-based *in situ* hybridization probes targeting intervals of highly repetitive DNA at the scale of genomes"

### bt2_repeat_disc.png

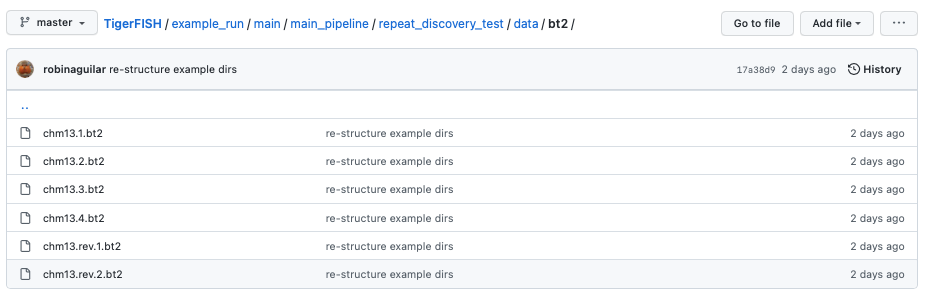

### check_repeat_disc.png

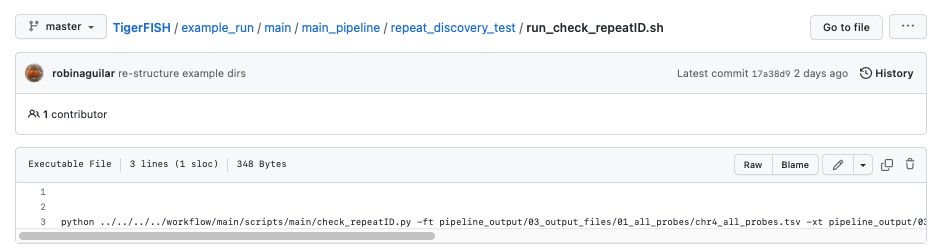

### chm13_chr9_yaml.png

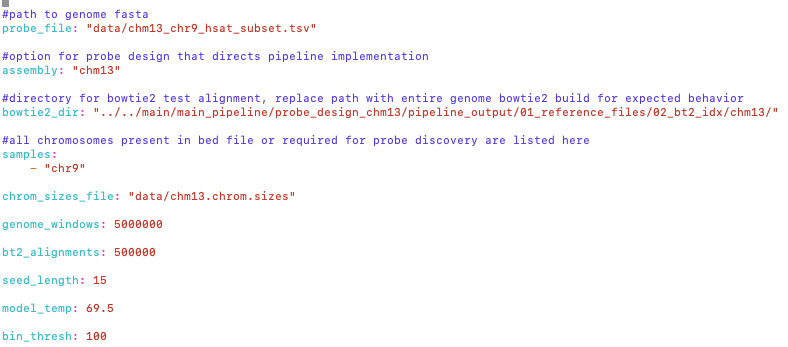

### chm13_probe_design_dir.png

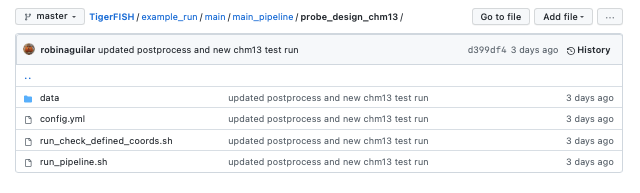

### chm13_yaml.png

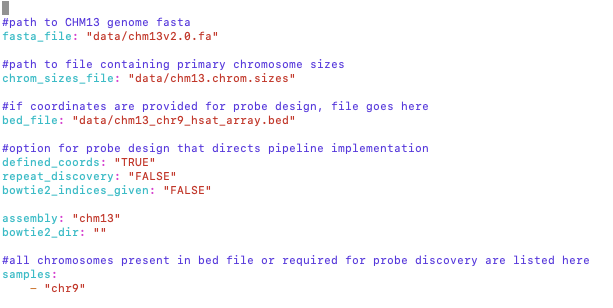

### chr4_genome_view.png

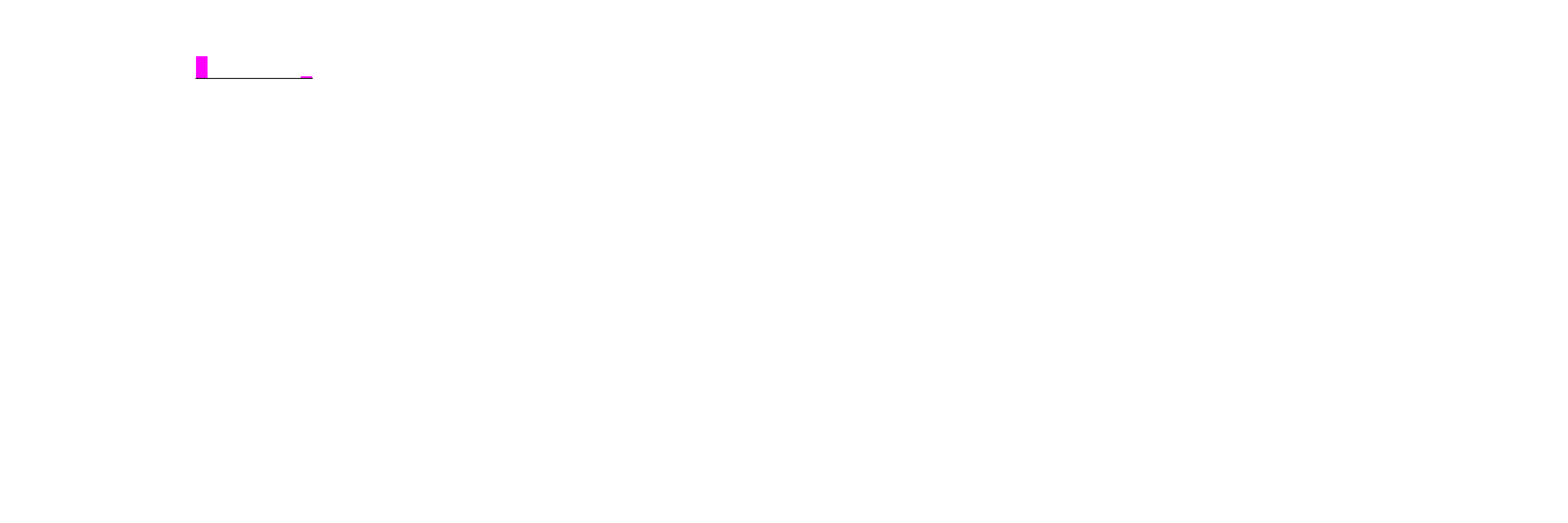

### chr9_genome_view.png

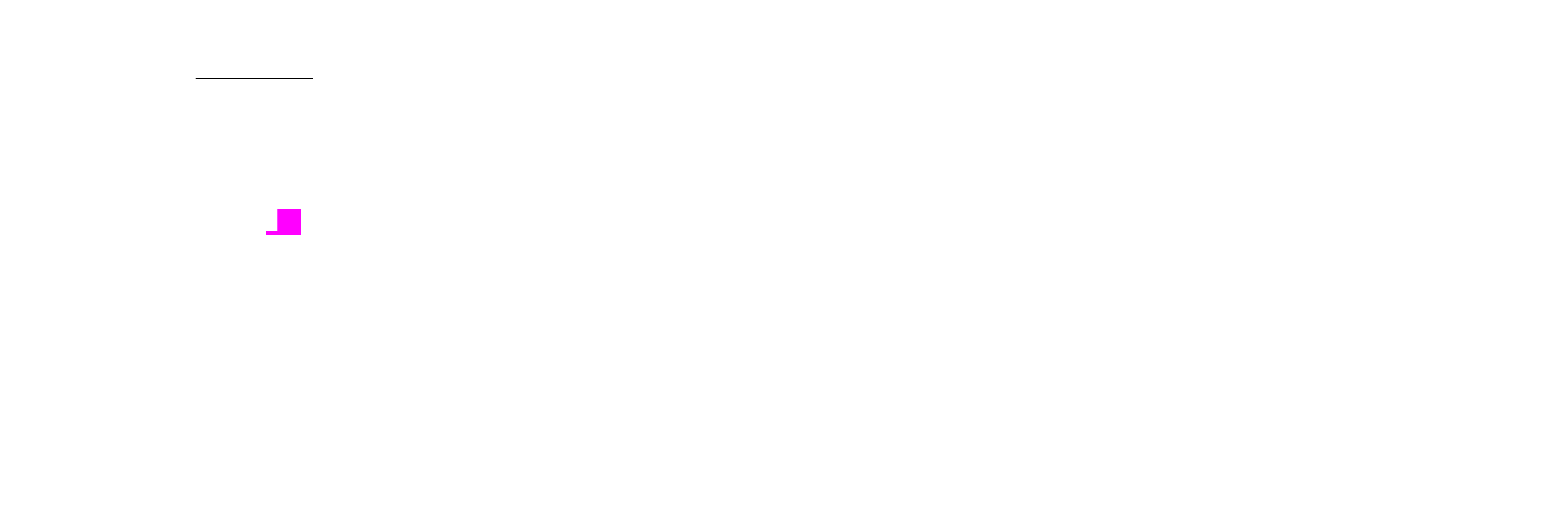

### chrom_sizes_repeat_disc.png

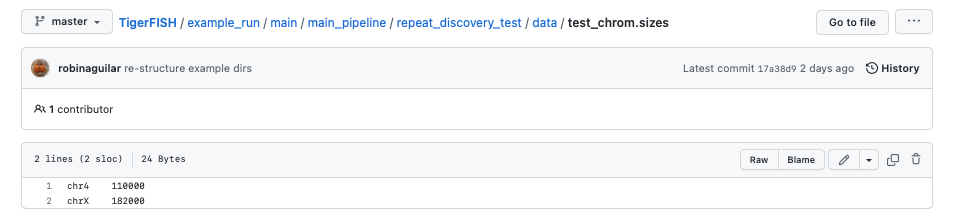

### chrX_genome_view.png

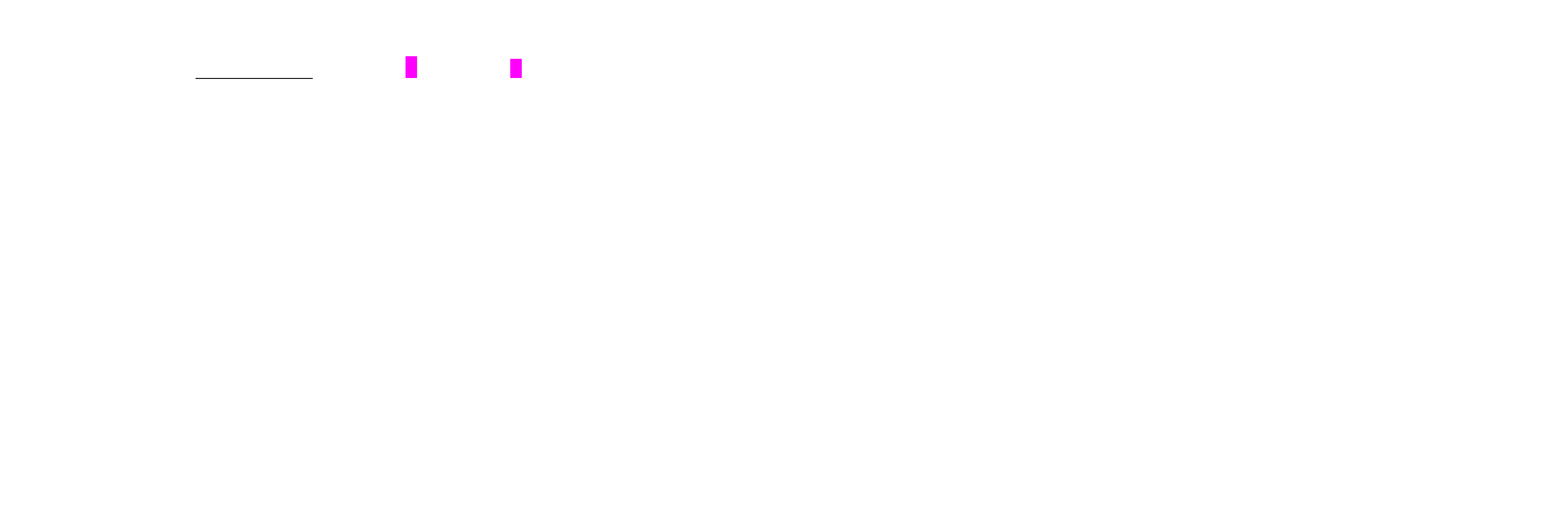

### DAG.pdf

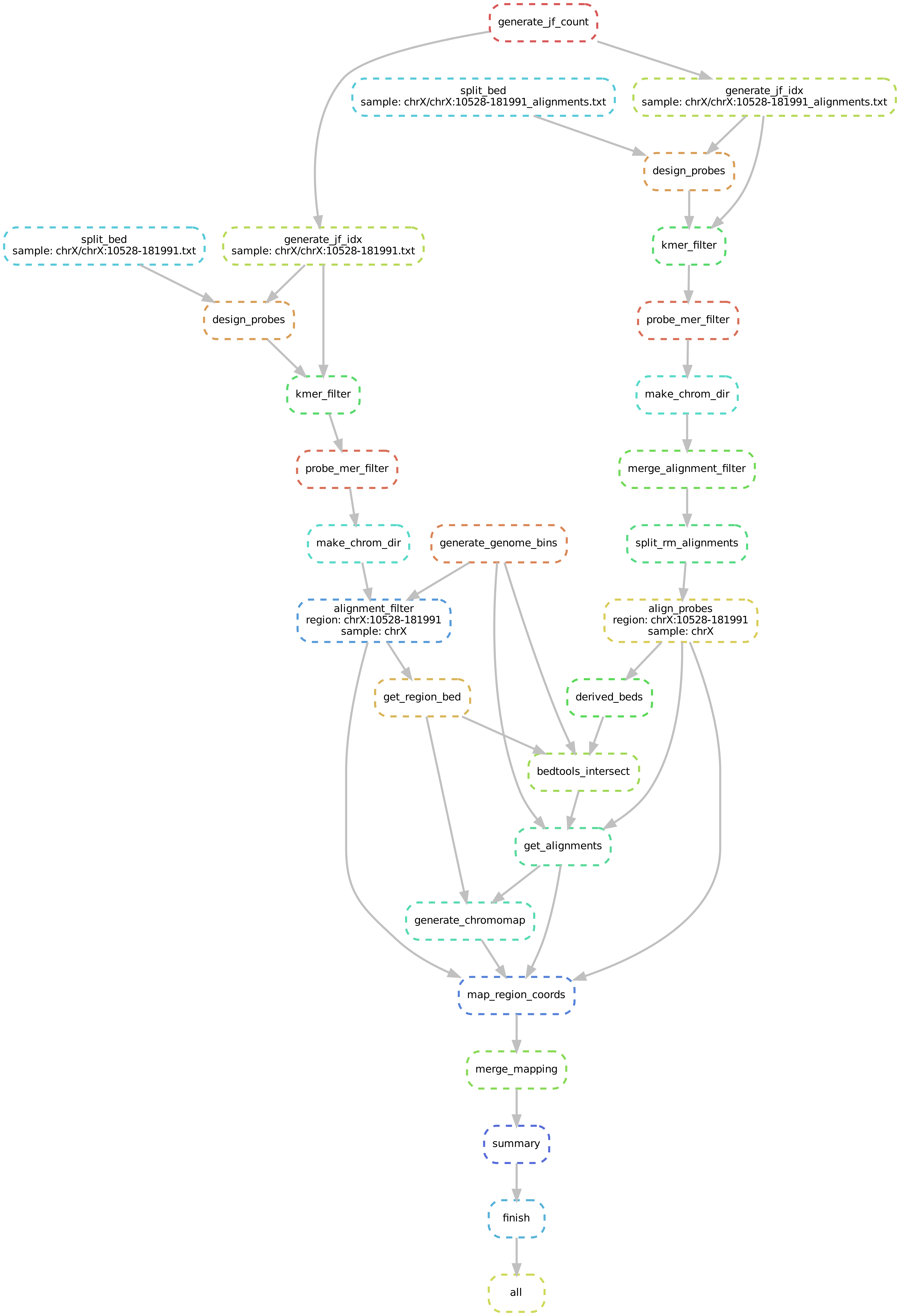

### genome_threshold_chr9.png

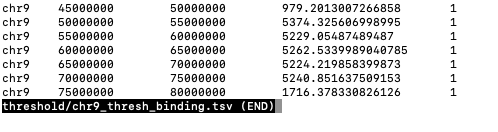

### genome_wide_binding_chr9.png

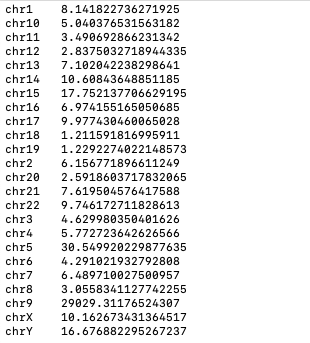

### pd_params.png

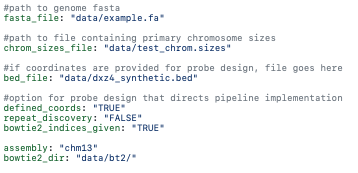

### pipeline.pdf

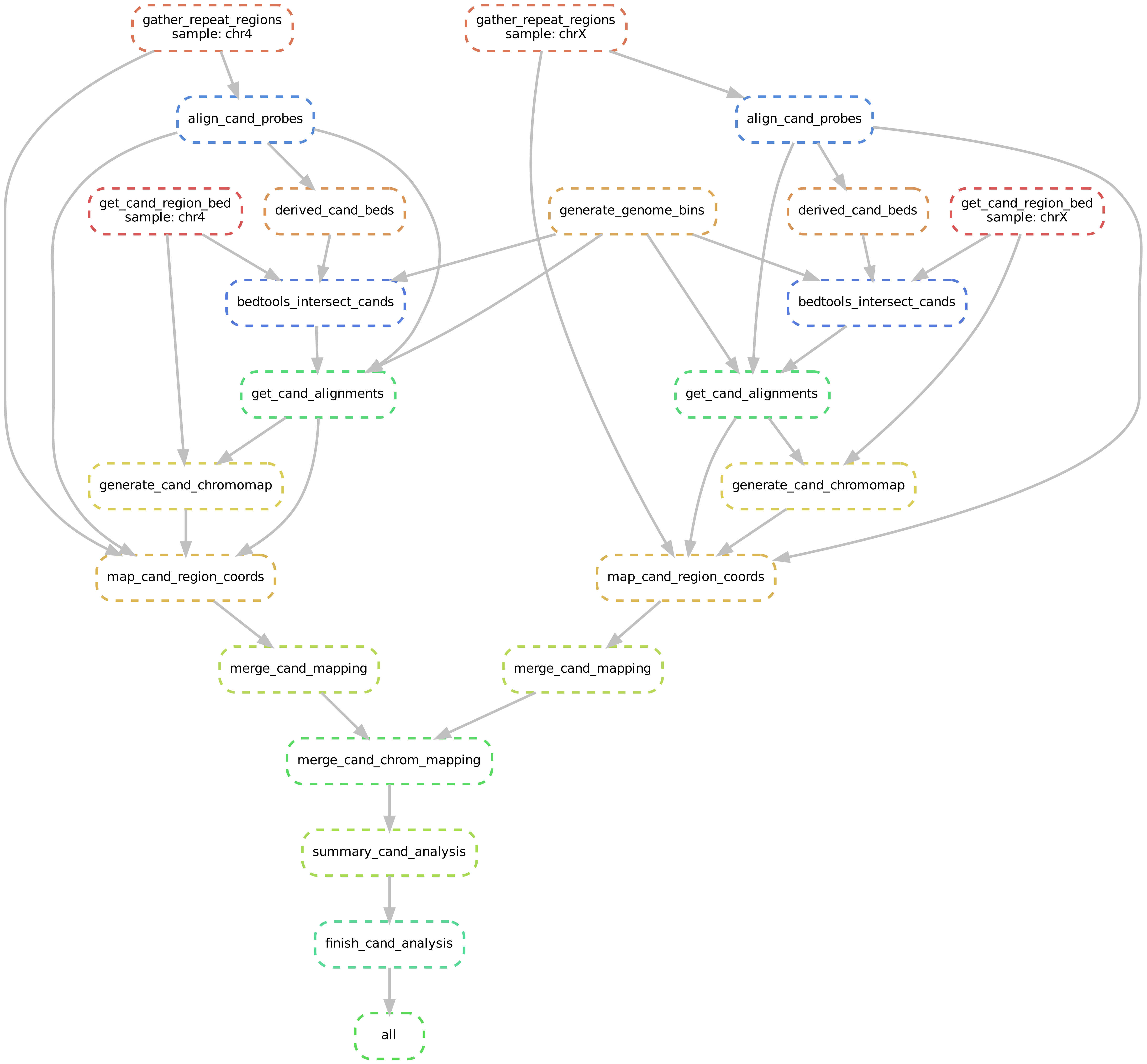

### pipeline.pdf

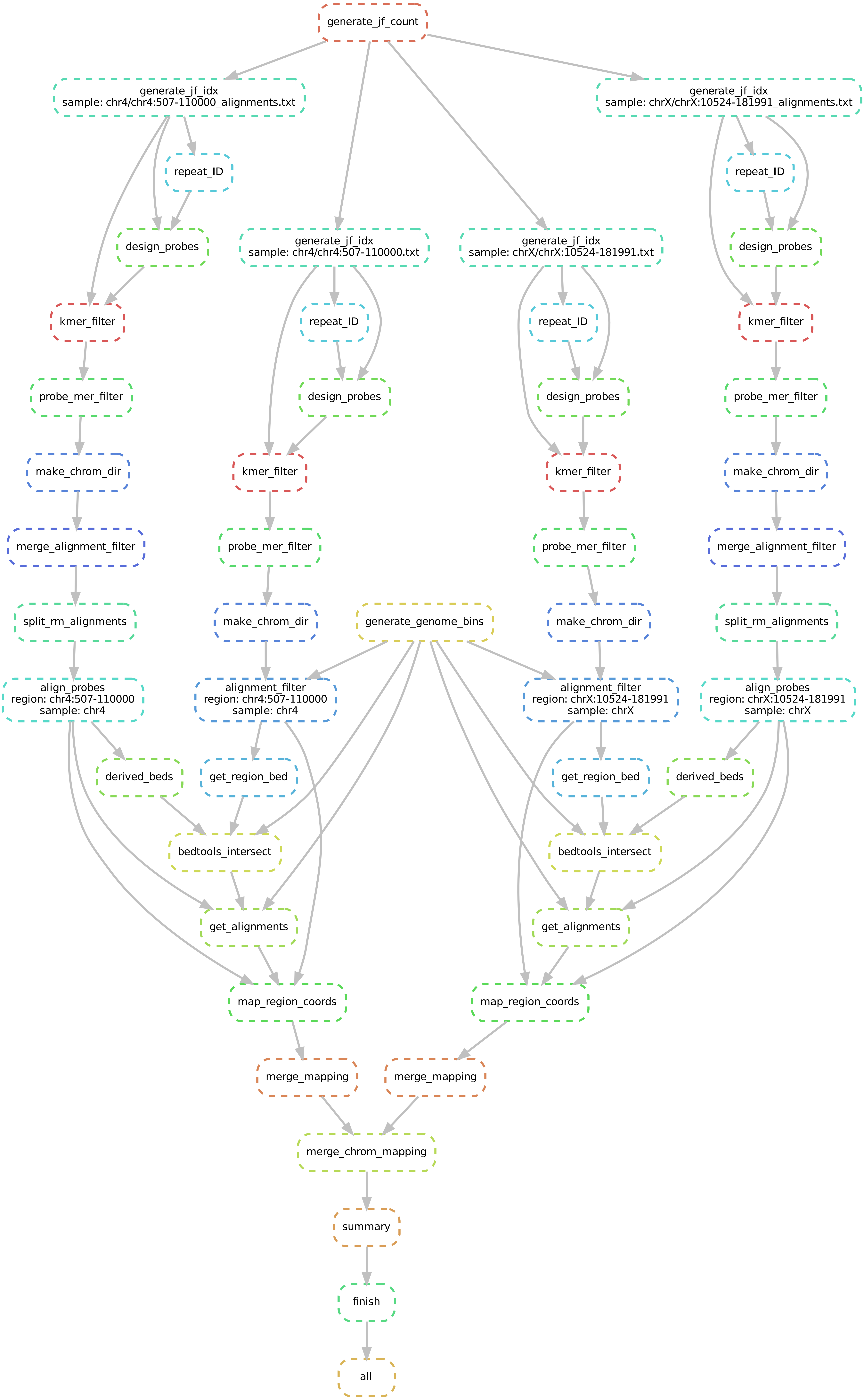

### pipeline.pdf

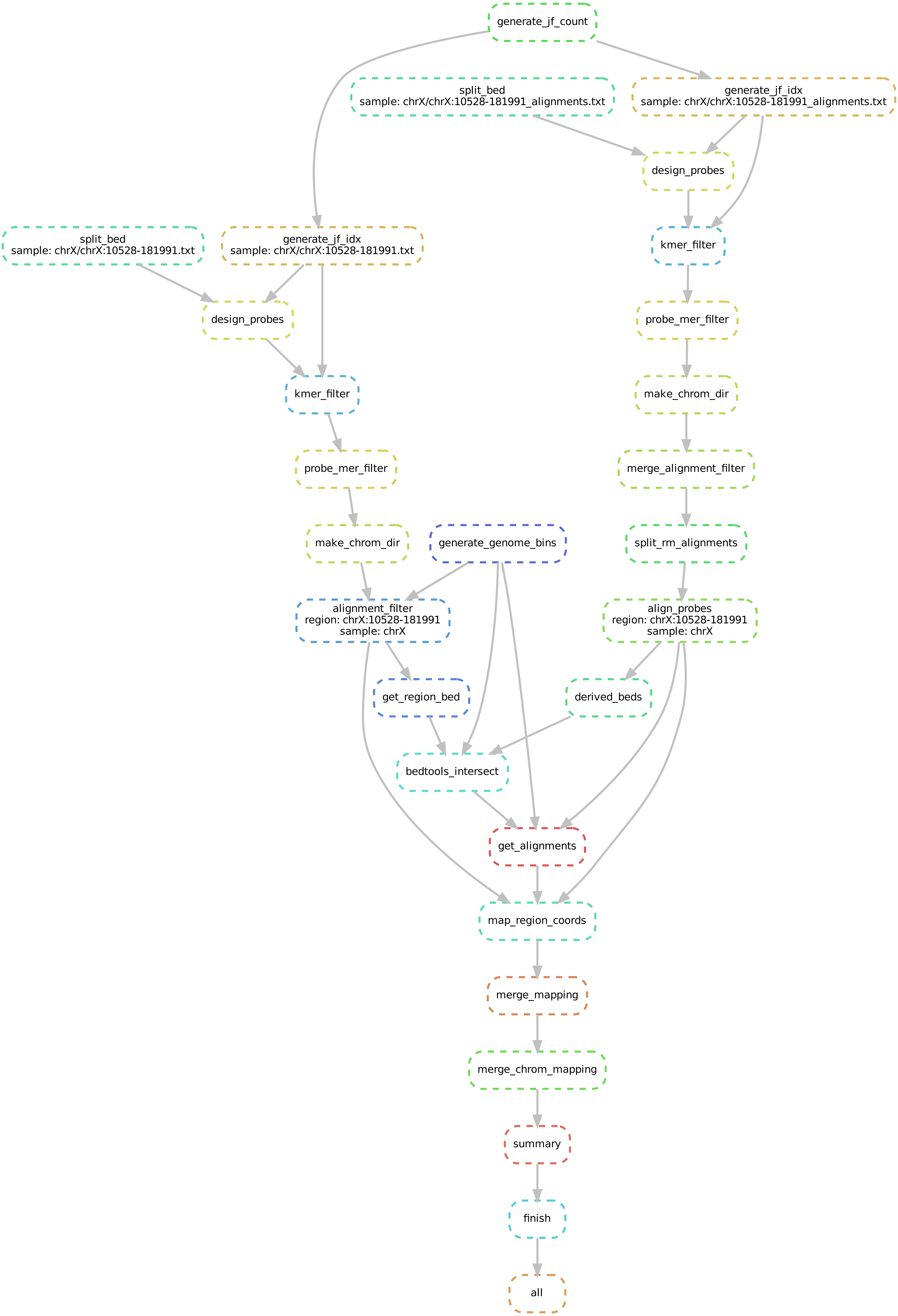

### postprocess_config.png

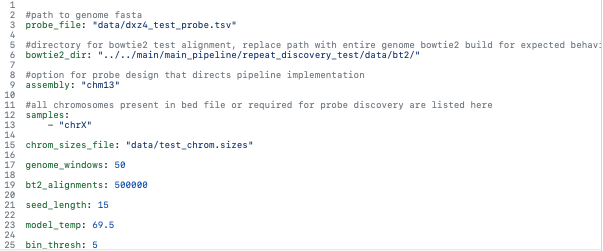

### postprocess_run.png

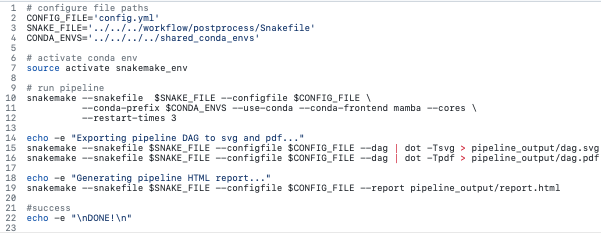

### postprocess_step_2.png

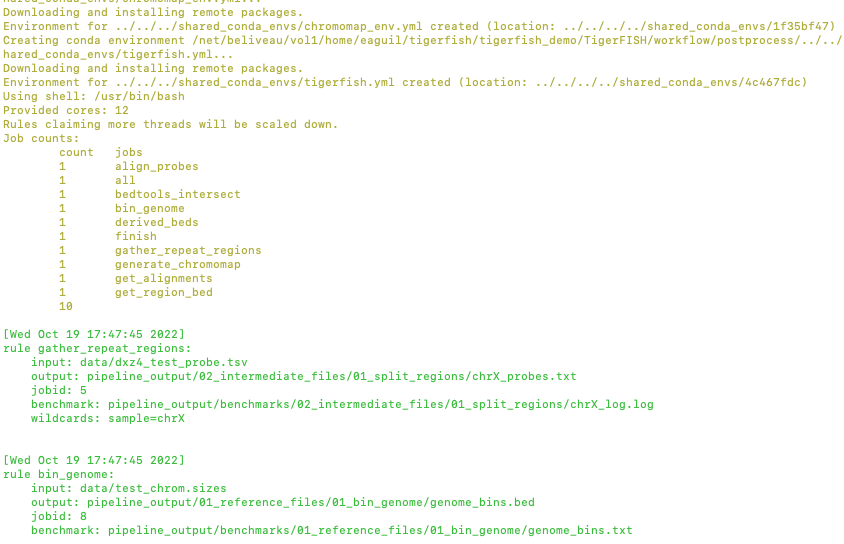

### postprocess_step_3.png

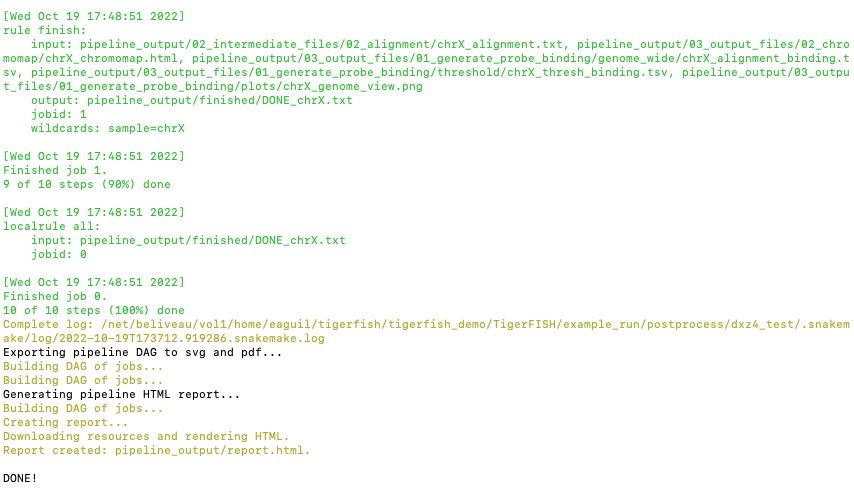

### probe_design_config.png

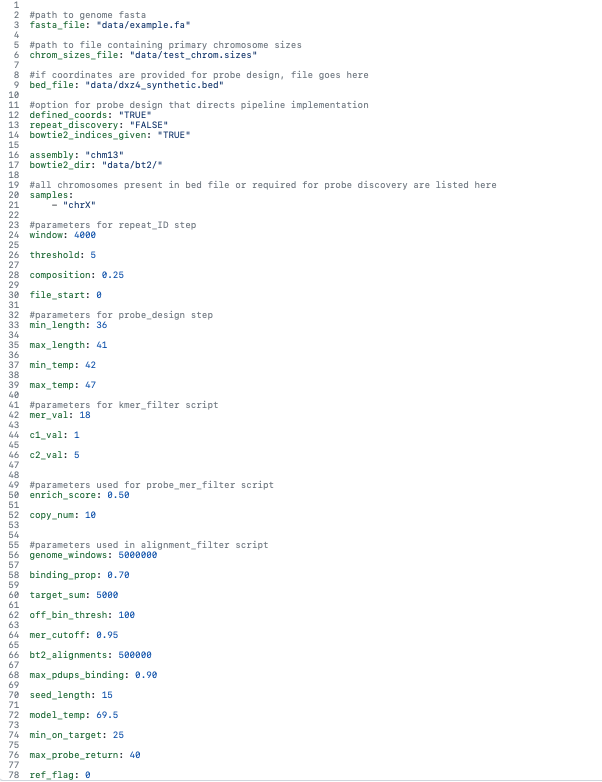

### rd_params.png

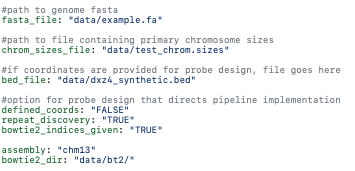

### repeat_disc_fasta.png

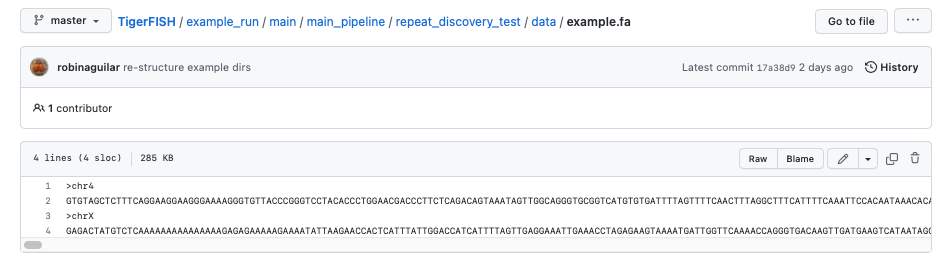

### repeat_discovery_config.png

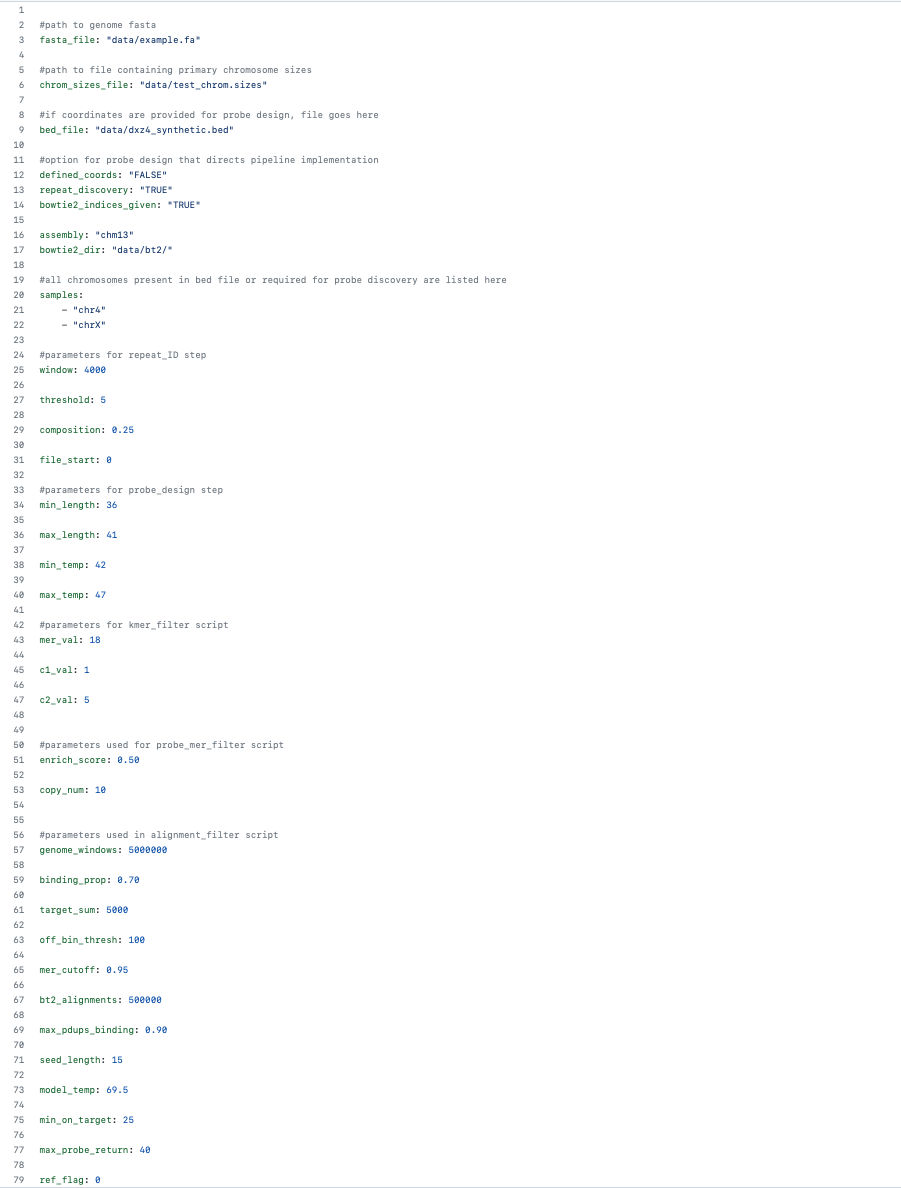

### run_pipeline_repeat_disc.png

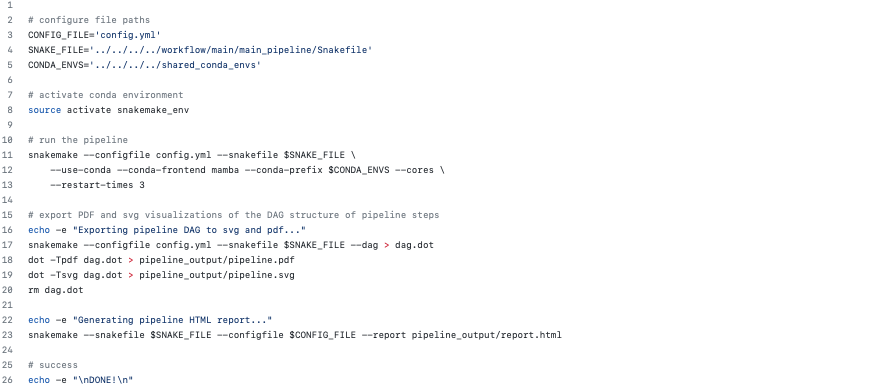

### sample_out_probe.png

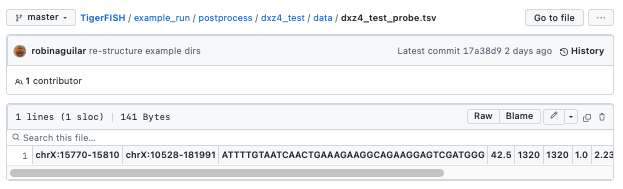

### step_1_probe_design.png

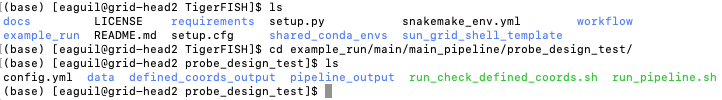

### step_2_probe_design.png

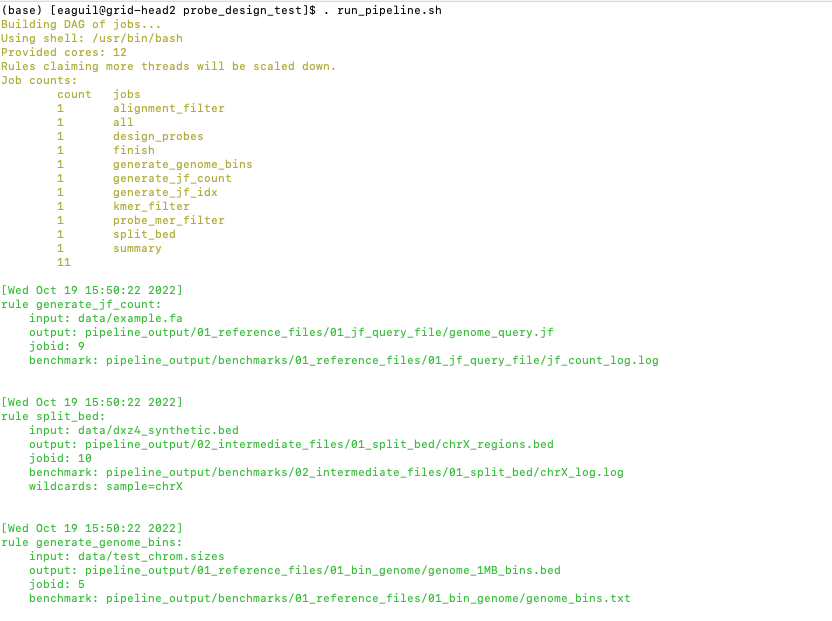

### step_2_repeat_disc.png

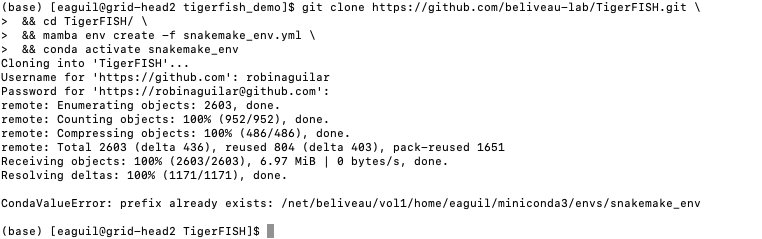

### step_3_probe_design.png

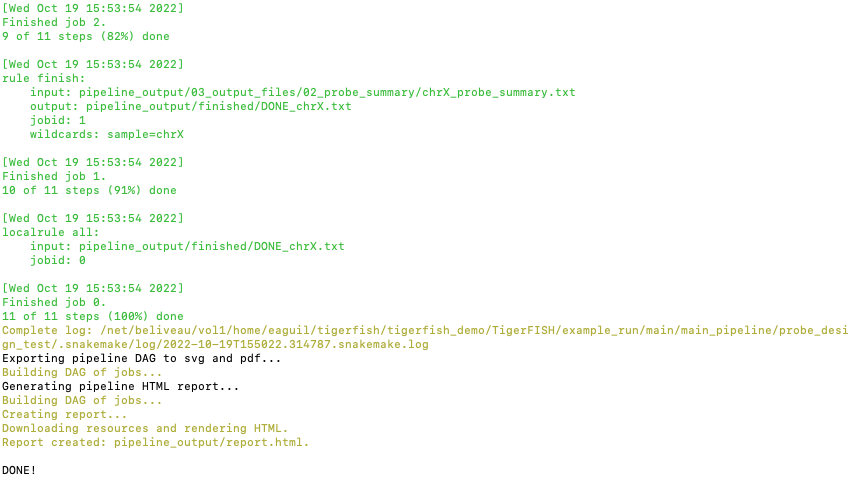

### step_4_probe_design.png

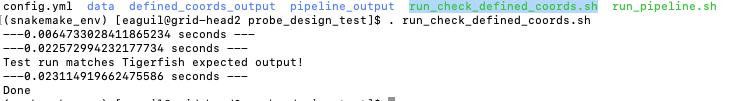

### step_4_repeat_disc.png

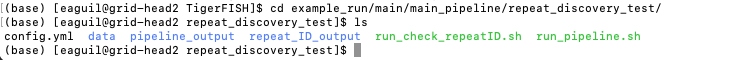

### step_5_repeat_disc.png

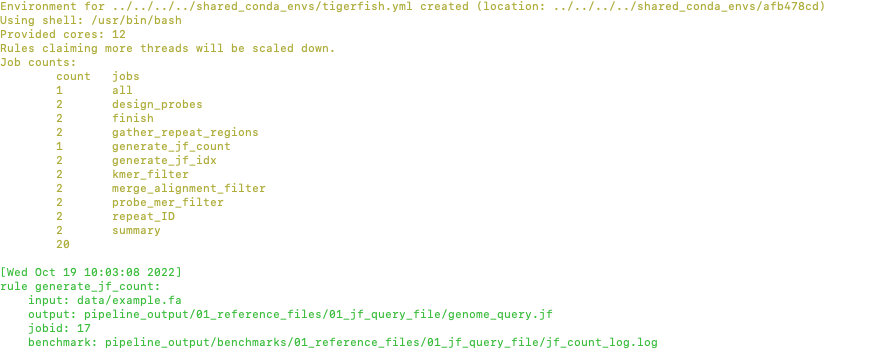

### step_6_repeat_disc.png

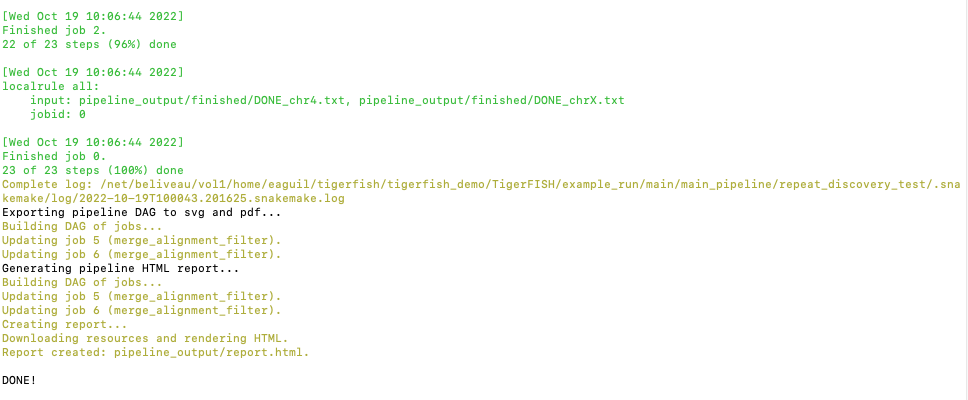

### step_7_repeat_disc.png

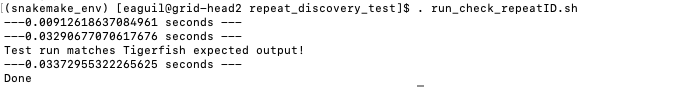

### tiger.jpg

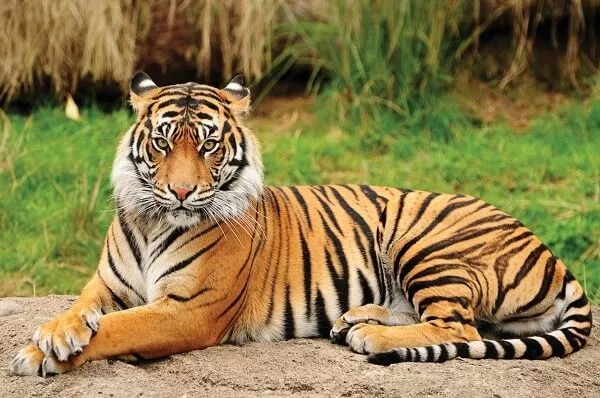

### tigerfish_fish.jpg

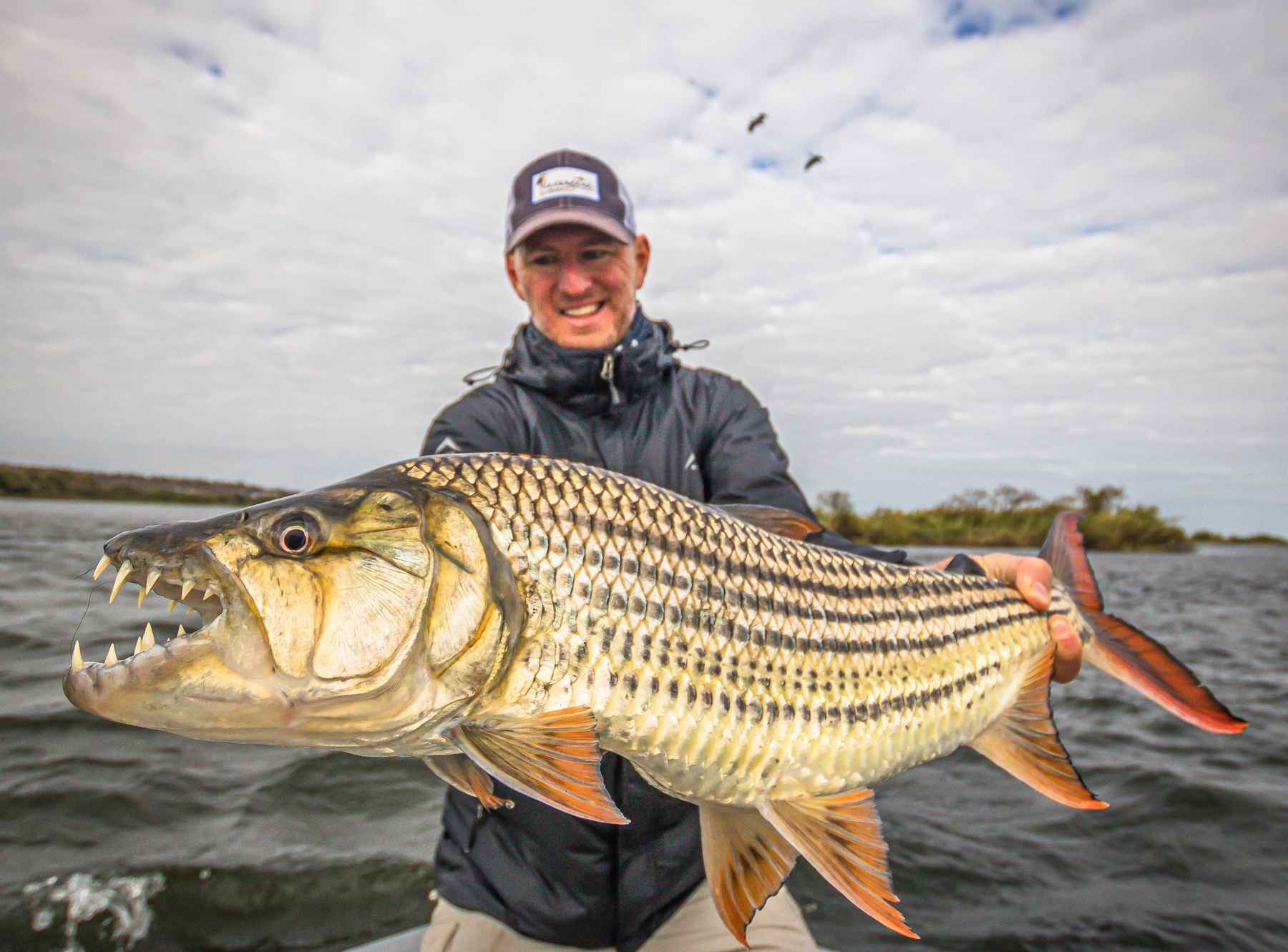

### tigerfish_logo.png

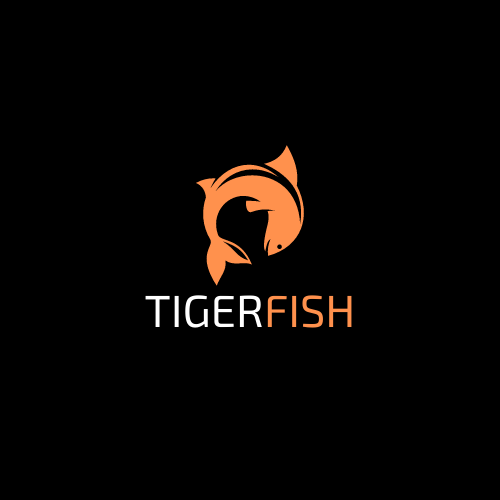

### tigerfish_main_overview.png

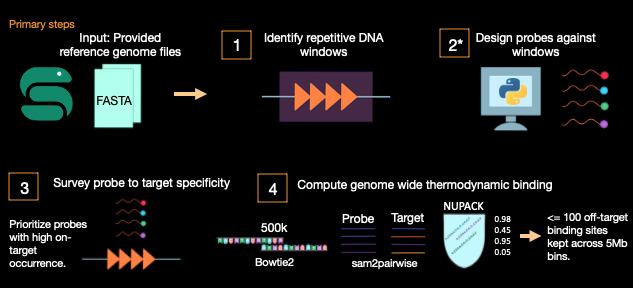

### tigerfish_postprocess_overview.png

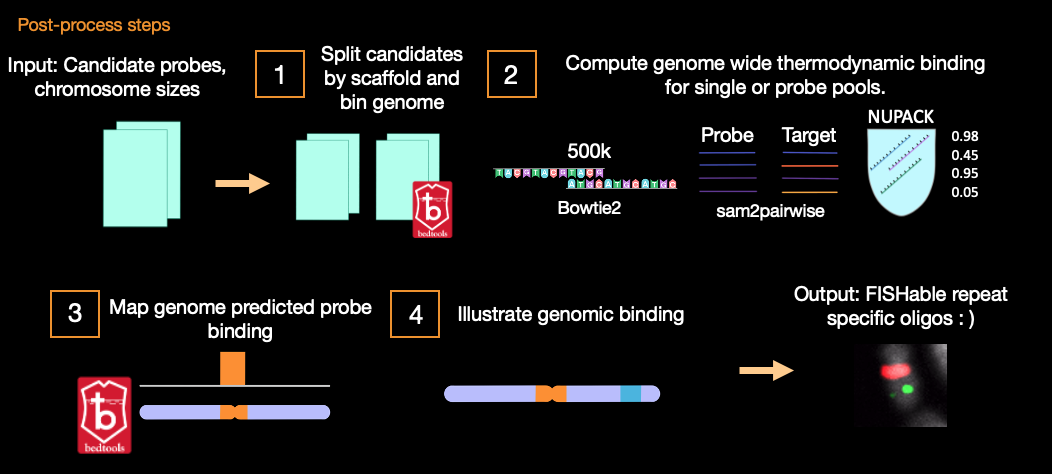

### tutorials_summary.png

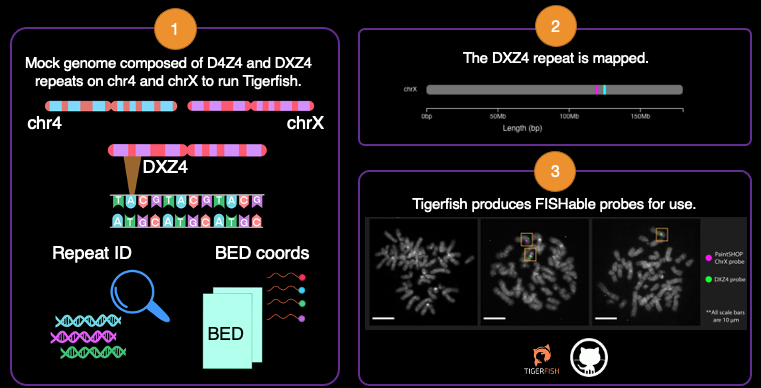
